# Bayesian Least-Squares Supertrees (BLeSS): Flexible Inference of Large Time-Calibrated Phylogenies

**DOI:** 10.1101/2024.11.29.625936

**Authors:** David Černý, Graham J. Slater

## Abstract

Time-calibrated phylogenies are key to macroevolutionary hypothesis testing and parameter inference, but their estimation is difficult when the number of tips is large. Despite its attractive properties, the joint Bayesian inference of topology and divergence times remains computationally prohibitive for large supermatrices. Historically, supertrees represented a popular alternative to supermatrix-based phylogenetic methods, but most of the existing supertree techniques do not accommodate branch lengths or topological uncertainty, rendering them unfit to supply input for modern comparative methods. Here, we present Bayesian Least-Squares Supertrees (BLeSS), a new approach that takes a profile of time trees with partially overlapping leaf sets as its input, and returns the joint posterior distribution of supertree topologies and divergence times as its output. Building upon the earlier exponential error model and average consensus techniques, BLeSS transforms the profile into path-length distance matrices, computes their arithmetic average, and uses MCMC to sample time-calibrated supertrees according to their least-squares fit to the average distance matrix. We provide a fast, flexible, and validated implementation of BLeSS in the program `RevBayes`, and test its performance using a comprehensive set of simulations. We show that the method performs well across a wide range of conditions, including variation in missing data treatment and the steepness of the error function. Finally, we apply BLeSS to an empirical dataset comprising 33 time trees for 260 species of carnivorans, illustrating its ability to recover well-supported clades and plausible node ages, and discuss how the method can best be used in practice, outlining possible extensions and performance boosts.

## Introduction

Time trees, or phylogenetic trees with branch lengths scaled to calendar time, are widely used to investigate questions ranging from trait evolution (Ree, 2005; Slater and Harmon, 2013; Hopkins and Smith, 2015; Slater, 2015; Caetano and Harmon, 2018) to diversification rate estimation (Nee et al., 1994; Rabosky, 2014; Maliet et al., 2019; Barido-Sottani et al., 2020) and historical biogeography (Ree and Smith, 2008; Landis et al., 2013; Meseguer et al., 2014). Over the last two decades, joint Bayesian inference of tree topology and divergence times (Drummond et al., 2006; Ronquist et al., 2012) has become the preferred method for producing time trees thanks to its ability to synthesize multiple types of evidence and accommodate the different sources of uncertainty that are associated with them (dos Reis et al., 2016). Unfortunately, due to the intrinsic difficulty of sampling from complex posterior distributions, Bayesian co-estimation of topology and node ages remains computationally unfeasible for datasets comprising thousands or tens of thousands of tips, which are increasingly used to shed light on the patterns and processes underlying major radiations across the tree of life (Jetz et al., 2012; Rabosky et al., 2018; Upham et al., 2019; Janssens et al., 2020).

Two alternatives have emerged to the intractable fully Bayesian inference of very large time trees. One is the assembly of a comprehensive molecular dataset, or “supermatrix” (Delsuc et al., 2005; de Queiroz and Gatesy, 2007), followed by a time-free analysis employing a less computationally demanding method, such as a heuristic maximum likelihood search. The resulting phylogram is then time-scaled using nonparametric or semiparametric rate smoothing (Rabosky et al., 2018; Janssens et al., 2020). The other approach involves dividing the clade of interest into a number of non-overlapping subclades, each of which is sufficiently small for fully Bayesian divergence time estimation to remain feasible. The full-sized tree is then assembled by attaching the taxonomically comprehensive subclade phylogenies to a sparsely sampled backbone in which every subclade is represented by a single pair of tips. A pseudoposterior distribution of topologies and divergence times can be obtained by repeating the attachment step on multiple samples from the subclade and backbone posteriors (Jetz et al., 2012; Tonini et al., 2016; Upham et al., 2019). Though occasionally contrasted with more traditional “supertree” methods (Upham et al., 2019, 2021), this “backbone-and-patch” approach can in fact be viewed as their special case, since the final tree is in both cases assembled from previously inferred source trees rather than estimated directly from character data (Sanderson et al., 1998; Bininda-Emonds et al., 2002; Wilkinson et al., 2005).

Despite their widespread use, both the *post hoc* time-scaling of phylograms and the backbone-and-patch approach present important shortcomings. The former method lacks some of the key properties of fully Bayesian time tree estimation, including the estimation of divergence times jointly with (rather than conditional on) tree topology, and the ability to accommodate various sources of uncertainty, such as variation in branch length estimates and fossil calibration error (Smith and O’Meara, 2012). The latter framework retains the advantages of fully Bayesian inference at the level of source trees, but fails to propagate uncertainty throughout the tree assembly process as a result of the limited overlap between the backbone and the patch trees. Moreover, the delimitation of patch clades requires potentially controversial monophyly assumptions (Lloyd and Slater, 2021), and the pseudoposterior produced by the method is not a statistically valid approximation of the full posterior, since the likelihoods of the backbone and patch trees are treated as independent despite being partly derived from the same set of sequences (Álvarez-Carretero et al., 2022).

The shortcomings of the backbone-and-patch framework can be overcome in a general supertree setting; however, existing supertree methods – including the widely used matrix representation approaches (Baum and Ragan, 2004; Ross and Rodrigo, 2004; Nguyen et al., 2012) – suffer from important drawbacks of their own. First, their ability to compete with alternative approaches to the inference of very large phylogenies is hampered by a lack of conceptual clarity on whether supertrees should be viewed merely as summaries of the information contained in the source trees, or estimates in their own right (Steel and Rodrigo, 2008). This has implications for whether supertree methods should attempt to resolve topological conflict, or merely represent it as a lack of resolution (the “voting” vs. “veto” approaches; Ranwez et al., 2007), and whether it is acceptable for the supertree to contain bipartitions not present in any of its source trees (Cotton et al., 2006; Wilkinson et al., 2007). Second, most supertrees do not come with branch lengths, which makes them of limited use as input for phylogenetic comparative methods. Branch lengths in units of expected substitutions per site or calendar time can be added to a supertree *post hoc* using sequence data (Binet et al., 2016; Kimball et al., 2019) or stratigraphic age ranges associated with the tips (Sakamoto et al., 2016), but few supertree methods are capable of estimating topology and branch lengths simultaneously, without relying on an increasingly tenuous chaining together of inferences (though see Willson, 2004; Lapointe and Levasseur, 2004; and Criscuolo et al., 2006 for exceptions). Third, most existing supertree methods yield point estimates without any measure of uncertainty. While extensions of the nonparametric bootstrap to the supertree setting have been proposed (Burleigh et al., 2006), they suffer from a variety of problems, including the impossibility of ensuring that boostrap pseudoreplicates will have the same leaf set as the original supertree (Bininda-Emonds, 2014). Bayesian posterior probabilities represent an attractive, easily interpretable alternative to bootstrap support values (Alfaro and Holder, 2006), but their use in supertree estimation remains uncommon (Ronquist et al., 2004; Akanni et al., 2015; Karcher et al., 2021).

Here, we present a new approach to supertree inference, the Bayesian Least-Squares Supertrees (BLeSS), which takes a profile of time trees with overlapping leaf sets as its input, and returns a posterior distribution of time-calibrated supertrees as its output. BLeSS can be used in a manner similar to other supertree methods, with each dataset providing a single source tree representing a point estimate of topology and divergence times. However, the framework can also accommodate uncertainty about source tree topology by including the entire posterior distribution estimated from a given dataset in the profile, as proposed by Ronquist et al. (2004) and similar to the “metatree” pipeline of Lloyd and Slater (2021). Unlike the backbone-and-patch approach, it is capable of efficiently propagating this uncertainty into the posterior supertree distribution by allowing for an arbitrary degree of overlap between the source trees. Re-use of the same data by multiple source trees is allowed but not required, and can be addressed in a nonparametric fashion by means of optional downweighting of non-independent source trees, without treating each one as providing an independent contribution to the likelihood term. Moreover, the use of a birth-death tree prior allows BLeSS to incorporate other sources of information in addition to the profile, such as node calibrations or topological constraints. We validate our implementation of BLeSS using simulation-based calibration checking, employ a comprehensive simulation scheme to assess its accuracy and precision, benchmark its performance across a wide range of supertree sizes, and finally demonstrate its utility by inferring a time-calibrated supertree for extant members of the mammalian order Carnivora.

## Materials and Methods

### Theory

The BLeSS method combines elements from the average consensus (Lapointe and Levasseur, 2004) and maximum likelihood (Steel and Rodrigo, 2008) approaches to supertree inference, and consists of two main components: an average distance matrix summarizing the topological and divergence time information contained in a profile of source trees, and an exponential error model used to evaluate the likelihood of a proposed supertree conditional on this matrix. Like any probabilistic model, the approach outlined here can be used in both maximum-likelihood and Bayesian settings. Here, we choose to focus on the Bayesian implementation, which provides a natural way of quantifying the uncertainty associated with parameter estimates in the form of the posterior distribution. In addition, previous attempts to use the exponential error model for supertree inference revealed that the sampling techniques used for Bayesian inference were more efficient than the optimization techniques employed by maximum likelihood, making the former approach better suited for analyses of large datasets (Akanni et al., 2014, 2015).

#### Average distance matrix

Consider a profile of source trees 𝒫 = {𝒯_1_*, …,* 𝒯*_K_*} defined over a set of leaves ℒ, where ℒ(𝒯*_k_*) is the leaf set of tree 𝒯*_k_* such that ℒ(𝒯*_k_*) ⊆ ℒ. Each tree is assumed to be ultrametric, with branch lengths in units of calendar time. We exploit the fact that each tree 𝒯*_k_* can be converted to a unique path-length distance matrix **D***_k_* (Lapointe and Levasseur, 2004), such that the path-length distance *d_k_*(*i, j*) of any two leaves *i* and *j* is computed by taking the sum of the lengths of the intervening branches, and corresponds to twice the time elapsed since their divergence. If we denote by *π_i,j_*(𝒫) the (possibly empty) set of indices of all trees in 𝒫 that contain both *i* and *j*, that is, *π_i,j_*(𝒫) = {*k* ∈ {1*, …, K*} : *i* ∈ ℒ(𝒯*_k_*) ∧ *j* ∈ ℒ(𝒯*_k_*)}, the *average distance matrix* **D̅** can be calculated element-wise as follows:

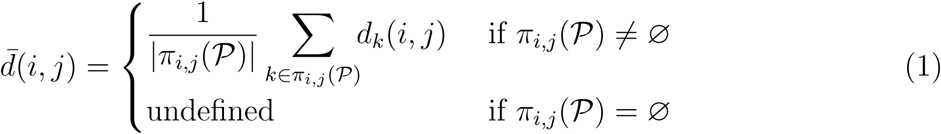

The undefined entries in the resulting matrix, corresponding to missing distances, can either be treated as such and disregarded in subsequent steps (the “direct” approach of Levasseur et al., 2003; see Makarenkov and Leclerc, 1999; Kettleborough et al., 2015), or they can be filled in with estimated distances using a variety of imputation schemes (the “indirect” approach; see Landry and Lapointe, 1997 and the references in Criscuolo and Gascuel, 2008).

Compared to related methods (Lapointe and Levasseur, 2004; Criscuolo et al., 2006), several simplifying assumptions are made here. The distances *d_k_*(*i, j*) are obtained by transforming trees rather than estimated from sequence data, and as such are guaranteed to be ultrametric. Branch length heterogeneity was found to negatively impact the accuracy of supertree inference by Lapointe and Levasseur (2004), leading to attempts to “standardize” (Lapointe and Levasseur, 2004) or “deform” (Criscuolo et al., 2006) individual distance matrices by using a linear combination of the original distances to minimize the among-matrix least-squares difference. This step is unnecessary here, since all source trees are assumed to have branch lengths measured in time units (e.g., millions of years), and as long as these units are consistent, calendar time itself provides a natural common scale for all path-length matrices.

#### Exponential error model

Once **D̅** has been calculated, the goal is to find a supertree 𝒯̂ with the full leaf set ℒ that minimizes the least-squares difference between its own path-length matrix **D̂** and **D̅** :

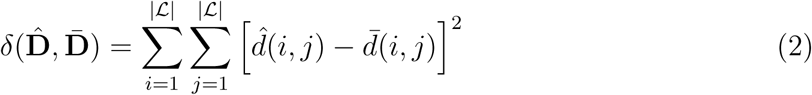

However, the problem of minimizing *δ*(**D̂**, **D̅**) is NP-hard (Day, 1987; Makarenkov and Lapointe, 2004). Consequently, attempts to estimate supertrees from the average distance matrix, or more generally, to infer phylogenies from distance matrices, have generally relied on heuristic optimization to obtain a point estimate of 𝒯̂ (Makarenkov and Leclerc, 1999; Lapointe and Levasseur, 2004). In contrast to these earlier approaches, BLeSS recasts the problem as one of sampling the posterior distribution of 𝒯̂, and accordingly requires an explicit error model to estimate the likelihood of a given supertree given the average distance matrix. To this end, Steel and Rodrigo (2008) proposed the following exponential error model:

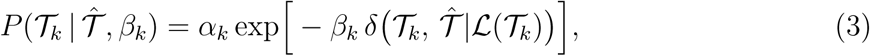

where 𝒯̂|ℒ(𝒯*_k_*) is the restriction of 𝒯̂ to the leaf set of source tree 𝒯*_k_*, and *β_k_* is a tree-specific positive constant that can be assigned to each source tree to reflect data quality, or inferred using maximum likelihood. This model is not immediately applicable to BLeSS, since it evaluates the likelihood of 𝒯̂ conditional on the untransformed tree profile 𝒫 rather than the average distance matrix **D̅**. Moreover, since the model was originally developed to estimate supertree topologies rather than supertrees with branch lengths, *δ* was assumed to be a discretely varying metric (such as the Robinson-Foulds distance; Robinson and Foulds, 1981), and *P* (·) was assumed to be a discrete probability distribution. This requires the inclusion of a normalizing term *α_k_*, which proved to be dependent on the shape of 𝒯̂ (Bryant and Steel, 2009). Evaluating *α_k_* is extremely computationally expensive, and while it is possible to choose the *β_k_* coefficients in such a way that the ranking of possible supertrees in terms of their likelihood remains unaffected by *α_k_*, this property is only sufficient for maximum likelihood inference (Bryant and Steel, 2009). However, previous attempts to perform maximum likelihood estimation under the model of Steel and Rodrigo (2008) proved incapable of efficient tree searches for large values of |ℒ| (Akanni et al., 2014), whereas the Bayesian MCMC implementation of Akanni et al. (2015) performed better but could no longer be guaranteed to be valid given its reliance on inaccurately normalized likelihoods (Bryant and Steel, 2009).

Here, we overcome the normalization problem by replacing the discrete distribution describing the probability of observing tree 𝒯*_k_* with a density function describing the probability of observing an average distance matrix **D̅**, using a continuously varying metric *δ* corresponding to Eq. 2:

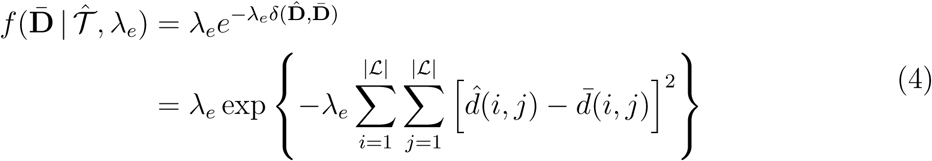

The rate parameter *λ_e_* of the exponential determines how strongly trees with larger values of *δ* will be penalized. Conveniently, actual supertree evaluation is likely to take place in terms of log-likelihoods, which are simply proportional (up to a multiplicative and additive constant) to the raw least-squares difference between the average distance matrix and the path-length distance matrix of the proposed supertree:

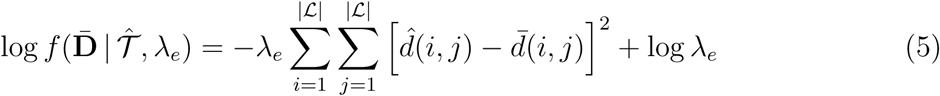

Using the likelihood function above, Bayesian inference becomes possible if an appropriate prior is specified. Since the estimated supertree 𝒯̂ is itself a time tree, any joint prior distribution over topologies and node heights can be used for this purpose, including the phenomenological “uniform” prior of Ronquist et al. (2012) or priors induced by the birth-death family of models (Gernhard, 2008). A realistic case of BLeSS supertree estimation under a birth-death prior with incomplete sampling and a uniform hyperprior on the time of origin is shown in Fig. 1. If the parameters of the birth-death process are jointly denoted as **Φ** and assumed to be estimated from the data along with the supertree, the joint posterior distribution of (𝒯, **Φ**) is given by

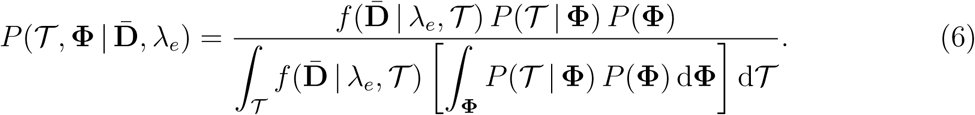

**Fig. 1.**
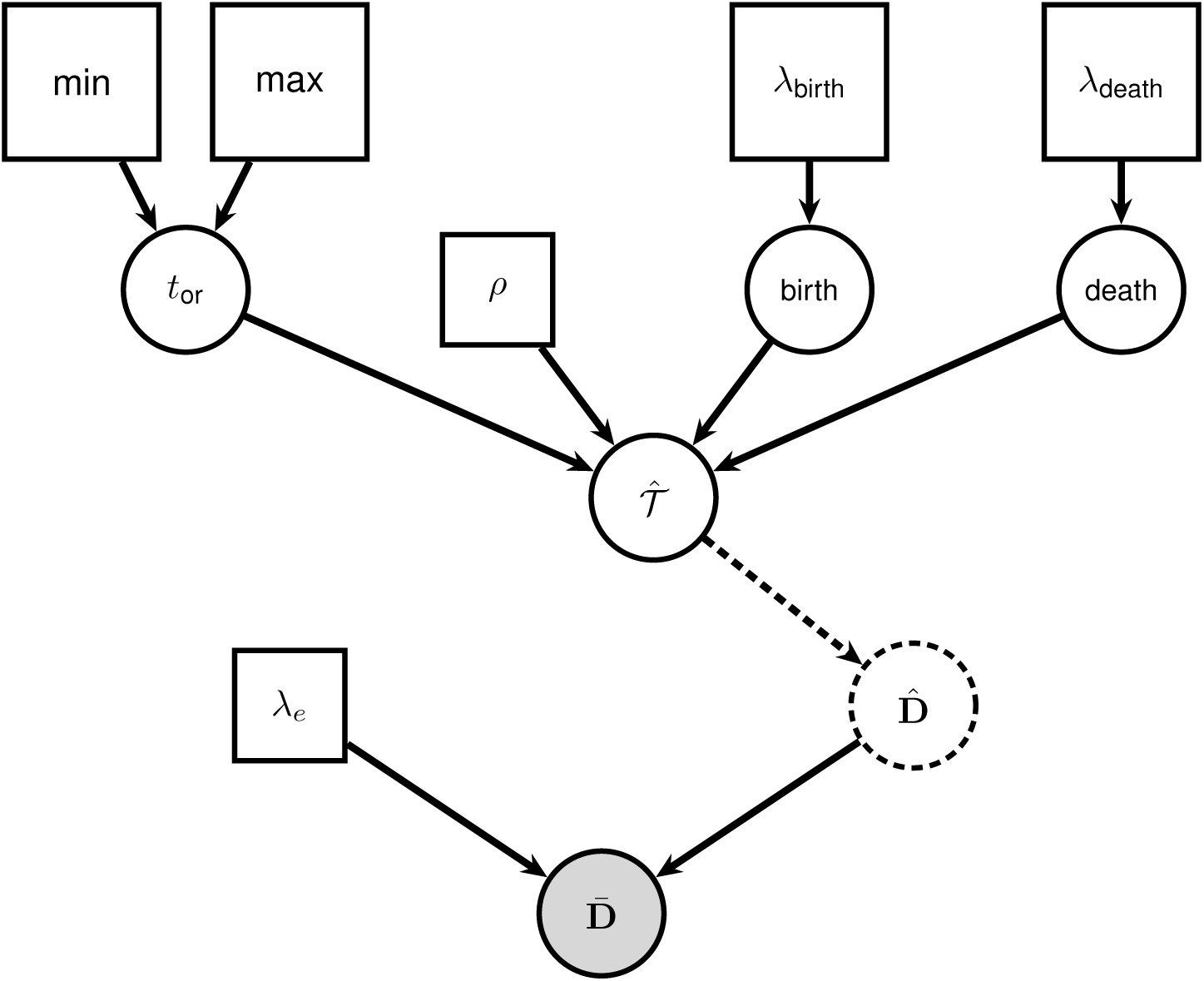
Probabilistic graphical model representation of a hypothetical BLeSS analysis. Notation after Höhna et al. (2014). *t*_or_ = time of origin of the birth-death process; *ρ* = extant sampling fraction; *λ*_birth_ = rate of the exponential hyperprior on the birth rate; *λ*_death_ = rate of the exponential hyperprior on the death rate.

Alternatively, instead of setting *λ_e_* to a fixed value as in Fig. 1, one can also assign it a prior distribution and use MCMC sampling to marginalize it out of the model:

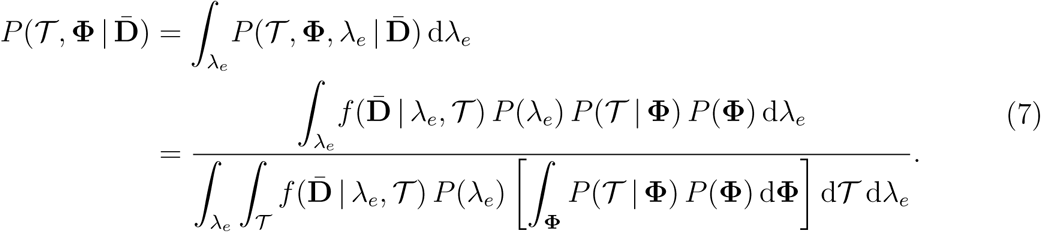

### Implementation

We implemented BLeSS in `RevBayes` (Höhna et al., 2016), open-source Bayesian phylogenetics software with C++ backend and an R-like scripting language called Rev. To enable the “direct” approach, in which a supertree is inferred from a sparse average distance matrix without any intermediate distance imputation step, we re-implemented the solution previously adopted in the fitch program (Felsenstein, 1997) from the PHYLIP suite (Felsenstein, 1989) and the MW algorithm (Makarenkov and Lapointe, 2004) from the T-REX software (Makarenkov, 2001), which deals with missing distances by generalizing Eq. 2 to a weighted least-squares case. Specifically, we introduced a new data type, `AverageDistanceMatrix`, which combines an underlying distance matrix with a Boolean mask matrix of the same dimensions. The mask assigns a unit weight to the known entries of the distance matrix and a null weight to its missing entries, ensuring that the latter do not contribute to the error function. An `AverageDistanceMatrix` object can be created from a vector of distance matrices using an `fnAverageDistanceMatrix()` utility function, which also accepts an optional vector of weights equal in length to the vector of matrices, allowing the user to account for possible differences in quality among the various source datasets. The BLeSS error function is implemented as an `ExponentialErrorDistribution` over distance matrices, which is parameterized by *λ_e_* and can be “clamped” (*sensu* Höhna et al., 2014) to an `AverageDistanceMatrix` object to evaluate the likelihood of a proposed supertree. All of the above-described constructs can be freely combined with other functions, distributions, and MCMC moves already available in `RevBayes`, allowing BLeSS analyses to make use of a wide range of features including topological constraints, node age calibrations, and precomputed matrices with imputed distances.

### Simulation Tests

#### Simulation-based calibration checking

To validate our implementation of BLeSS in an easily interpretable setting, we conducted a series of simulations involving 4-tip and 5-tip trees and used simulation-based calibration checking (SBC; Talts et al., 2020; Modrák et al., 2023) to test whether the method produces unbiased posteriors. SBC builds upon the earlier posterior quantile test of Cook et al. (2006), which has seen previous use in phylogenetic settings (Slater et al., 2012b; Uyeda and Harmon, 2014; McHugh et al., 2022). Like the posterior quantile test, SBC exploits the self-recovering properties of Bayesian inference, and can be understood as extending the usual assessment of credible interval coverage (based on the expectation that, e.g., a 95% credible interval should contain the true value in 95% of simulations) by performing it for all credible interval widths simultaneously. However, compared to the posterior quantile test, SBC has the advantage of explicitly accounting for MCMC autocorrelation and the fact that the finite size of posterior samples results in discrete rather than continuous cumulative distribution functions (Gelman, 2017; Talts et al., 2020).

Simulation-based calibration checking employs a generator that repeatedly draws i.i.d. “true” values *θ̃*^(1)^*, …, θ̃*^(*N*)^ of a parameter from its prior distribution *p*(*θ*), and then simulates data *y*^(1)^*, …, y*^(*N*)^ conditional on each true value according to some sampling distribution *f* (*y* | *θ̃*). The algorithm to be validated is then used to re-infer the parameter from the simulated data, producing *N* posteriors *p*(*θ* | *y*) of *L* samples each. Because the vectors of simulated data and parameter values represent draws from the joint distribution of the model, i.e., (*θ̃, y*) ∼ *f* (*θ, y*), and because *f* (*θ, y*) = *p*(*θ* | *y*) *f* (*y*) by the axiomatic definition of conditional probability, it follows that the true values *θ̃*^(1)^*, …, θ̃*^(*N*)^ represent draws from *p*(*θ* | *y*); that is, the same posterior distribution that the algorithm is attempting to sample from. Accordingly, if the model, generator, and posterior-sampling algorithm are all correctly matched, the rank statistic of *θ̃* with respect to the posterior sample *θ*_1_*, …, θ_L_*, computed for the *i*-th case as 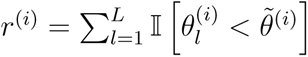, should be uniformly distributed on [0*, L*].

We implemented the SBC procedure for five different scenarios corresponding to two tree sizes (4 or 5 tips) and all possible rooted tree shapes for a given size (two shapes for the 4-tip trees, three shapes for the 5-tip trees). To facilitate interpretation, we further required that the trees conform to specific labeled tree shapes (= topologies) as well. For each scenario, we drew 100 trees from a birth-death process with a high extinction fraction, complete sampling, and an Exp(1.0) prior placed on the age of origin; following Slater et al. (2012a), the birth and death rates were set to 0.1 and 0.09, respectively. We conditioned on the number of taxa, and employed rejection sampling to ensure the draws were of the desired topology. Each simulated tree was then converted to a distance matrix **D̅** (in this case obtained directly, without averaging), which was analyzed using BLeSS with all priors set to their true values and *λ_e_* = 1. Given the small tree sizes employed in the simulation, this value was chosen to ensure that the BLeSS log likelihoods would primarily reflect *δ*(**D̂**, **D̅**) instead of being dominated by the log *λ_e_* term (Eq. 5). In addition to analyzing the complete distance matrix, we also aimed to evaluate the effect of missing distances. To this end, we removed one above-diagonal entry [*i, j*] from **D̅** at a time, producing 6 additional incomplete matrices per replicate in the 4-tip scenarios and 10 additional matrices per replicate in the 5-tip scenarios. Each of these was then analyzed under the same settings as the complete matrix. Accordingly, we carried out a total of 2 × 7 × 100 (4-tip trees) + 3 × 11 × 100 (5-tip trees) = 4700 analyses. Initially, each analysis was run with a pre-burnin period of 5000 iterations and a sampling period of 10,000 iterations, recording 1 sample every 25 iterations.

For SBC to be feasible, the parameter of interest *θ* has to be a univariate scalar, or transformable to such by some test quantity *f* (·) (Modrák et al., 2023). To validate BLeSS, we chose to focus on branch durations, or branch lengths in units of calendar time, such that for an *n*-tip tree, *θ* ∈ {*l*_1_*, …, l*_2*n*−2_}. Although time trees are commonly described in terms of node heights (divergence times), we found branch durations to be more convenient for the purposes of localizing the impact of missing distances. For a parent node *A* at height *t_A_* and its child node *B* at height *t_B_*, the duration of the connecting branch *l_A_*_→_*_B_* is simply computed as *t_A_* − *t_B_*(Cheon and Liang, 2014). To identify matching branches, we adopted the criterion proposed by Carruthers et al. (2022), whereby a branch in one tree can be considered equivalent to a branch in another tree only if the sets of tips descended from their parent nodes are identical and the sets of tips descended from their child nodes are identical as well. This definition does not require the two branches to be present in topologically identical trees, although some branches may only be compatible with a single topology in practice.

To mitigate the impact of MCMC autocorrelation, we followed Algorithm 2 of Talts et al. (2020) and standardized the effective sample size (*N*_eff_) of each branch duration to *L* states, with *L* chosen to be one less than a large power of 2 to facilitate re-binning when plotting the rank statistic distribution as a histogram. Specifically, we set *L* = 255, or one less than the first power of 2 to exceed the arbitrary but widely used *N*_eff_ threshold of 200 (Lanfear et al., 2016; Rambaut et al., 2018). Although recent results indicate that substantially higher *N*_eff_ values may be necessary to reach a desired level of precision (Fabreti and Höhna, 2021), we consider our choice of *L* to represent an acceptable trade-off between precision and chain length (directly proportional to the computational cost of the analyses). To extract effective samples from a given analysis, we first calculated *N*_eff_ using the coda R package (Plummer et al., 2006). If the *N*_eff_ value exceeded *L*, the chain was truncated; if it failed to reach *L*, the analysis was re-run with the original number of iterations multiplied by *L/N*_eff_. To keep the calculation of *N*_eff_ computationally feasible, we set the maximum allowed sampling period to 400,000 iterations. Analyses that failed to reach *L* even with this much greater chain length were repeatedly re-run for the maximum allowed number of iterations until the *N*_eff_ ≥ *L* condition was satisfied. Next, we uniformly thinned the full, correlated MCMC sample of size *N_s_* by the autocorrelation time *N_s_/N*_eff_. More sophisticated thinning strategies are possible (Talts et al., 2020; Säilynoja et al., 2022), but their performance does not appreciably differ from the traditional approach (Säilynoja et al., 2022).

While Talts et al. (2020) recommended thinning the correlated sample only once by the longest observed autocorrelation time when multiple parameters are of interest, this guidance is only applicable when every parameter is sampled at every iteration. In our case, thinning the chain just once to a single set of *L* states would result in effective sample sizes of less than *L* for at least some of the branch durations, as not every sampled tree contained all the branches of interest. Accordingly, we first extracted all samples containing a given branch, and only then applied the truncation, thinning, and rank calculation steps to the branch-specific subsample. While performing SBC separately for each branch provides a way of dealing with the interdependence between branch durations and topologies, it limits our evaluation of BLeSS to marginal rather than joint branch duration distributions, and fails to address parameter interactions (Yao and Domke, 2023).

To assess the uniformity of the rank statistics, we implemented the graphical methods proposed by Talts et al. (2020) using R code adapted from Säilynoja et al. (2022). We generated rank histograms with 20 bins and 95% confidence bands, whose lower and upper bounds were computed as the 2.5th and 97.5th percentiles of the Binom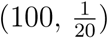 distribution, respectively. This corresponds to an expectation that, on average, no more than 1 of the 20 bins should fall outside the band. While intuitive and easy to interpret, rank histograms are known to be sensitive to the number and placement of individual bins (Säilynoja et al., 2022). We therefore also assessed the results by transforming the *r*^(*i*)^ values to fractional ranks (distributed on the unit interval), computing the empirical cumulative distribution function (ECDF) of the latter, subtracting the uniform expectation (i.e., the identity function on the unit interval), and plotting the resulting difference. To visualize expected deviations from exact uniformity, we followed Säilynoja et al. (2022) and computed 95% simultaneous confidence bands at 100 fixed evaluation quantiles (i.e., one for every ECDF value), based on inverse binomial distributions parameterized by a coverage adjustment value found via numerical optimization.

Visualizations are the preferred way of summarizing SBC results, as they make it possible to distinguish between different types and degrees of miscalibration (Modrák et al., 2023); however, more formal tests of deviation from uniformity may be desired. In their pioneering application of the posterior quantile test to phylogenetics, Slater et al. (2012b) used the Kolmogorov-Smirnov (KS) test for this purpose, but the traditional KS test is not valid when the reference distribution is discrete. To address this shortcoming, we used the recently developed exact Kolmogorov-Smirnov fast Fourier transform (Exact KS-FFT) method (implemented by the authors in the R package `KSgeneral`; Dimitrova et al., 2020), which generalizes the KS test to mixed and purely discrete reference distributions and allows calculating exact *p*-values for arbitrarily large sample sizes. We note that this generalization does not address other weaknesses of the KS test, such as its low power to detect deviations in the tails (Säilynoja et al., 2022). Moreover, since we applied the Exact KS-FFT method separately to each component of the branch duration vector and to different matrix completeness scenarios, the resulting *p*-values are subject to the multiple comparisons problem (Yao and Domke, 2023). To avoid an excessive decrease in test power and discourage a binary interpretation of the results, we opted to use uncorrected *p*-values with multiple thresholds instead of applying a formal multiple testing correction.

#### 200-tip simulations

To evaluate BLeSS in a setting more closely approaching a typical supertree analysis in terms of tree size, we used a hierarchical simulation scheme involving trees of 200 tips (Fig. 2). Specifically, we addressed four aspects of the method’s performance: (1) its sensitivity to source tree estimation error, which may cause the distances in **D̅** to deviate from ultrametricity; (2) its robustness to different ways of sampling source trees, which induce different proportions and distributions of missing entries in **D̅** ; (3) the relative success of the direct and indirect (imputation-based) approaches to dealing with missing distances; and (4) the impact of the choice of the error-penalizing rate parameter *λ_e_*. We used `RevBayes` to simulate 100 birth-death trees conditioned on the number of tips under a speciation rate (*λ*) of 0.1, an extinction rate (*µ*) of 0.09, complete sampling, and a broad U(0, 200) prior on the time of origin (*t*_or_).

**Fig. 2.**
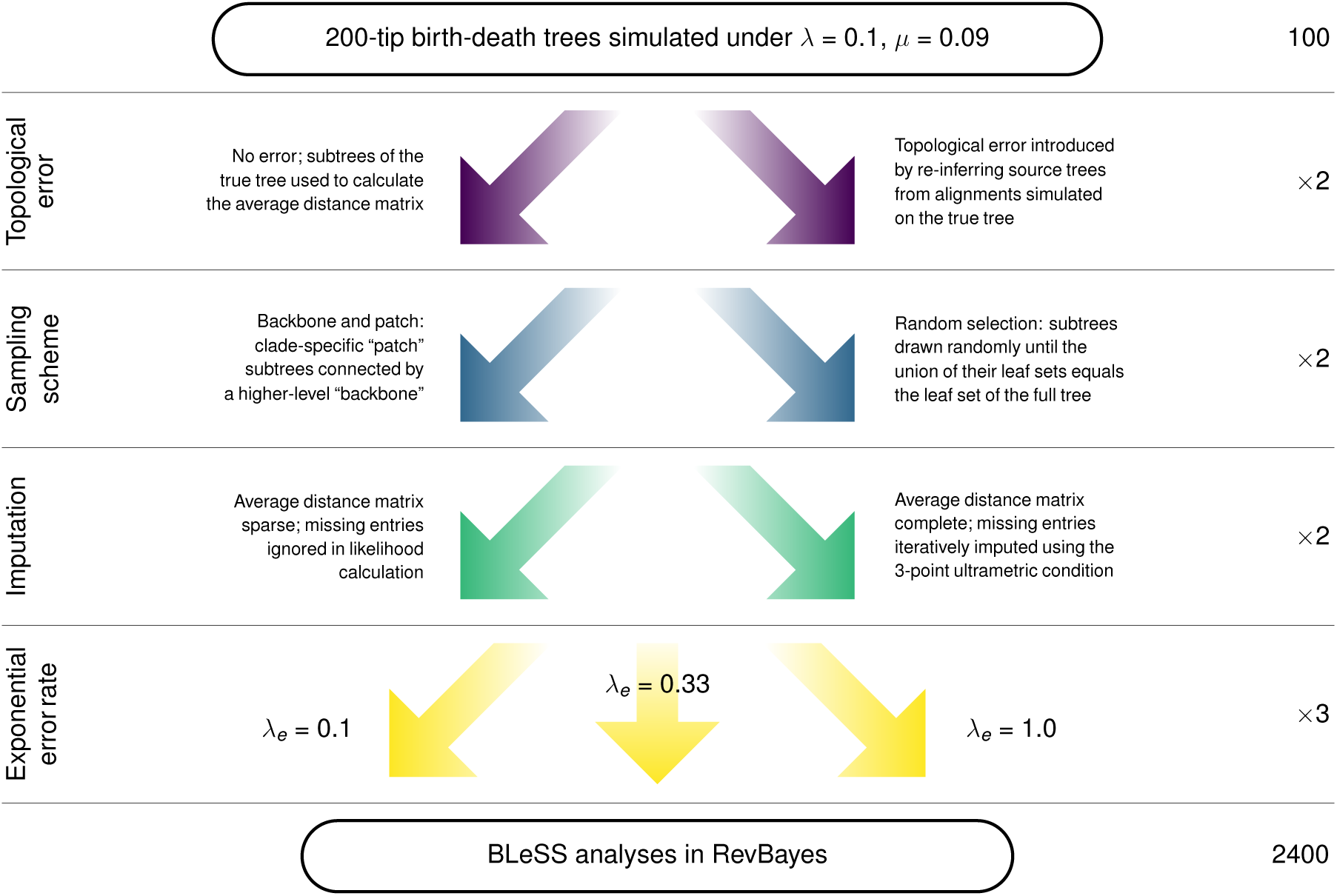
Flowchart illustrating the simulation scheme used to evaluate BLeSS.Fig. 2. Flowchart illustrating the simulation scheme used to evaluate BLeSS.

To address the impact of source tree error, we carried out two sets of simulations. In the first, source trees were obtained simply by dropping tips from each simulated phylogeny, amounting to the assumption that their topologies and divergence times were known without error. The purpose of these analyses was to identify possible biases intrinsic to the method and establish a baseline for more complex tests. In the second scenario, topological and node age error was introduced by simulating DNA sequence data on each complete tree. For each of the 100 replicates, multiple subalignments were extracted from the simulated dataset so that the subsets of included taxa corresponded to the leaf sets of the source trees created for the simulations without error. Finally, source trees were re-inferred from these subalignments using `RevBayes`. The strict correspondence between the leaf sets of the source trees used in the simulations with and without error helped ensure that any differences in BLeSS performance between the two scenarios were due to source tree quality rather than differential sampling. To make it computationally feasible to analyze the subsampled alignments in large numbers, and to generate realistic amounts of error across the full range of source tree sizes (3–60 tips), we employed a `RevBayes` re-implementation of the protocol outlined by Zhang et al. (2016). A strict clock with a rate of 0.003 substitutions per unit time was applied to each of the 100 synthetic trees to convert its branch lengths from units of calendar time to substitutions per site. The JC69 model (Jukes and Cantor, 1969) was then used to simulate 500-bp alignments on the resulting phylograms by using MCMC to sample from the unclamped phylogenetic continuous-time Markov chain distribution. A custom R script was used to break up the resulting data into subalignments, which were subsequently re-analyzed in `RevBayes`. Priors with a mean equal to the true value were used for the rate of speciation and the clock rate (*r*), the latter specified so as to ensure that the 95% highest prior density interval spanned two orders of magnitude: *λ* ∼ Exp(10), *r* ∼ Lognorm(−6.499, 1.175). The priors on the rate of extinction and origin time were slightly misspecified: *µ* ∼ Exp(10), *t*_or_ ∼ U(10, 200). The substitution process was modeled using JC69, the true model.

At the second level of the simulation scheme (Fig. 2), we further varied the manner in which each true (simulated) tree was broken up into individual source trees. First, we used a “backbone-and-patch” protocol designed to mimic the common phylogenomic approach of the same name (Jetz et al., 2012; Upham et al., 2019), in which the complete tree was subdivided into “patch” trees by making a cut at a particular height above the root. The height of the cut was randomly drawn from a uniform distribution bounded by 0 and the root height of the full tree, with rejection sampling used to ensure that the cut resulted in at least 3 patch clades of 3 or more tips each. Any tip that did not belong to a sufficiently large patch clade was instead added to the backbone, which also included one randomly selected tip from each patch clade (Fig. 3a). Second, we employed a “random selection” protocol, in which subsets of 20–60 leaves were sampled with replacement from the full leaf set of each synthetic tree until their union reached complete coverage of the original full set. The source trees were then obtained by dropping all leaves not included in a given subset from the full tree (Fig. 3b). The two sampling schemes gave rise to different distributions of missing distances in the resulting average distance matrices (Fig. 3), and to different levels of completeness (backbone-and-patch: 0.069–0.778, mean = 0.311; random selection: 0.517–0.890, mean = 0.685; see Supplementary Information, Table S1).

**Fig. 3.**
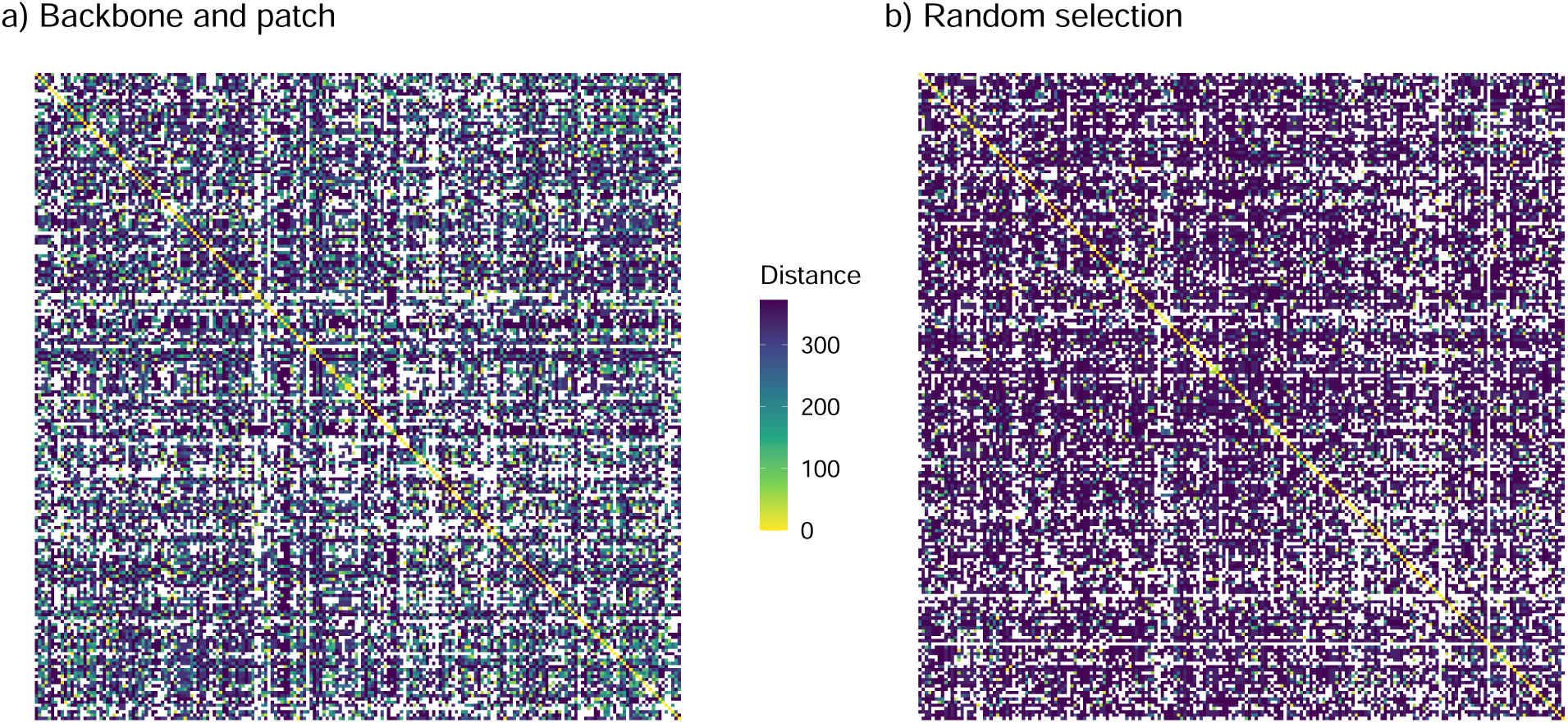
Distribution of known (filled cells) and missing (blank cells) distances in arbitrary time units under the (a) backbone-and-patch and (b) random selection sampling schemes. The replicate with median completeness was chosen to represent each scheme.

In the next stage, we repeated all previously conducted simulations on average distance matrices in which the missing entries were estimated from the available data. We used a variant of the ultrametric imputation approach (De Soete, 1984) following Landry and Lapointe (1997) and Levasseur et al. (2003), who found this algorithm to outperform the more complex additive method when dealing with higher fractions of missing data, missing distances between sister tips, or near-ultrametric matrices. The algorithm relies on the three-point condition satisfied by ultrametric distances (Hartigan, 1967):

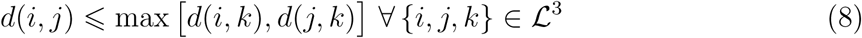

Accordingly, a missing distance between *i* and *j* can be estimated by minimizing over all triplets {*i, j, k*} for which *d*(*i, k*) and *d*(*j, k*) are known, such that

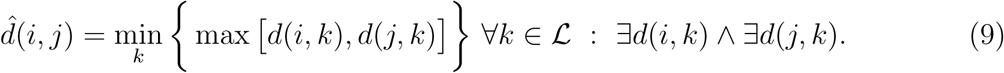

Instead of applying Eq. 9 in a fully recursive fashion, we performed at most two passes across the matrix. In the first, Eq. 9 was independently applied to each missing distance for which at least one *k* could be found, so that the imputation of one distance did not immediately affect the number of {*i, j, k*} triplets available for the imputation of the remaining distances. If no *k* could be found, a second pass was performed in which the results of the first pass were added to the known distances. The imputation algorithm was implemented in a custom R script, with precomputed average distance matrices passed to `RevBayes` as text files formatted using `phangorn` (Schliep, 2010). The imputed entries were assigned the same unit weight as the known distances.

Lastly, at the fourth and final level of the simulation scheme, we varied the value of the rate parameter of the exponential error model, *λ_e_* (Fig. 2). In effect, *λ_e_* determines the ruggedness of the likelihood surface, or the extent to which suboptimal supertrees are penalized. Smaller values of the parameter (corresponding to a larger mean) flatten the likelihood function and make it more difficult to reject estimates that poorly fit the average distance matrix, whereas larger values increase the ability to discriminate between alternative supertrees and prevent their path lengths from deviating too far from **D̅**. Our preliminary simulations suggested that large values of *λ_e_*(10–100) led to low MCMC acceptance ratios, resulting in likelihood traces marked by long plateaus and extremely slow convergence. To keep the analyses computationally tractable, we restricted our exploration of the penalty parameter to three smaller values spanning one order of magnitude: *λ_e_* ∈ {0.1, 0.33, 1.0}. Accordingly, we carried out a total of 2 × 2 × 2 × 3 × 100 = 2400 analyses. Initially, each analysis was run with a pre-burnin period of 40,000 iterations and a sampling period of 225,000 iterations, recording 1 sample every 500 iterations.

To assess MCMC convergence, we first determined the burnin proportion individually for each replicate using the previously suggested criterion (Beiko et al., 2006; Harrington et al., 2021) of discarding all samples taken before the point at which the chain first reached a likelihood higher than the mean of the last 10% of the run. After removing the burnin, we calculated the effective sample sizes (ESS) of all monitored scalar parameters using the `coda` R package. We further divided the post-burnin trace into 20 intervals of equal length, and calculated the average standard deviation of split frequencies (Lakner et al., 2008) between the last two intervals using R code adapted from the `rwty` package (Warren et al., 2017). Finally, we used `rwty` to calculate approximate topological ESS (Lanfear et al., 2016), employing the squared path difference to quantify topological distances. A given replicate was regarded as having reached convergence if ESS *>* 200 for all sampled parameters (including topology) and ASDSF *<* 0.05.

The post-burnin tree traces of all replicates that satisfied the convergence criteria described above were summarized as maximum clade credibility (MCC) and maximum *a posteriori* (MAP) supertrees. These summary supertrees were then compared to the corresponding true tree using a variety of statistics: the Robinson-Foulds (RF; Robinson and Foulds, 1981) and Kuhner-Felsenstein (KF; Kuhner and Felsenstein, 1994) distances, as implemented in `phangorn` (Schliep, 2010); the path length difference (Steel and Penny, 1993), as implemented in the R package `castor` (Louca and Doebeli, 2017); the matching split (Bogdanowicz and Giaro, 2012; Lin et al., 2012) and clustering information (Smith, 2020a) distances, as implemented in the package `TreeDist` (Smith, 2020b); the Billera-Holmes-Vogtmann (BHV) metric (Billera et al., 2001), as implemented in the package `distory` (Chakerian and Holmes, 2012); the symmetric quartet divergence (Smith, 2019a), as implemented in the package `Quartet` (Sand et al., 2014; Smith, 2019b); the Kendall-Colijn (KC) metric (Kendall and Colijn, 2016), as implemented in the package `treespace` (Jombart et al., 2017) and with the contribution of branch lengths to the distance ranging from 0 to 1 in increments of 0.1; and the difference between the true and inferred root height, calculated using the package `phytools` (Revell, 2012).

To effectively summarize the results, which involved assessing the relative importance and marginal effects of potentially covarying discrete and continuous predictors, we employed Bayesian additive regression trees (BART; Chipman et al., 2010; Bleich et al., 2014; Hill et al., 2020) as implemented in the R package `bartMachine` (Kapelner and Bleich, 2016). BART is a nonparametric sum-of-trees machine learning method, in which every individual regression tree represents a “weak learner” whose contribution to explaining the variation in the response variable is constrained by a regularization prior to prevent overfitting. We used both the RF and KF distances between the estimated and true supertrees as response variables, as these metrics are only imperfectly correlated with one another (Supplementary Information, Fig. S10) and present distinct advantages. In contrast to the RF distance, the KF distance quantifies both topological and branch length estimation error, but unlike the former, it cannot be normalized, and is therefore less easy to interpret. In addition to the four categorical variables described above (topological error, source tree sampling scheme, missing data treatment, and *λ_e_* value), we also used two continuous predictors: true tree imbalance, as quantified by the Colless index (Colless, 1982), and **D̅** completeness, defined as the fraction of non-missing entries prior to imputation. Only the replicates that reached convergence according to the criteria listed above were used for the BART analyses, resulting in highly unequal sample sizes for the different scenarios. For both analyses (of RF and KF distances), we randomly allocated 80% of the response and predictor values to the training set and the remaining 20% to the testing set. We used out-of-sample root-mean-square error to choose the optimum number of trees (testing numbers from 20 to 120 in increments of 2), and conducted an MCMC analysis with a burnin period of 120,000 iterations and a sampling period of 120,000 iterations under the best tree number and with all other hyperparameters set to their default values (*α* = 0.95, *β* = 2, *k* = 2, *q* = 0.9, *ν* = 3; Kapelner and Bleich, 2016). Following standard practice (Chipman et al., 2010), we used the proportion of times each predictor was chosen for a splitting rule (out of all splitting rules and across the entire posterior sample) as a measure of that predictor’s importance, and partial dependence plots to visualize the marginal effects of continuous predictors.

In addition to evaluating point estimates (summary supertrees), we also aimed to assess each posterior sample as a whole in terms of node age coverage. Since our simulations involved the simultaneous inference of tree topology and divergence times, we specifically focused on the age of the root, which represented the only internal node that was guaranteed to be present in all sampled supertrees regardless of their topology. Coverage probability was calculated as the proportion of replicates in which the true root age fell into the corresponding 95% highest posterior density (HPD) interval, out of all those that reached convergence. Additionally, we computed the 95% HPD intervals for all nodes in a given MCC tree that were also present in the true tree, and determined what percentage of these contained the corresponding true ages. Näıvely, one may expect this to be the case for 95% of nodes on average if the method is well-behaved, with higher proportions indicating overly diffuse posteriors (too much uncertainty) and lower proportions indicating overly narrow posteriors (too much certainty). However, since ancestors necessarily predate their descendants, individual nodes of the same tree do not represent independent replicates in terms of age, and topological mismatches between the true tree and the summary tree may cause the same proportion to correspond to very different rates of overall inferential success. We mitigated the latter problem by calculating the relevant proportions both out of correctly recovered nodes only and out of all internal nodes present in the tree (*n* = 199), but while the resulting percentages remain informative of the overall performance of the method, they cannot be interpreted strictly as coverage probabilities.

#### Scalability tests

In addition to testing for implementation correctness and accuracy, we performed one final set of simulations focusing on computational efficiency to evaluate how well BLeSS scales with respect to tree size. Specifically, we extended one of the scenarios explored in the 200-tip simulations (no topological error, backbone-and-patch sampling scheme) to trees of 500, 1000, 5000, and 10,000 tips. As before, true trees were simulated under *λ* = 0.1, *µ* = 0.09, and complete sampling, and conditioned on the number of tips. Since the simulations were aimed at assessing computational speed rather than accuracy, only a single tree was generated for each scenario (Supplementary Information, Table S8). The process of creating the backbone and patch trees was similar to that used in the 200-tip simulations, except that a minimum of 10 patch clades of at least 10 tips each was required.

First, we performed MCMC-free benchmarks to determine the amount of time required for those operations that are carried out upfront and only once per analysis (e.g., computing **D̅**, drawing a tree from a prior distribution), as well as those operations that collectively make up a single MCMC iteration (converting a proposed tree to a path-length matrix, and computing the natural logarithm of its probability density under the exponential error distribution). All measurements employed the internal `RevBayes time()` function, which provides a Rev wrapper around the microsecond clock from the Boost C++ library (http://www.boost.org). For each combination of tree size (500, 1000, 5000, and 10,000 tips) and missing data treatment (imputation vs. no imputation), 10 replicates with different random seeds were performed. The benchmarks were conducted with millisecond precision on a single compute node consisting of Intel Xeon Gold 6248R processors, with one thread per replicate. To confirm the current implementation of BLeSS gains no performance boost from parallelization, we repeated the benchmarks using the MPI (Message Passing Interface) version of `RevBayes` with 32 threads per replicate (Supplementary Information, Table S10).

Second, we ran 5 chains with different random seeds on each of the 8 datasets (4 tree sizes × 2 missing data treatments) for 5 million iterations or 180 hours, whichever was reached first. In contrast to the settings used for the SBC and 200-tip analyses, the `moveschedule` argument of the MCMC function was set to “`single`” so that each proposed move would be counted as a separate iteration (Höhna et al., 2017). Sampling frequencies were scaled so as to obtain a comparable number of samples per unit runtime for all tree sizes (Supplementary Information, Table S9). The `mnScreen` monitor was used to write elapsed time to the standard output, which was redirected to a plain-text file. These files were subsequently parsed and used to compute the number of moves per unit runtime. In addition, we computed the previously proposed statistic of ESS increment per unit runtime (Aberer et al., 2016; Fourment et al., 2018) for all monitored scalar parameters by applying the `effectiveSize` function from the `coda` package to cumulative subsamples of each MCMC run. Due to limitations imposed by the cluster on which the analyses were performed, the 180-hour period consisted of 36-hour windows, with the `RevBayes` checkpointing functionality used to restart each chain from the last state saved before a timeout. Extra iterations that were run after the last saved state but before the timeout were included when calculating the total number of moves per unit runtime but omitted from the calculation of effective sample sizes. All MCMC analyses were carried out on the same hardware as the MCMC-free benchmarks.

Third, we evaluated sampling efficiency by recording the acceptance rates of the various MCMC moves used to update both the supertree itself and the hyperparameters of the birth-death prior from which the supertree is drawn. Deviations from the default `RevBayes` target rate of 0.44 can be due to tree size-related problems with the exploration of the parameter space, which may in turn reflect genuine features of the posterior: low acceptance rates of topology moves may be indicative of the chain having reached a tree with a high posterior probability that is difficult to improve upon, while high acceptance rates may occur when the data are insufficient to discriminate among different topologies (Harrington et al., 2021). The acceptance rates were calculated from chains of different lengths for different tree sizes, without a dedicated pre-burnin period or autotuning.

### Empirical Application to the Order Carnivora

As an empirical demonstration, we used BLeSS to sample the posterior distribution of time-calibrated supertrees for extant representatives of the mammalian order Carnivora based on a set of published time-scaled phylogenies. Carnivora is a compelling system in which to explore our approach for two reasons: several relatively complete time-scaled molecular phylogenies are available for comparison (Agnarsson et al., 2010; Slater and Friscia, 2019; Upham et al., 2019), and the order has been the focus of several previous supertree efforts (Bininda-Emonds et al., 1999; Nyakatura and Bininda-Emonds, 2012; Akanni et al., 2015).

We first performed a literature review to identify time-calibrated source trees that could be included in the profile. We ignored studies that obtained complete or near-complete sampling of the order to ensure that not all entries in the average distance matrix were filled, though studies that achieved relatively complete sampling of families or superfamilies were included. In order to be considered for inclusion, we required that one of two possible sources of data be associated with the paper: a tree file containing the topology annotated with branch lengths, or else a figure of the topology with mean divergence times provided in the text or in an associated table that would enable manual transcription of the time tree. If divergence times were provided only for a subset of figured nodes, we entered the topology corresponding to that subset of bipartitions into the profile. In cases where multiple sets of divergence times were provided, typically corresponding to different calibration schemes, we recorded all. We did not require the source trees to be based on any particular data type or time calibration approach, although we did note both. Our decision to reject source studies that did not provide a time tree in a usable format, rather than downloading, aligning, and reanalyzing the sequence data used in them, was motivated by a desire to replicate the situations in which BLeSS, or any other supertree method, might most profitably be used in place of analysis of primary data.

Our literature review identified 31 studies for which at least one tree file was available or from which at least one time tree could be manually reconstructed based on figures and tables provided. Our final profile included 33 trees from 30 source studies, with one outlier study excluded because it yielded unusually young divergence times. After removing duplicates and domesticated taxa, the union of the leaf sets of these trees comprised a total of 260 species, including a few latest Pleistocene taxa that we treated as extant for the purposes of this study. Sampling was variable among families. Based on the American Society of Mammalogists’ Mammalian Diversity Database (Burgin et al., 2018, v1.91; downloaded July 8, 2022), we achieved complete sampling of extant Hyaenidae (*n* = 4), Mephitidae (*n* = 14), Phocidae (*n* = 19), Prionodontidae (*n* = 2), and Ursidae (*n* = 8) among families with two or more species. The sampling fraction for other families containing more than two species was variable, ranging from as high as 0.96 in Felidae to as low as 0.29 in Herpestidae. Low sampling in some families is not due to a lack of available studies, but instead reflects a lack of associated tree files or divergence time estimates for figured trees in those studies that are available. A full description of source studies evaluated for inclusion is provided in the Supplementary Information.

As in our simulation study, we performed BLeSS analyses with and without imputation. Several of the source studies in our profile of carnivoran time trees are based on mitochondrial DNA data alone, which have been suggested to yield divergence time estimates that are too old for younger nodes and too young for older nodes (Fulton and Strobeck, 2010). We therefore performed a second set of BLeSS analyses in which the contribution of mitogenome-based source trees to the average distance matrix was downweighted by a factor of 10. These downweighted distance matrices were again analyzed both with and without imputed distances. Based on our simulation studies, we set *λ_e_* = 0.1 for the main analyses. However, as mentioned above, an alternative strategy is to marginalize over *λ_e_*. We therefore performed a final analysis using the average distance matrix derived from downweighted mitochondrial data and with imputed missing distances (see Results for rationale) while integrating out *λ_e_*, which was drawn from an Exp(1.0) prior.

For each analysis, we placed exponential priors on the speciation and extinction rates, *λ* ∼ Exp(10), *µ* ∼ Exp(10). We used an offset exponential prior for the root age, *t*_or_ ∼ Exp(0.1645, offset = 38), where the offset approximately corresponds to the first appearance dates of the canid *Hesperocyon gregarius* and the amphicyonid *Daphoenus lambei* (Tomiya, 2011), and the rate parameter is scaled so as to place the 99th percentile at the Cretaceous–Paleogene boundary (66 Ma). We performed 4 simultaneous runs to ensure convergence, with a pre-burnin period of 40,000 iterations and a sampling period of 500,000 (no imputation + no downweighting and imputation + downweighting + estimated *λ_e_*) or 1,000,000 iterations (all other scenarios). Parameter posteriors were visually inspected as trace plots, and convergence was further assessed based on the ESS and ASDSF values calculated for scalar parameters and sampled topologies, respectively. Combining all 4 runs almost always resulted in extremely low ESS values for the log likelihood (LnL), log prior (LnPr), and their sum (the log posterior), but moderate or high effective sample sizes for the remaining scalar parameters and low ASDSF values, in agreement with previous observations that there is little correlation between LnL ESS and topological convergence (Harrington et al., 2021). In such cases, we combined the posterior samples from all 4 runs as long as their ASDSF did not exceed 0.01, and computed MAP and MCC trees both from the pooled sample and from the single run with the highest post-burnin LnL values. For those analyses where ASDSF was *>* 0.01, only the run exhibiting the highest LnL values was summarized.

To determine how the constrained least-squares estimator underlying BLeSS compares to a simple application of existing distance-based methods to an average distance matrix, we also inferred carnivoran supertrees using neighbor-joining (NJ; Saitou and Nei, 1987) as implemented in phangorn (Schliep, 2010). NJ represents a natural point of comparison, as it was historically viewed as a fast approximation to ordinary least squares (OLS) (Felsenstein, 2004), although more recent studies demonstrated that it actually seeks to optimize a different criterion (Gascuel and Steel, 2006) which allows it to outperform OLS in practice (Willson, 2005). Since existing implementations of NJ can deal neither with the ultrametricity constraint nor with sparse matrices (though see Criscuolo and Gascuel, 2008), we only estimated NJ supertrees from imputed average distance matrices (both with and without downweighting mtDNA-based source trees) and focused only on the topology of the inferred supertree when comparing the results to those of BLeSS. The latter was necessary because the lack of ultrametricity prevents the branch lengths recovered by NJ from being interpreted as amounts of time elapsed between successive divergences, thus rendering them essentially meaningless.

A common problem with evaluating method performance in empirical settings is the fact that in contrast to simulations, the true topology remains unknown. We adopted the approach of Agnarsson and May-Collado (2008) and evaluated the analyses by computing the fraction of “benchmark” clades recovered in the summary supertrees. Such clades have to be of undisputed monophyly, supported by multiple lines of evidence, and selected prior to the analysis. We recognized a total of 53 benchmark clades with ranks ranging from suborder to subtribe; the full list is given in the Supplementary Information (Table S13) along with the expected composition of each clade in terms of our species sample.

## Results

### Simulation Tests

#### Simulation-based calibration checking

The number of iterations needed to reach the target effective sample size for all branch lengths of interest varied as a function of tree size, tree symmetry, and matrix completeness. Across all five scenarios, reaching the target ESS took a greater number of iterations on average for analyses performed on the complete **D̅** than for analyses with missing distances. On average, the shortest runs were required for the 4-tip balanced-topology scenario (i.e., ((1, 2), (3, 4)); ∼49,930 iterations), while the longest runs characterized the 5-tip scenario with a single tip sister to a balanced subtree (i.e., (1, ((2, 3), (4, 5))); ∼233,830 iterations). For the latter scenario, achieving high ESS values proved to be particularly challenging, and a number of replicates had to be re-run multiple times with the maximum allowed chain length (400,000 iterations, corresponding to 16,000 total samples). This difficulty in inducing BLeSS to sample the true imbalanced topology in a trivial case is surprising given the previously reported bias of average distance matrix methods in favor of pectinate source trees (Wilkinson et al., 2005). However, it is not unexpected if the posterior ends up being dominated by the birth-death prior, which is uniform over labeled histories, and accordingly favors balanced topologies, which are compatible with more labeled histories than imbalanced ones (Velasco, 2008).

The graphical summaries generally indicate that the posteriors inferred by BLeSS are well-calibrated, with the empirical CDF mostly falling within the envelope of theoretical expectation (Fig. 4) and only a small number of rank histogram bins extending outside the approximate confidence interval. In the 4-tip balanced-topology scenario, a near-exact match between the empirical and theoretical CDFs was observed when analyzing the complete **D̅**. Even in the cases of mismatch, the observed failure modes were predictable from the properties of the underlying data, such as when the removal of a distance between two terminal sister taxa resulted in excessive certainty in the duration of their branches, producing a strongly significant deviation from uniformity according to the Exact KS-FFT test (the “no *d*(1, 2)” case in Fig. 4). This illustrates the well-known problem that missing distances between terminal sister taxa (“cherries” *sensu* Erdős et al., 1997) pose to all average consensus methods, as there is no information in the rest of the matrix that can be exploited to estimate the length of the corresponding branches (Lapointe and Kirsch, 1995; Lapointe and Levasseur, 2004). As expected, the posteriors and their summaries (i.e., rank histograms and ECDF plots) were identical for each pair of branches forming a cherry, reflecting the fact that their durations were not estimated independently but simply obtained by halving the corresponding distance.

**Fig. 4.**
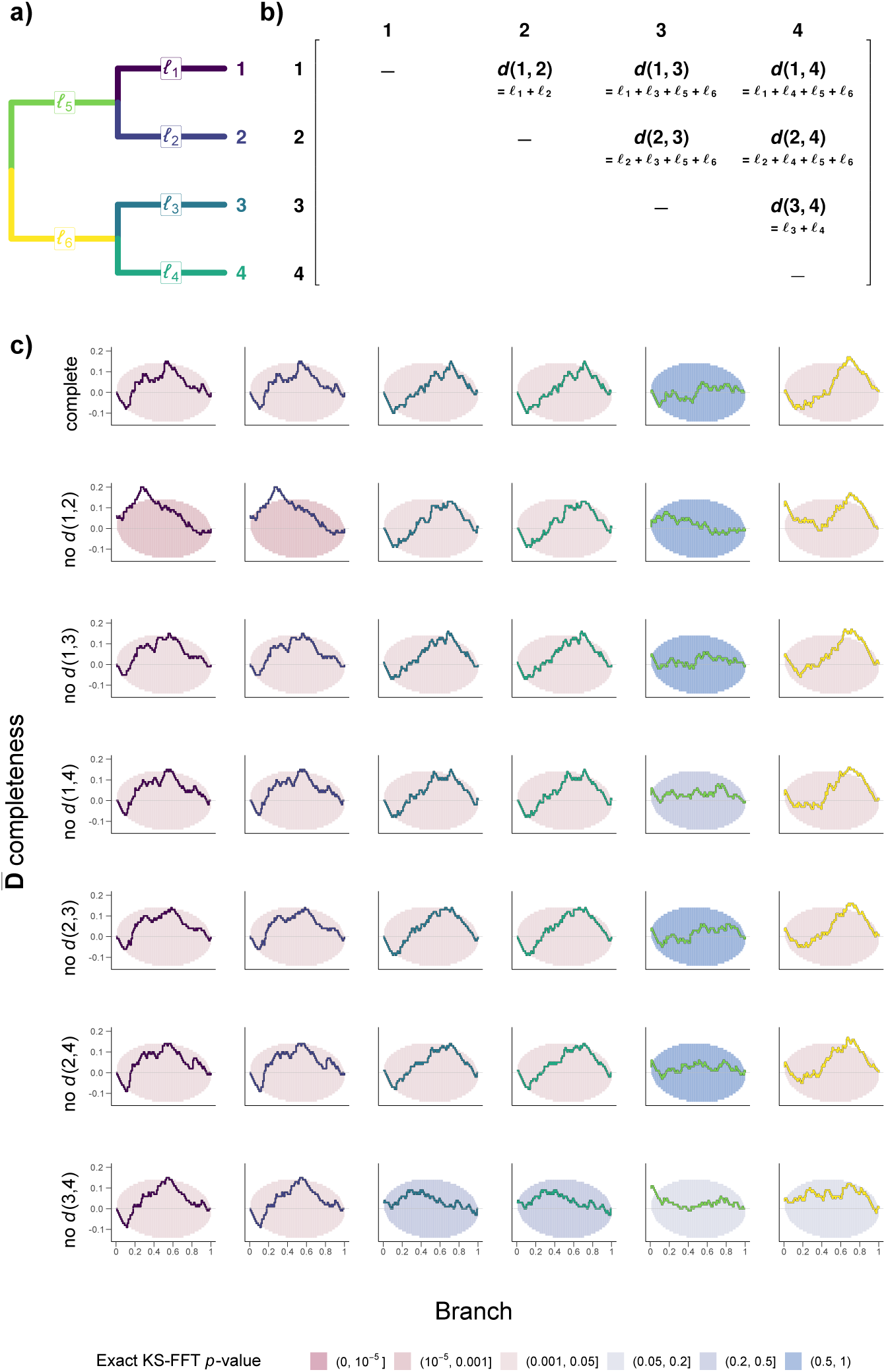
Results of simulation-based calibration checking for the 4-tip balanced tree scenario. (a) The topology with labeled branches. (b) The corresponding distance matrix, with observed distances expressed as sums of the underlying branch durations. (c) The difference between the empirical cumulative distribution function (ECDF) and the discrete uniform expectation (stepwise identity function) with 95% simultaneous confidence bands for individual branch durations and different average distance matrix (**D̅**) completeness scenarios. The confidence bands are colored according to the *p*-values from uniformity tests conducted using the exact Kolmogorov-Smirnov fast Fourier transform (Exact KS-FFT) method.

When the true tree is imbalanced, BLeSS posteriors largely remain free of bias but may exhibit excessive uncertainty, as indicated by concave rank histograms and S-shaped ECDF curves around the diagonal. In fully pectinate 4-tip and 5-tip trees, this problem was particularly severe for the branch subtending the earliest-diverging tip (Supplementary Information: *l*_5_ in Figs. S2, S3 and *l*_7_ in Figs. S6, S7), whose duration is equivalent to the root height of the tree. The next most strongly affected branch subtended the second earliest-diverging tip (Supplementary Information: *l*_3_ in Figs. S2, S3 and *l*_5_ in Figs. S6, S7), suggesting a trend of excessive uncertainty progressively accumulating with increasing node depth. Compared to the 4-tip balanced-topology scenario, the matrices computed from fully pectinate trees were much less sensitive to the removal of individual entries, including even the distance between the two tips of the only cherry present in such trees (Supplementary Information: the “no *d*(3, 4)” case in Figs. S2, S3 and the “no *d*(4, 5)” case in Figs. S6, S7). The remaining two scenarios, which involved 5-tip trees intermediate between fully balanced and fully pectinate topologies, also showed minimal sensitivity to distance removal but less clear overall trends (Supplementary Information, Figs. S4, S5, S8, S9). In particular, the scenario with a single tip sister to a balanced subtree gave rise to the only instance of bias observed in our SBC analyses, which affected the branches subtending the two cherries (Supplementary Information: *l*_5_ and *l*_6_ in Figs. S4, S5). The durations of these two branches exhibited posteriors with an overabundance of particularly high estimates, and thus low ranks (Supplementary Information, Fig. S5). Plausibly, this bias may be related to the convergence issues reported above for the same scenario, or to the solution we adopted in response to these issues (i.e., re-running the inference until a sufficient number of effectively independent samples was obtained within a specified chain length).

**Fig. 5.**
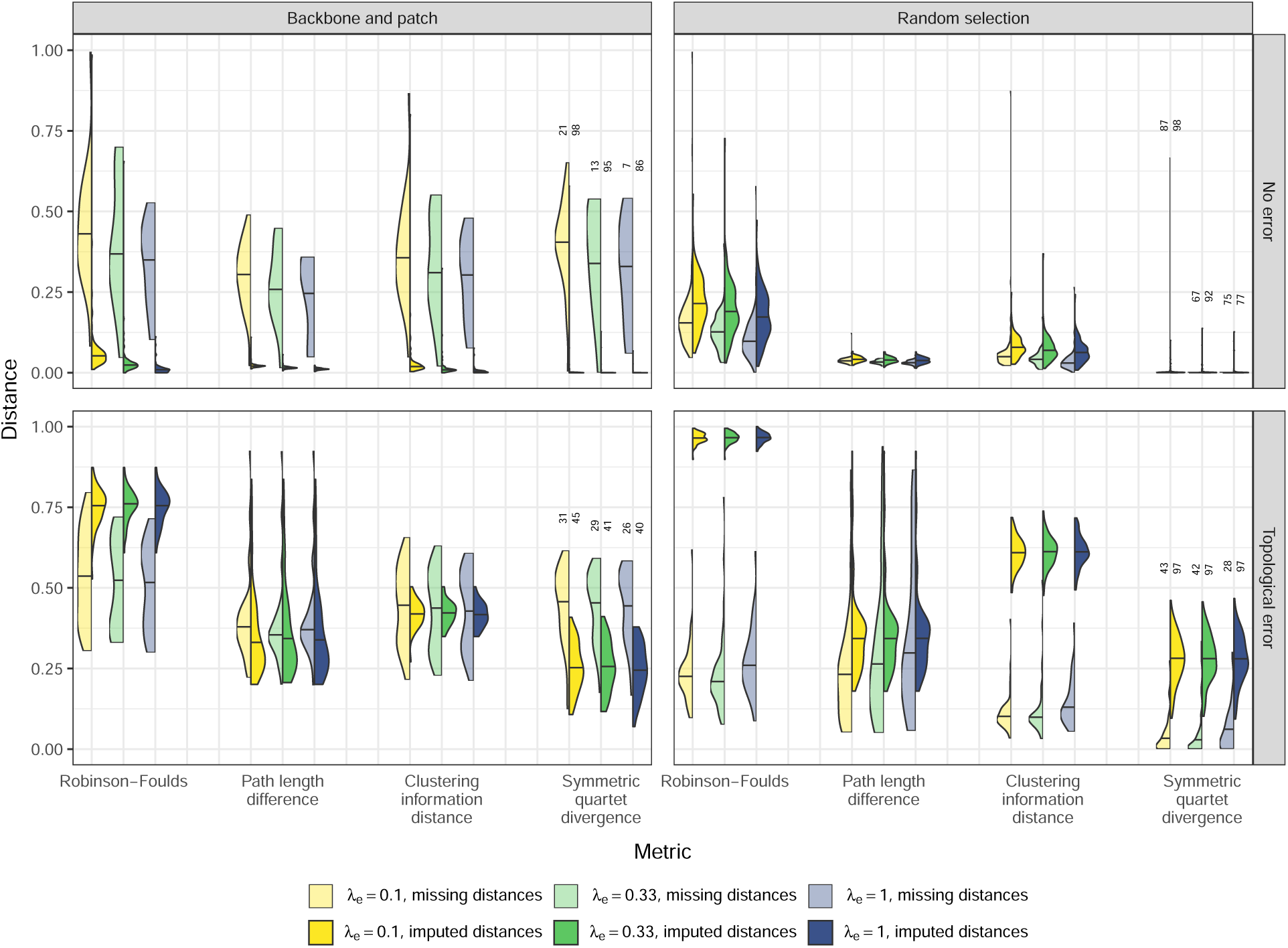
Split violin plots showing the distributions of accuracy metrics across the 24 scenarios illustrated in Fig. 2. All metrics are normalized and expressed as distances (rather than similarities) between the true (simulated) supertree and the BLeSS maximum clade credibility estimate, so that 0 indicates perfect agreement and 1 indicates no agreement. The numbers above the rightmost set of violin plots indicate the number of replicates that reached convergence for a given scenario.

#### 200-tip simulations

The 2,400 BLeSS analyses of simulated large-scale trees were run for 225,000–1,350,000 generations, with 1,432 analyses reaching convergence according to the predefined criteria (see Materials and Methods). The proportion of replicates satisfying our convergence metrics was subject to considerable variation, ranging from 7 out of 100 to 98 out of 100. The rate of achieving convergence was generally higher for analyses with rather than without imputation, and for analyses conducted under lower values of *λ_e_* (Fig. 5). A preliminary examination of the results showed that the MCC trees slightly but consistently outperformed the MAP trees (Supplementary Information, Tables S2–S4), and we therefore restricted subsequent evaluation to the former. For ease of interpretation, we primarily focused on those distance metrics that can be normalized to the unit interval (with 0 indicating perfect agreement and 1 indicating no agreement) when comparing the inferred and true supertrees (Fig. 5); the values for the remaining metrics are given in the Supplementary Information (Tables S5–S7; Fig. S21). The RF and KF distances were additionally investigated using BART analyses. These reached excellent convergence (Supplementary Information, Figs. S13, S14) when run with the optimum number of regression trees (RF distances: *m* = 56; KF distances: *m* = 62), but their residual distributions exhibited significant deviations from normality (Supplementary Information, Fig. S15), suggesting that prediction intervals and other output that assumes normally distributed noise may be of limited reliability. Indeed, while the 95% prediction intervals contain the actual response values from the testing set in 89.2% of cases for the RF model, indicating good predictive performance, this is only true in 74.1% of cases for the KF model (Supplementary Information, Fig. S16).

The mean values of normalizable distance metrics vary widely across the 24 simulation scenarios, ranging from 0.012 to 0.965 for the RF distance, from 0.012 to 0.408 for path length difference, from 0.006 to 0.613 for the clustering information distance, and from 0.001 to 0.446 for the symmetric quartet divergence (Fig. 5). When accounting for the well-known tendency of the RF metric to rapidly saturate (Steel and Penny, 1993; Smith, 2020a), which helps explain the very large distances observed in several scenarios, these values generally indicate moderate to excellent accuracy. At the same time, the corresponding scenario-specific distributions exhibit a considerable overlap, suggesting that the variation in the properties of individual simulated datasets (replicates) was at least as impactful as the differences in analytical choices (Fig. 5).

This is confirmed by the BART analyses, which show the imbalance of the true tree and the completeness of the corresponding average distance matrix to be the two most important predictors of both RF and KF distances, as determined by the frequency with which they were chosen for regression tree splitting rules (Supplementary Information, Fig. S17). Similarly, all three variable selection procedures suggested by Bleich et al. (2014) consistently identify tree imbalance and **D̅** completeness as the only two predictors to be included in both the RF model and the KF model (Supplementary Information, Fig. S20). Nevertheless, comparing the pseudo-*R*^2^ value of the original model to a null distribution derived from 50 permuted datasets shows that all predictors (including the four categorical variables corresponding to the different treatments outlined in Fig. 2) have a significant effect on the RF distance between the true and inferred supertrees, and all but two (*λ_e_*: *p* = 0.98; and source tree sampling scheme: *p* = 0.96) have a significant effect on the corresponding KF distance. As expected, the partial dependence function shows that the RF distance between the true and inferred supertrees decreases with increasing **D̅** completeness in an almost linear fashion, but the relationship is less straightforward for the corresponding KF distance (Supplementary Information, Fig. S19). After marginalizing out the contributions of the remaining predictors, the contribution of true tree imbalance to the two accuracy metrics is also highly non-monotonic (Supplementary Information, Fig. S19). While the average consensus method was previously shown to resolve within-profile conflicts in favor of imbalanced source trees (Wilkinson et al., 2005), our estimates of the partial dependence function average the effect of true tree imbalance over supertrees derived from both perfectly congruent and conflicting source trees, possibly accounting for the complex relationship observed.

Qualitative comparisons show that in the absence of source tree conflict, *λ_e_* has the expected effect on topological accuracy, with larger values corresponding to smaller distances between the true tree and the BLeSS summary supertree (Fig. 5). However, this effect is attenuated when topological error is introduced into the profile, likely because forcing the estimated supertrees to adhere more closely to **D̅** is of lesser utility when the **D̅** entries themselves deviate from the branch durations of the true tree because of source tree conflict. The BART analyses of the RF distances between the true and estimated supertrees confirm that of all categorical predictors (i.e., variables other than **D̅** completeness or true tree imbalance), *λ_e_* interacts most strongly with the presence or absence of source tree error, although the same is not true of the corresponding KF distances (Supplementary Information, Fig. S18). In the latter case, interactions of *λ_e_* with categorical predictors are infrequent, reflecting the fact that the penalty parameter has no significant effect on the response and is rarely used as a splitting variable in the first place (Supplementary Information, Figs. S17, S18).

Complex interactions between the presence or absence of error in the profile, source tree sampling scheme, and missing data treatment are apparent from the scenario-specific distributions of accuracy metrics (Fig. 5) as well as the BART analyses (Supplementary Information, Fig. S18). The best-performing combinations of treatments include backbone-and-patch analyses with imputation and no source tree error, followed by random-selection analyses with no source tree error or imputation (see Fig. 6 for an example). Under the backbone-and-patch sampling scheme, the imputation of missing **D̅** entries strongly improves BLeSS accuracy in the absence of source tree error, but its effect is more ambiguous when error is introduced into the profile, with most metrics (especially the symmetric quartet divergence) still showing improvement but the RF distance indicating strongly degraded performance. In contrast, under the random-selection sampling scheme, imputation always results in lower accuracy, with the difference being relatively minor in the absence of source tree error but increasing to such an extent in its presence that the distributions of several metrics are completely disjunct between the two missing data treatments (Fig. 5). The BART analyses of both RF and KF distances show the interactions of source tree error and missing data treatment to be the most important pairwise interactions between any two categorical predictors, followed by interactions between missing data treatment and sampling scheme, and finally by interactions between sampling scheme and source tree error (Supplementary Information, Fig. S18).

**Fig. 6.**
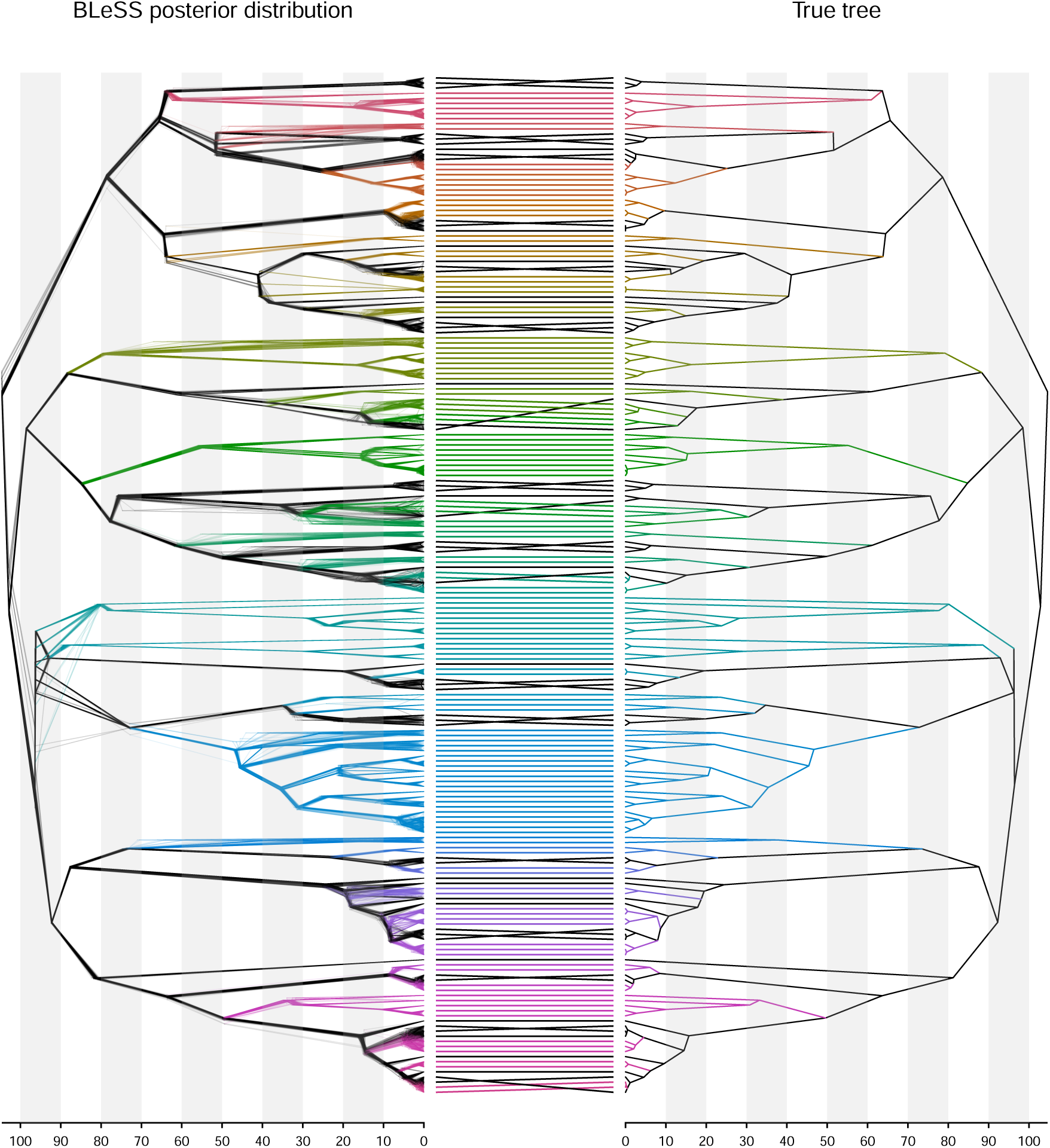
Tanglegram illustrating the topological and node height accuracy of a random BLeSS replicate conducted without imputation on randomly selected error-free source trees under *λ_e_* = 0.1. A DensiTree (Bouckaert, 2010) representation of 50 samples from the BLeSS posterior is superimposed onto the maximum clade credibility (MCC) supertree computed from all samples on the left, and compared to the true tree on the right. Edges of the same color indicate the most inclusive non-nested subtrees that are congruent between the true tree and the MCC supertree; the inter-tree edges connect corresponding tips, and their crossings indicate topological conflict. The numbers along the *x*-axis represent arbitrary units of node height.

The observed interplay between imputation and source tree error may appear intuitive: when the available **D̅** entries are free of error (perfectly ultrametric), the missing entries can be imputed without error as well, thus increasing the completeness of **D̅** while still allowing it to perfectly fit the true supertree. A more complete average distance matrix can better discriminate between near-optimal estimates, leading to more accurate inference. In contrast, using ultrametric estimation to impute missing entries based on distances that deviate from ultrametricity is likely to compound pre-existing error. This expectation is partly borne out by decomposing the topological error in the inferred supertrees into two components: the mean topological distance between the source trees re-estimated from simulated sequences and the restrictions of the true supertree to their leaf sets, and the mean topological distance between the source trees and the restrictions of the inferred supertree to their leaf sets. The second component quantifies additional error introduced by the BLeSS inference itself, and is dramatically increased by imputation (Table 1). However, this expected relationship is modulated by the influence of the source tree sampling scheme. When source trees are sampled at random, overall error as well as its second component are low when missing entries are treated as such, and increase especially drastically with imputation. The backbone-and-patch sampling scheme is less sensitive to imputation, as both types of error are higher in its absence, but increase to a lesser extent when it is employed (Fig. 5, Table 1). Plausibly, this differential effect of imputation might not be directly due to the sampling scheme itself, but rather to its covariates such as **D̅** completeness, which strongly interacts with it (Supplementary Information, Fig. S18) and whose distribution differs markedly between the two schemes (Supplementary Information, Table S1).

**Table 1.**
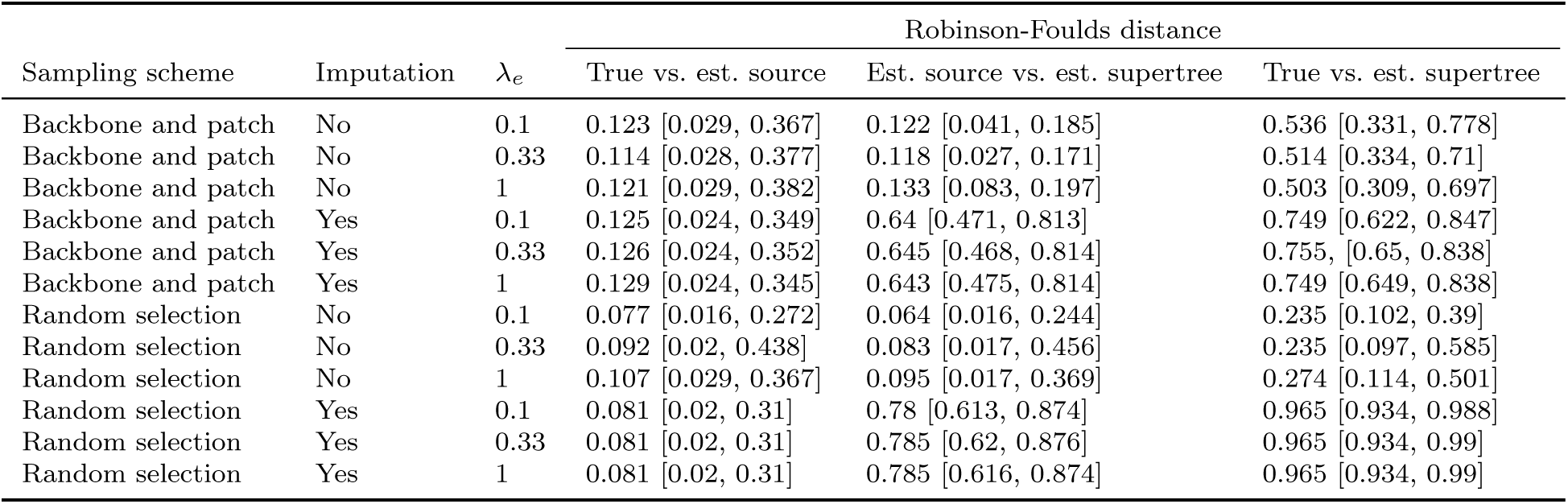
The relative contributions of source tree error and BLeSS error to the topological error in the estimated supertree (*𝒯̂*) in those simulations where source tree error is present. The contribution of source tree error is quantified as the normalized Robinson-Foulds (RF) distance between a source tree *𝒯_k_* and *𝒯̃|ℒ*(*𝒯_k_*) (the restriction of the true tree to the leaf set of *𝒯_k_*), averaged across all source trees (“True vs. est. source”). The contribution of BLeSS error is quantified as the normalized RF distance between *𝒯_k_* and *𝒯̂|ℒ*(*𝒯_k_*), averaged across all source trees (“Est. source vs. est. supertree”). The overall error is quantified as the normalized RF distance between *𝒯̂* and *𝒯̃* (“True vs. est. supertree”). Every value is reported as the mean and 95% interpercentile range over all replicates that reached convergence. While the distances between *𝒯_k_* and *𝒯̃|ℒ*(*𝒯_k_*) do not vary with imputation or *λ_e_*, the values for different scenarios are averaged over different sets of replicates, resulting in different means and 95% ranges.

| Sampling scheme | Imputation | $\lambda_e$ | Robinson-Foulds distance | | |
| --- | --- | --- | --- | --- | --- |
|  |  |  | True vs. est. source | Est. source vs. est. supertree | True vs. est. supertree |
| Backbone and patch | No | 0.1 | 0.123 [0.029, 0.367] | 0.122 [0.041, 0.185] | 0.536 [0.331, 0.778] |
| Backbone and patch | No | 0.33 | 0.114 [0.028, 0.377] | 0.118 [0.027, 0.171] | 0.514 [0.334, 0.71] |
| Backbone and patch | No | 1 | 0.121 [0.029, 0.382] | 0.133 [0.083, 0.197] | 0.503 [0.309, 0.697] |
| Backbone and patch | Yes | 0.1 | 0.125 [0.024, 0.349] | 0.64 [0.471, 0.813] | 0.749 [0.622, 0.847] |
| Backbone and patch | Yes | 0.33 | 0.126 [0.024, 0.352] | 0.645 [0.468, 0.814] | 0.755, [0.65, 0.838] |
| Backbone and patch | Yes | 1 | 0.129 [0.024, 0.345] | 0.643 [0.475, 0.814] | 0.749 [0.649, 0.838] |
| Random selection | No | 0.1 | 0.077 [0.016, 0.272] | 0.064 [0.016, 0.244] | 0.235 [0.102, 0.39] |
| Random selection | No | 0.33 | 0.092 [0.02, 0.438] | 0.083 [0.017, 0.456] | 0.235 [0.097, 0.585] |
| Random selection | No | 1 | 0.107 [0.029, 0.367] | 0.095 [0.017, 0.369] | 0.274 [0.114, 0.501] |
| Random selection | Yes | 0.1 | 0.081 [0.02, 0.31] | 0.78 [0.613, 0.874] | 0.965 [0.934, 0.988] |
| Random selection | Yes | 0.33 | 0.081 [0.02, 0.31] | 0.785 [0.62, 0.876] | 0.965 [0.934, 0.99] |
| Random selection | Yes | 1 | 0.081 [0.02, 0.31] | 0.785 [0.616, 0.874] | 0.965 [0.934, 0.99] |

In terms of root age accuracy and coverage, the single largest difference is observed between the simulations conducted with and without source tree error, with further influence exerted by the sampling scheme (Table 2). The correlation between true and estimated root ages is strong under all scenarios (*R*^2^ range: 0.68–1), but introducing error into the profile leads to the overestimation of small root heights (by a factor of up to 21.2) and underestimation of large root heights (by a factor of up to 1.6; Supplementary Information, Fig. S22). In the absence of source tree error, the ratio of estimated to true root ages tends to approximate 1 extremely closely (Supplementary Information, Tables S2–S4), and the relative precision of the estimates (width of the 95% HPD interval divided by the true age) is extremely high (range of scenario-specific means: 0.0003–0.0060). While these values mostly translate to high coverage probabilities, this is not the case for analyses conducted under the backbone-and-patch sampling scheme and without imputation, for which the normalized 95% HPD interval widths are still narrow (mean values of 0.0060, 0.0040, and 0.0034 for *λ_e_*= 0.1, 0.33, and 1, respectively) but the ratio of estimated to true root ages appreciably deviates from 1 (mean values of 0.803, 0.849, and 0.939 for *λ_e_* = 0.1, 0.33, and 1, respectively). The introduction of source tree error does little to reduce relative precision (range of scenario-specific means: 0.0006–0.0103) but results in drastically degraded accuracy (Supplementary Information, Tables S2–S4). Since younger root ages tend to be overestimated to a much greater extent than old root ages tend to be underestimated, the scenario-specific mean ratios of estimated to true values always exceed 1 (range: 1.419–2.109). This combination of inaccurate but extremely precise estimates yields coverage probabilities that are uniformly zero, meaning that the true values are never included in the corresponding 95% HPD intervals (Table 2). For the ages of the remaining nodes, the trend is similar but somewhat attenuated; in those replicates where at least some of the nodes present in the true tree are recovered in the MCC supertree, a nonzero number of such correctly recovered nodes is often also associated with 95% HPD age intervals that contain the true value (Supplementary Information, Fig. S23). This may be due to the tendency of age estimates to become less precise for younger nodes (see below and Fig. 7).

**Table 2.** Root age coverage probability (proportion of simulation replicates in which the true root age falls within the 95% highest posterior density interval) across the 24 scenarios illustrated in Fig. 2. The number of replicates which reached convergence for a given scenario and from which the coverage probability was calculated is given in parentheses next to the respective value.

| Topological error | Sampling scheme | Imputation | Root age posterior coverage ($n$) | | | | | |
| --- | --- | --- | --- | --- | --- | --- | --- | --- |
| | | | $\lambda_e = 0.1$ | $\lambda_e = 0.33$ | $\lambda_e = 1$ | | | |
| No | Backbone and patch | No | 0.095 (21) | 0.077 (13) | 0.286 (7) |  |  |  |
| No | Backbone and patch | Yes | 0.99 (98) | 1 (95) | 1 (86) |  |  |  |
| No | Random selection | No | 0.989 (87) | 1 (67) | 1 (75) |  |  |  |
| No | Random selection | Yes | 1 (98) | 1 (92) | 1 (77) |  |  |  |
| Yes | Backbone and patch | No | 0 (31) | 0 (29) | 0 (26) |  |  |  |
| Yes | Backbone and patch | Yes | 0 (45) | 0 (41) | 0 (40) |  |  |  |
| Yes | Random selection | No | 0 (43) | 0 (42) | 0 (28) |  |  |  |
| Yes | Random selection | Yes | 0 (97) | 0 (97) | 0 (97) |  |  |  |

**Fig. 7.**
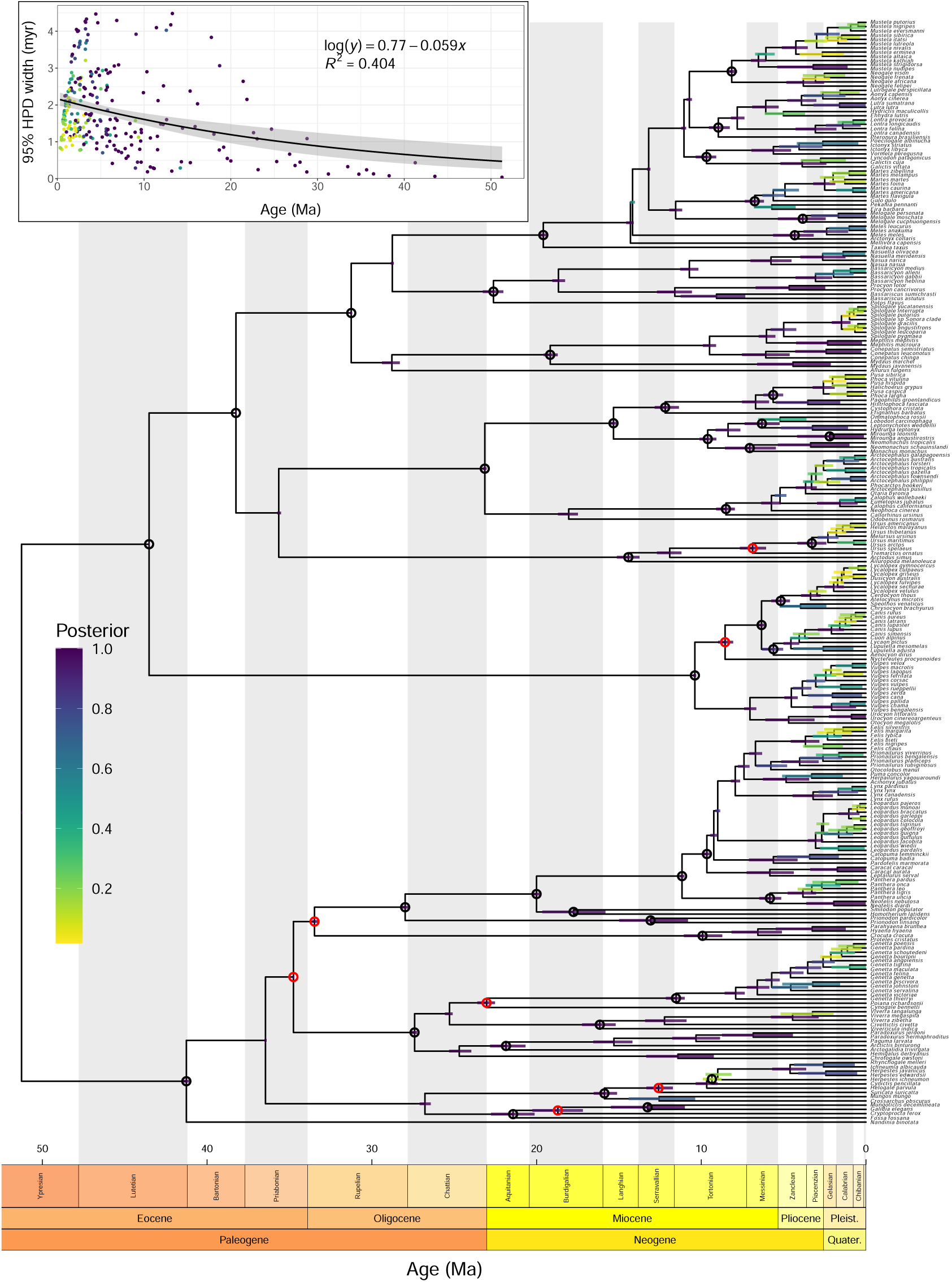
Results of a BLeSS analysis of 260 species of extant carnivorans. Main panel: the maximum clade credibility tree with mean node heights. The horizontal bars denote the 95% highest posterior density (HPD) intervals for the ages of internal nodes, and are colored by their posterior probability. The circles indicate nodes congruent with (black) or contradicting (red) the 53 predefined benchmark clades. Inset: the relationship between the precision of node age estimates (quantified as the width of the corresponding 95% HPD intervals) and their depth in the tree.

#### Scalability tests

Encouragingly, our benchmarks show that the most time-intensive tasks needed for conducting BLeSS inference are those that only need to be performed once at the beginning of each analysis, including the calculation of the average distance matrix **D̅**, the initial draw from the birth-death prior, and fixing (or “clamping”; Höhna et al., 2014) a realization of the exponential error model to an observed value in the form of **D̅** (Table 3). In actual analyses of very large average distance matrices, the real computational bottleneck is likely to consist of the conversion of a proposed supertree to a distance matrix (𝒯̂ → **D̂**); at 10,000 tips, this operation is 13.5 times (imputation) to 35 times (no imputation) more time-consuming than the calculation of the least-squares difference that underlies the BLeSS likelihood function (Table 3). None of the steps involved in BLeSS inference currently benefits from the native RevBayes parallelization via MPI (Supplementary Information, Table S10). As expected, the wall-clock runtime of BLeSS scales approximately with the square of tree size, with little difference between missing data treatments (i.e., sparse or dense average distance matrices; Supplementary Information, Table S11).

**Table 3.**
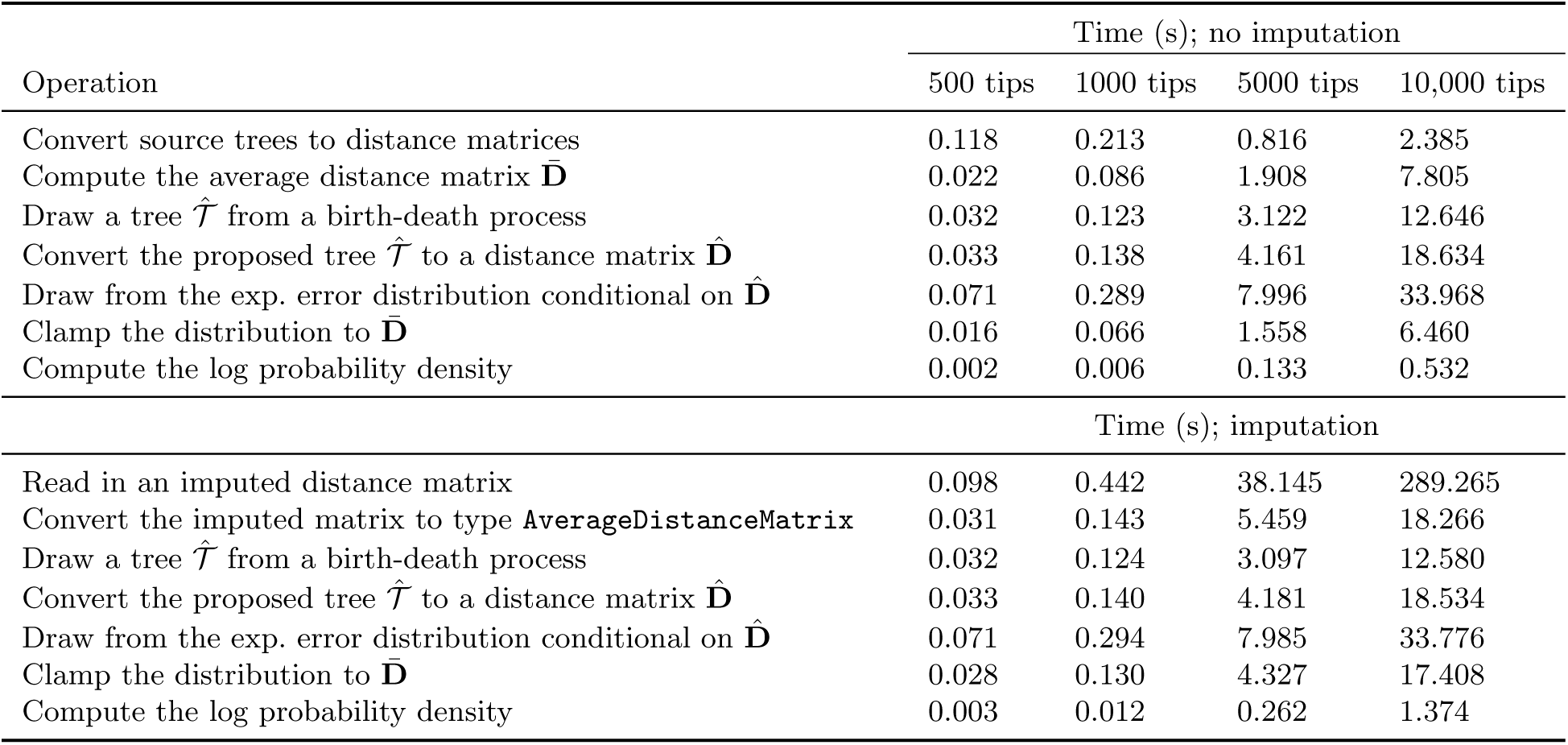
Wall-clock time spent on individual operations underlying BLeSS as a function of supertree size and missing data treatment. All values are averaged over 10 replicates and measured with millisecond precision for the single-core version of RevBayes v1.2.2 (commit b083532).

At 34.5–38.8 and 8.8–9.2 moves per second, respectively, BLeSS analyses of 500-tip and 1000-tip supertrees take less than one week to complete 5 million single-move MCMC iterations (Supplementary Information, Fig. S24), a number that is generally sufficient to attain moderate (*>*200) to good (*>*625; Fabreti and Höhna, 2021) effective sample sizes for LnL and the root age, though not for the hyperparameters of the birth-death process (Supplementary Information, Fig. S25). Note, however, that because of computational constraints, we did not calculate approximate topological ESS (Lanfear et al., 2016) per unit runtime, so sampling efficiency remains unclear for a key parameter of interest. Moreover, even for the remaining parameters, our simplistic calculation of ESS increments per unit runtime likely underestimated sampling efficiency by not treating any of the initial iterations as pre-burnin (a period used for move tuning rather than sampling) or burnin (early samples to be discarded). Except for the speciation rate scaler, most moves exhibit acceptance rates that increase as a function of supertree size (Supplementary Information, Table S12). However, based on an examination of the corresponding LnL traces, this relationship largely reflects the fact that the sampler had not yet burned in for very large supertrees (of 5000 or 10,000 tips), and proposed moves were highly likely to increase the likelihood of the initial low-probability parameter samples. For supertrees of 500 and 1000 tips, the acceptance rates of almost all moves fall short of the default target of 0.44 (by a factor of up to ∼5400 for the root age), resulting in relatively inefficient sampling (Supplementary Information, Table S12). In practice, this problem can likely be mitigated by autotuning.

### Empirical Application to the Order Carnivora

Some general trends were observable in the convergence diagnostics for the carnivoran analyses with and without imputed distances, regardless of whether source studies based on mitochondrial data were downweighted or not. Topological convergence across runs, as measured by ASDSF, was very good (*<* 0.01) for analyses without imputation, but less so when missing distances were imputed (0.01 *<* ASDSF *<* 0.015). However, analyses without imputation took longer to reach stationarity, yielding lower per-run effective sample sizes for most parameters. In general, within-chain post-burnin ESS values were moderate (*>*200) for speciation and extinction rates as well as the root age, regardless of missing data treatment. Due to the large burnin proportions required for analyses without imputed distances, within-chain ESS values were low (∼ 100) for LnL, LnPr, and their sum (log posterior), while they exceeded 200 in analyses based on the imputation of missing distances.

A common pattern where ESS values dropped for all but birth-death parameters and the root age when combining runs showed that each chain sampled a slightly different part of the joint posterior distribution. This effect was weakest when missing distances were treated as such (ESS *>* 100 in most cases), and strongest when they were imputed (ESS *<* 10 for some parameters). We therefore summarized only the single run with the highest post-burnin LnL values for analyses based on imputed distances, while both combining chains and summarizing the highest-likelihood chain alone for those without imputation (Supplementary Information, Table S14). For the analysis in which *λ_e_* was estimated, a visual inspection of the LnL traces indicated there was no need to discard initial samples as burnin, and combining all 4 chains resulted in ESS *>* 450 for all scalar parameters as well as an ASDSF of *<* 0.001. We therefore combined the chains for this analysis prior to summarizing. As in the 200-tip simulations, we found that when the MAP and MCC supertrees differed from each other, it was more often the latter that represented the more accurate point estimate (in terms of the number of benchmark clades recovered; Supplementary Information, Table S14). Accordingly, we focus on the MCC supertrees when describing the relevant results.

Evaluation of benchmark clade recovery in MCC supertrees reveals that inference based on average distance matrices without imputed distances performed poorly. Regardless of whether we evaluated supertrees from combined runs or the single highest-likelihood run, the number of benchmark clades recovered was low (30–32%; *n* = 16–17). Missing benchmark clades spanned a range of taxonomic levels; at higher levels, we failed to recover a monophyletic Feliformia, Caniformia or Arctoidea, while at lower levels we missed a number of tribes and subtribes. Downweighting source trees inferred from mtDNA sequences had a limited effect on the MCC supertree derived from combined chains (36%; *n* = 19), but substantially increased the number of benchmark clades recovered in the summary tree from the highest-likelihood run (58%; *n* = 31), albeit with a similar hierarchical spread of missing clades. In contrast, supertrees inferred from average distance matrices with imputed missing distances consistently yielded a large number of benchmark clades (85–87%; *n* = 45–46), with the highest number found in the MCC tree based on downweighted mitochondrial source trees (Fig. 7). This point estimate narrowly outperformed the NJ supertrees inferred with and without downweighting mtDNA-based source trees, both of which were consistent with the monophyly of 44 benchmark clades (83%) when appropriately rerooted. The clades that BLeSS failed to recover even with imputation included not only subfamilies (Euplerinae, Hemigalinae, Mungotinae, as well as Herpestinae in the analysis without downweighting) and tribes (Arctotheriini, Vulpini) but also the infraorder Viverroidea (Viverridae + Herpestoidea) and superfamily Herpestoidea (Hyaenidae, Eupleridae, Herpestidae), which were contradicted by an unusual set of interfamilial relationships within feliforms. In contrast, both NJ supertrees additionally lack a monophyletic Galidinae, Canini, Phocini, and Canina, but do recover Arctotheriini, Viverroidea, and Herpestoidea (Supplementary Information, Figs. S26, S27). No benchmark clades were recovered in the analysis where *λ_e_* was estimated.

Comparisons with previous species-level phylogenies of Carnivora (Supplementary Information, Figs. S28–S30) show that in terms of benchmark clade recovery, the best BLeSS estimate is intermediate between the MRP supertree of Nyakatura and Bininda-Emonds (2012) (42 out of 51 applicable clades; 82%) and the backbone-and-patch tree of Upham et al. (2019) (48 out of 52 applicable clades; 92%); as expected, the supermatrix-based tree of Slater and Friscia (2019) outperforms all other methods (43 out of 44 applicable clades; 98%) (Supplementary Information, Table S14). Pairwise comparisons indicate that the best BLeSS estimate is closer to the supermatrix-based tree of Slater and Friscia (2019) than to the phylogenies obtained with supertree or supertree-like approaches according to almost all metrics, but that the phylogeny of Slater and Friscia (2019) is not always closer to the BLeSS estimate than to the MRP and backbone-and-patch supertrees (Supplementary Information, Table S15). This is true even when the differences in the number of tips shared by each pair of trees are eliminated by pruning all taxa other than those that are common to all four studies (Supplementary Information, Table S16).

Similar to topological accuracy, the precision of divergence time estimates was also affected by the treatment of missing data. For analyses that treated missing distances as such, the mean precision of node age estimates is lower in both absolute (mean width of the 95% HPD intervals: 2.01 myr) and relative (mean ratio of 95% HPD interval widths to the corresponding posterior means: 0.881) terms, and there is a weak negative relationship (mean slope = −0.026; mean *R*^2^ = 0.014) between posterior mean estimates and their absolute precision, which yields a slow decrease in expected HPD interval width from ∼ 2.2 myr (mean intercept) at shallow nodes to ∼ 0.8 myr at the root node. However, a stronger negative relationship of greater magnitude is found for inference based on imputed missing distances (mean slope = −0.047; mean *R*^2^ = 0.145), which also yielded more precise estimates (mean absolute precision = 1.83 myr, mean relative precision = 0.783). Our preferred estimate illustrates that the actual difference between the widest (Galidinae: 4.48 myr; relative precision = 0.338) and narrowest (root: 0.0347 myr; relative precision = 6.77 × 10^−4^) HPD intervals is larger than predicted by linear regression, and exponential regression produces a better fit (*R*^2^ = 0.404; Fig. 7). A similar relationship is observed between node depth and node support, with the youngest and oldest nodes receiving the lowest and highest posterior probabilities, respectively (Fig. 7). Both correlations are expected, as deep nodes are informed by a greater number of **D̅** entries than shallow ones, making it more difficult to fit a supertree containing a contradictory branching order or divergence times. The imputation of missing distances further increases this number, rendering the correlation even stronger.

The posterior mean root ages estimated by BLeSS (43.4–56.1 Ma) are plausible in light of the carnivoran fossil record and previous molecular dating studies. In particular, our preferred estimate of 51.3 Ma (associated with the supertree estimated from downweighted mitochondrial source trees and with imputed missing distances; Fig. 7) falls within the 95% HPD interval yielded by a recent phylogenomic study that was not included in the profile (Álvarez-Carretero et al., 2022). The extremely narrow 95% HPD interval about the root age never includes the prior mean (44.1 Ma) in the analyses where *λ_e_* was fixed; however, in the analysis where *λ_e_* was estimated as a free parameter, the posterior (43.4 Ma) and prior mean almost perfectly coincide, further suggesting that the resulting estimates are driven by the birth-death prior.

## Discussion

BLeSS marries two previously introduced approaches to supertree inference (Lapointe and Levasseur, 2004; Steel and Rodrigo, 2008) into a single flexible framework that allows previously inferred time trees to be combined in a principled and statistically rigorous manner. Because it recasts the problem in a Bayesian setting, BLeSS addresses the two main shortcomings of existing supertree methods – the lack of branch lengths and the reliance on point estimates without accounting for the associated uncertainty – by providing the joint posterior distribution of supertree topologies and divergence times. This property allows it to move beyond the traditional dichotomy between “veto” and “voting” methods that has long featured in the supertree literature (Ranwez et al., 2007; Brinkmeyer et al., 2011), since a posterior sample of BLeSS supertrees needs neither to represent topological conflict as a lack of resolution nor to eliminate it by deciding in favor of one topology or the other. We provide a performant implementation of BLeSS in RevBayes (Höhna et al., 2016), validate it using a rigorous simulation-based calibration checking pipeline, and demonstrate its practicality for problems involving up to *>* 1000 tips. Using an extensive set of 200-tip simulations, we further investigated the influence of a variety of analytical settings. Over the range of values we tested, the penalty parameter *λ_e_* has only a weak effect on accuracy that is offset by its impact on the rate of MCMC convergence (Fig. 5; see below). The ability of BLeSS to incorporate source trees that overlap more extensively than those employed by the backbone-and-patch approach (Jetz et al., 2012; Tonini et al., 2016; Upham et al., 2019) represents a clear advantage, as shown by the fact that simulations with source trees whose leaf sets comprised randomly selected subsets of the full leaf set yielded more complete average distance matrices (Supplementary Information, Table S1) that led to more accurate inference of both topologies and divergence times (Fig. 5; Table 1). The effect of imputing missing distances via ultrametric estimation (De Soete, 1984; Lapointe and Kirsch, 1995) is less straightforward and mediated by a number of other variables; however, at least in the empirical application of BLeSS presented here, imputation appears to outperform inference from a sparse average distance matrix even when using conflicting source trees.

Our analyses of both synthetic and empirical datasets show BLeSS to generally perform well under a wide range of conditions, albeit with several caveats. These are illustrated especially well by the empirical case, in which both the width of the 95% HPD intervals on node ages and topological uncertainty declines from the tips to the root (Fig. 7). The high precision of the age estimates for deep nodes may result in overconfidence and low coverage (i.e., lower than nominal probability of the true value falling within a credible interval). The tendency to overestimate the ages of shallow nodes and underestimate the ages of deep nodes in the presence of source tree conflict, which was revealed by our simulations (Supplementary Information, Fig. S22), is also concerning and merits further investigation. At the same time, the flexibility of our implementation leaves open the possibility that this bias might be partially or completely mitigated by measures such as node calibration or downweighting of outlier source trees. Such analytical practices are particularly well-suited to the empirical application illustrated in this study, which involves estimating a time tree that is more densely sampled and/or spans a more inclusive clade from pre-existing time trees that are less densely sampled and/or span less inclusive clades. However, it is worth noting that BLeSS may also be applicable to other problems for which Bayesian supertrees have been proposed as a solution, such as devising divide-and-conquer strategies for *de novo* analysis of large datasets (Karcher et al., 2021) or the inference of species trees from incongruent gene trees (De Oliveira Martins et al., 2016). Investigating the statistical properties of BLeSS in the latter setting may be particularly worthwhile given recent results showing that relatively simple species tree estimators perform well even in circumstances that render more sophisticated supermatrix analyses statistically inconsistent (Dasarathy et al., 2015; Allman et al., 2019). Below, we outline several other analytical considerations, practical recommendations, and possible extensions that are broadly applicable to a variety of BLeSS use cases.

### Influence of the exponential error rate parameter

Despite exploring a relatively narrow range of possible *λ_e_* values, our analyses of simulated and empirical data show this penalty term to exert influence on supertree estimation in terms of both accuracy and MCMC performance. While treated here as a parameter that is internal to the BLeSS model and determines the shape of its posterior, *λ_e_* is closely related to auxiliary variables employed in MCMC tempering techniques to sample from distributions other than the posterior, which helps explain its behavior and provide guidance regarding its treatment.

To the extent that the log *λ_e_* term in Eq. 5 is negligible relative to the *λ_e_δ*(**D̂, D̅**) term, *λ_e_* approximates the temperature parameter *t* used to construct “power posterior” distributions of the form *f_t_*(*θ*|*y*) = *p*(*θ*)*f* (*y*|*θ*)*^t^* between the unnormalized posterior (*t* = 1) and the prior (*t* = 0) (Gelman and Meng, 1998; Friel and Pettitt, 2008). Such power posteriors are commonly used in Bayesian model comparison to estimate marginal likelihoods using path-sampling (Gelman and Meng, 1998; Lartillot and Philippe, 2006) and stepping-stone (Fan et al., 2011; Xie et al., 2010) techniques. Like the temperature parameter, *λ_e_* scales the likelihood relative to the prior. This feature is potentially problematic, since manipulating *λ_e_* could in theory be used to induce a posterior that is dominated to an arbitrary extent either by the user’s prior beliefs about the branching process, or by the information contained in the average distance matrix. In practice, however, MCMC mixing imposes constraints on the range of values that are viable for any given analysis (see below).

While the arbitrariness in the choice of *λ_e_* can conceivably be circumvented by estimating it from the data as a free parameter, our attempts to do so in preliminary analyses of the carnivoran dataset yielded unsatisfactory results. On the one hand, the posterior, which was approximately lognormal with a narrow peak close to 0 (mean = 5.4 × 10^−8^, 95% HPD = [6.60 × 10^−10^, 1.37 × 10^−7^] based on the pooled posterior sample from 4 runs), diverged sharply from the prior (a diffuse exponential with a mean of 1), suggesting that the average distance matrix is informative with respect to the value of *λ_e_*(Supplementary Information, Fig. S15). This is not unexpected, as distances averaged across source trees with a high degree of conflict will substantially deviate from ultrametricity, and any ultrametric supertree will thus only fit them imperfectly, favoring low values of *λ_e_*. Conversely, low amounts of conflict allow the supertree to strictly adhere to the near-ultrametric average distance matrix, making it possible to impose a heavier penalty (corresponding to a high value of *λ_e_*) for deviations therefrom. On the other hand, because the estimated penalty was extremely low, the posterior was entirely dominated by the birth-death prior, whose contribution exceeded that of the BLeSS likelihood term by more than 1800 log units. As a result, the analysis in which the supertree and *λ_e_* were co-estimated performed extremely poorly in practice, failing to recover even the best-supported benchmark clades that received a posterior probability of 1 in the analyses where *λ_e_* was fixed.

In addition to determining the prior-to-likelihood ratio, *λ_e_* also modulates the ruggedness of the likelihood surface. To the extent that the prior is negligible relative to the likelihood, its behavior approximates the *β* parameter (also referred to as “temperature”) employed in Metropolis-coupled MCMC (also known as MC3 or parallel tempering; Geyer, 1991). Unlike power posterior sampling, MC3 raises the entire posterior (rather than just the likelihood term) to a given temperature, producing a “heated” or “tempered” distribution of the form *f_β_*(*θ*|*y*) = *p*(*θ*)*^β^f* (*y*|*θ*)*^β^*. Choosing *β* ∈ (0, 1) has the effect of flattening the target distribution, making it easier to explore multimodal posteriors by traversing the low-probability valleys between local optima (Altekar et al., 2004). If swaps are attempted between the chain sampling from the posterior and parallel chains sampling from the tempered distributions, the former can benefit from the rapid mixing of the latter (Gilks and Roberts, 1996). These improvements in mixing closely resemble the behavior we observed with low values of *λ_e_*. In particular, our 200-tip simulations show that decreasing *λ_e_* improves mixing (quantified by the number of replicates satisfying our convergence criteria) at the cost of accuracy (quantified by the distance between the true and estimated supertrees) (Fig. 5). The same trade-off was also apparent in the empirical analyses. When *λ_e_* was freely estimated, its extremely low values made it easy for the 4 chains to converge to the prior-dominated posterior (ESS *>* 450, potential scale reduction factor *<* 1.015 for all scalar parameters; ASDSF *<* 0.001). In contrast, analyses performed with fixed values of *λ_e_* often yielded parameter traces in which sudden shifts punctuated long periods characterized by a very low amount of variation. Similarly, different chains almost always exhibited non-overlapping traces, with among-chain variation exceeding within-chain variation by several orders of magnitude. As a result, pooling the resulting samples resulted in extremely low ESS values. In our final analyses, this problem mostly affected LnL traces, whereas the chains mixed relatively well for the main parameters of interest such as the root height and (based on the ASDSF) topology. However, in several preliminary analyses (*λ*_0_ = 0.1–0.33, ∼325,000 iterations), the same behavior was also observed for the root age, with different chains sampling from different distributions even though the difference between the means of these distributions (*<* 0.2 myr) was well within the desired margin of error.

By analogy with *t* and *β*, we believe that *λ_e_* is better treated as an auxiliary variable exterior to the model, and that *λ_e_*= 1 represents the most natural choice when the unmodified posterior is of interest. If this relatively large value leads to convergence issues, these should be addressed using techniques such as MC3 rather than by manipulating *λ_e_* itself. One consequence of always setting the penalty term to a unit value regardless of tree size is that the posterior will be dominated by the prior for small supertrees (as observed in the SBC analyses with 4–5 tips) and by the likelihood for large supertrees (as observed in the benchmark analyses with *>* 1000 tips). This fact is probably best interpreted as a genuine feature of the relationship between data and evidence under the BLeSS model: larger average distance matrices can discriminate among trees much more effectively than smaller ones. A fruitful avenue for further research would be to investigate the sensitivity of BLeSS to the choice of the time unit (Groussin et al., 2011). The functional forms of both the BLeSS likelihood (Eq. 5) and the birth-death prior contain exponentiated branch durations, and both will therefore respond to a change of the time unit, possibly in ways that affect their relative weights.

### Potential efficiencies

Our simulations suggest that the current implementation of BLeSS is practical for supertrees of up to ∼1000 tips. This is approximately equivalent to the largest tree sizes at which full Bayesian co-estimation of tree topology and divergence times from sequence data remains feasible (e.g., du Plessis et al., 2021; Wisniewski et al., 2022), and substantially smaller than the recent megaphylogenies inferred by the time-scaling of maximum-likelihood trees or backbone-and-patch techniques (Jetz et al., 2012; Tonini et al., 2016; Rabosky et al., 2018; Upham et al., 2019). At first, BLeSS would therefore seem to offer little advantage over existing methods. Our benchmarks show it is the conversion of a proposed tree to a distance matrix that represents the main computational bottleneck in the current implementation of the method (Table 3). Accordingly, future efforts to improve the performance of the method should focus on exploiting efficiencies that can be achieved at this step.

Importantly, standard tree-perturbing MCMC moves implemented in RevBayes, such as nearest neighbor interchange (NNI), subtree prune and regraft (SPR), a whole-tree scaler, or a node time slider, either only affect several branches at a time, leaving large subtrees of the original tree intact, or scale all branches by the same factor. In both cases, recomputing **D̂** every time a tree has been updated is clearly wasteful. In the analyses of character data, this problem is circumvented by reusing partial likelihoods stored at the corresponding nodes or branches (Smith et al., 2024). However, unlike the likelihoods that result from modeling character substitution along the branches of a tree, which are computed using Felsenstein’s pruning algorithm (Felsenstein, 1981), the BLeSS likelihoods (Eq. 5) are not evaluated by means of a tree traversal. This solution is therefore not applicable to them, since calculating subtree-specific likelihoods would increase (rather than reduce) the total number of computations performed. Nevertheless, considerable economies could be achieved by storing submatrices of **D̂**, rather than partial likelihoods, at specific nodes or branches. This would make it possible to update only those distances that changed after a given move, and reuse the submatrices corresponding to identical subtrees. However, storing submatrices at each of the *n* − 2 non-root nodes or internal branches would drastically increase the memory footprint of BLeSS analyses, which is already substantial at 10,000 tips. A more efficient approach may be to only extract the submatrix after a move picks the node or branch on which to perform a given operation. This solution would require less far-reaching changes to the RevBayes code base (only involving a re-implementation of common tree moves as overloaded functions that operate differently when applied to tree objects or distance matrix representations thereof), and may offer an ideal trade-off between CPU time and memory usage.

Although the conversion of trees to distance matrices is by far the most computationally demanding step in BLeSS inference, our results show that the burden associated with likelihood evaluation proper (i.e., computing the squared differences of the non-missing elements of the two distance matrices) also becomes considerable at large tree sizes. Even if the time spent on all other operations could be reduced to zero, it would still take ∼379 hours (over two weeks) to run an MCMC analysis of 10^6^ iterations on a dense (i.e., imputed) 10,000-by-10,000 average distance matrix (Table 3). However, as an element-wise operation, matrix subtraction is both low in computational complexity (being solvable in O(*n*^2^) time) and embarrassingly parallel (Hunold et al., 2011; Chang et al., 2023). The task can be divided into as many submatrices as there are available processors, and assuming that both the average distance matrix **D̅** and the path-length matrix **D̂** of the proposed tree can be distributed among the processors in alignment (i.e., that for all *i, j*, **D̅** *_i,j_* and **D̂** *_i,j_* are assigned to the same processor), no interprocessor communication is required between the initial distribution step and the final re-assembly of the difference matrix (Nayak, 1999; Hunold et al., 2011; Gu et al., 2015; see Östermark, 2000 for the total cost incurred by communication overhead on a 2D mesh of processors). As a result, the use of *p* processors reduces the computational cost of the problem to O(*n*^2^*/p*), potentially leading to near-linear speedup (Nayak, 1999; Hunold et al., 2011; Chang et al., 2023). Because of this feature, large matrix subtraction is particularly well-suited to graphics processing units (GPUs), which are designed to support a large number of lightweight concurrent threads (Le, 2012). Thanks to recent efforts (Smith et al., 2024), RevBayes can already exploit GPUs for calculations of standard phylogenetic likelihoods by offloading them to the high-performance BEAGLE library (Suchard and Rambaut, 2009; Ayres et al., 2012), which supports both CUDA and OpenCL platforms (Ayres et al., 2019). Extending its GPU support to the evaluation of BLeSS likelihoods would represent a fruitful avenue for increasing the efficiency of the method.

### Generalizing BLeSS to non-ultrametric trees

Throughout this study, we have assumed that both the source trees and the estimated supertree represented chronograms of extant taxa, and as such were ultrametric. However, this restriction is not inherent to the method, and a way to relax it was devised early on in the development of average consensus approaches (Lapointe and Cucumel, 1997). As a result, it became possible to estimate average consensus supertrees from source trees that were merely additive rather than ultrametric. These were originally supposed to represent phylograms, but in the context of BLeSS, they can be interpreted as serially sampled time trees, i.e., time trees with non-contemporaneous tips.

There is a growing interest in extending comparative methods that were originally developed for extant taxa to paleontological data (Slater, 2013, 2014; Bapst, 2014; Soul and Wright, 2021), but the ability to investigate clade-wide evolutionary trends is severely hampered by the lack of fossil trees comparable in size to extant megaphylogenies inferred from molecular data. Phylogenetic analyses of extinct taxa are limited in size by the costly, time-consuming, and labor-intensive nature of collecting morphological data, which have to be manually scored from the relevant specimens (often spread across many different collections worldwide) based on first-hand observations or published illustrations (Burleigh et al., 2013; Eliason et al., 2019). This problem has not yet been alleviated by new techniques such as crowdsourcing (Chang and Alfaro, 2016; O’Leary et al., 2018) or deep learning (Hunt and Pedersen, 2022; Hodel et al., 2023; Tsutsumi et al., 2023; Weaver and Smith, 2023), which largely remain at the proof-of-concept stage. As a result, even the largest morphological phylogenetic datasets usually do not include more than 200 operational taxonomic units (OTUs) (Supplementary Information, Fig. S31); the most taxon-rich character matrices of which we are aware (Hartman et al., 2019; Cau, 2024) represent clear outliers with ∼500 OTUs. Moreover, since different datasets may encode the same morphological features using different subjectively delimited characters and character states, combining them will typically require an extensive process of character redefinition and rescoring whose difficulty does not substantially differ from collecting data *de novo* (Lloyd and Slater, 2021). Accordingly, despite attempts to promote the reuse of previously assembled morphological datasets (O’Leary and Kaufman, 2011), these cannot be easily concatenated into a single taxon-rich supermatrix in a manner reminiscent of molecular alignments, and supertrees remain the only viable solution to the problem of generating comprehensive large-scale fossil phylogenies (Lloyd and Slater, 2021).

A number of comparative and macroevolutionary paleontological studies have adapted to this constraint by constructing informal (i.e., hand-drawn) supertrees and subjecting them to *post hoc* time-scaling using simple techniques based on first appearance dates (Benson et al., 2014; Famoso et al., 2016; Sakamoto et al., 2016; MacLaren, 2021; Kiat and O’Connor, 2024). This procedure has become common enough that dedicated software packages are now available to facilitate it (Castiglione et al., 2022). However, regardless of their plausibility, the resulting time trees amount to little more than expert opinions, and the process of generating them lacks the basic features of scientific workflow, including testability (since there are no underlying data to which competing expert opinions could be fitted) and reproducibility. Matrix representation with parsimony (MRP) supertrees avoid this weakness but suffer from other shortcomings, particularly the lack of a satisfying treatment of branch length information and topological uncertainty (see Introduction and Lloyd and Slater, 2021 for a recent review).

BLeSS represents an attractive alternative to both informal and MRP supertrees, and even its current implementation can considerably increase the size of phylogenies available to paleontologists. Its generalization to the non-ultrametric case would simply require replacing the simple birth-death prior used throughout this study with an alternative branching-process prior accounting for serial sampling, such as the uniform prior of Ronquist et al. (2012) or the fossilized birth-death (FBD) process (Stadler, 2010; Gavryushkina et al., 2014; Heath et al., 2014). When imputing missing entries in **D̅**, the three-point ultrametric condition (Hartigan, 1967) would further need to be replaced by the four-point additive condition (Buneman, 1974) to account for the fact that root-to-tip path lengths can no longer be assumed equal (Lapointe and Levasseur, 2004). A potential weakness of BLeSS in paleophylogenetic settings is its need for time-calibrated source trees, which may limit its applicability given that fossil phylogenies have historically been predominantly inferred using time-free methods. However, this problem is rapidly disappearing thanks to the growing popularity of Bayesian tip-dating under the FBD model (Wright et al., 2022) and availability of rigorous *post hoc* time-scaling methods such as *cal3* (Bapst, 2013). We expect BLeSS to provide a viable and powerful solution to the problem of inferring large fossil and total-evidence phylogenies, facilitating the use of information from extinct taxa in downstream analyses that currently rely mainly or exclusively on data from extant lineages.

## Acknowledgements

We are indebted to Tracy Heath for her help in clarifying the conceptual basis of the method, and to June Walker and Jöelle Barido-Sottani for their invaluable assistance with the implementation of BLeSS in `RevBayes`. We thank Michael Landis for help with simulating sequence data, Luke Kelly for advice on diagnosing MCMC convergence, and Sebastian Höhna for guidance on designing `RevBayes` validation tests. We further wish to thank Michael Foote, Andrew Magee, Fernando Meléndez-Vazquez, Erin Molloy, Jonathan Nations, C. Tomomi Parins-Fukuchi, Benjamin Redelings, Richard Ree, Ulises Rosas-Puchuri, and Nathan Upham for enlightening discussions and encouragement. We are grateful to Aaron Schein for his suggestion to use Bayesian additive regression trees to evaluate our simulation results, and to Claudio Kozický for his help with making the RevBayes implementation of BLeSS more efficient, as well as for insightful comments about possible future directions. All simulations described in this study were performed on the Midway2 and Midway3 Research Computing Clusters at the University of Chicago.

## Supplementary Material

Data available from the Dryad Digital Repository: https://doi.org/10.5061/dryad.280gb5n0w.

