## Supplementary Information for "Bayesian Least-Squares Supertrees (BLeSS): Flexible Inference of Large Time-Calibrated Phylogenies"

### Bayesian Least-Squares Supertrees (BLeSS): Fast and Accurate Inference of Large Time-Calibrated Phylogenies

David Černý[1,*] & Graham J. Slater[1]
[1] *Department of the Geophysical Sciences, University of Chicago, Chicago 60637, USA*

**2 Empirical Application to the Order Carnivora** **33**

**3 Additional Figures** **54**

**References** **55**

### 1 Simulation Tests

#### 1.1 Simulation-based calibration checking

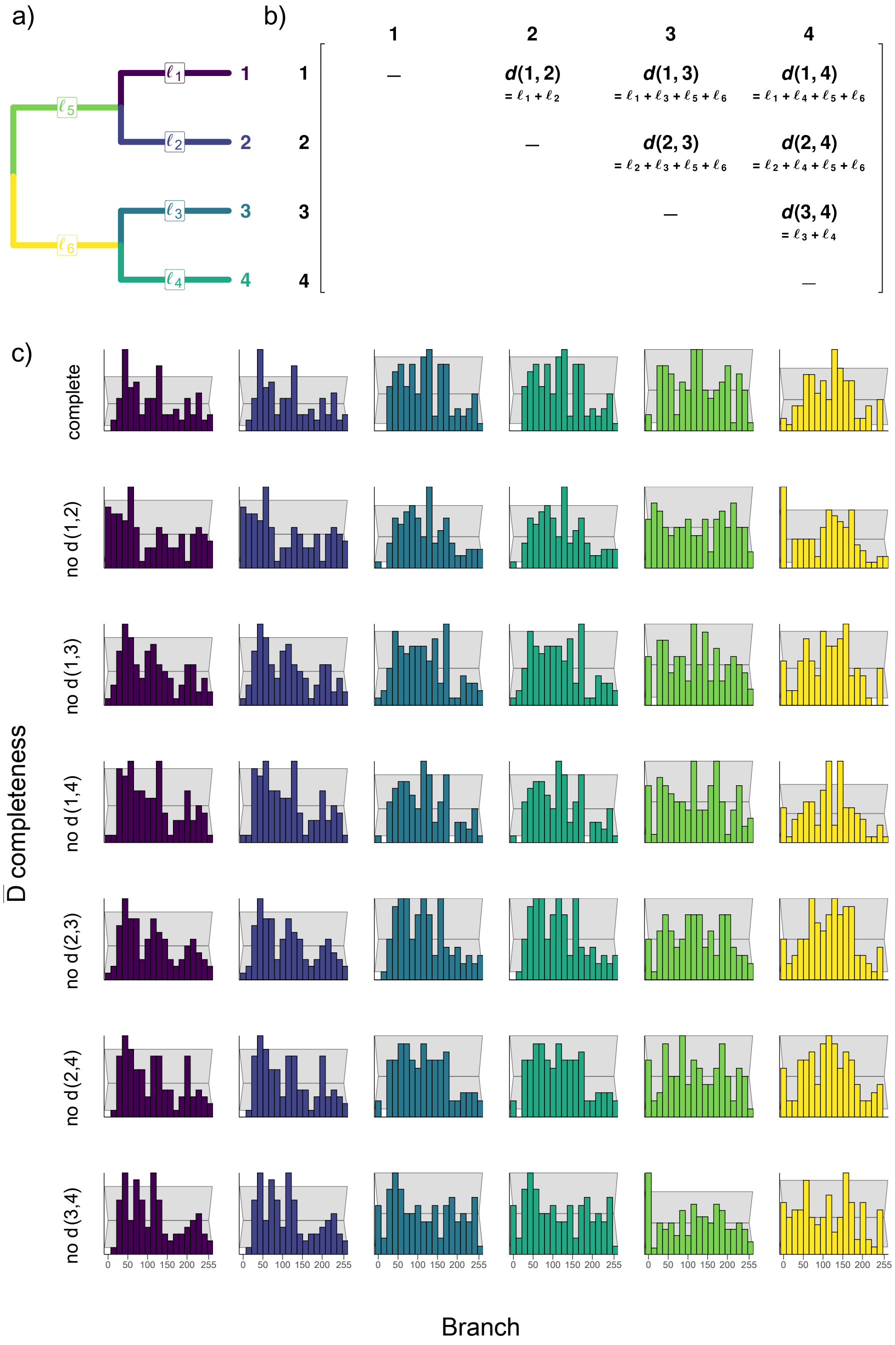

**Figure S1:** The 4-tip balanced tree with labeled branches (a) and its distance matrix, with observed distances expressed as sums of the underlying branch durations (b). Panel (c) shows the rank histograms with 95% confidence intervals (indicated by the light gray wedged rectangles in the background) for individual branch durations and different average distance matrix ($\bar{\mathbf{D}}$) completeness scenarios. See Figure 4 in the main text for the corresponding ECDF difference plots.

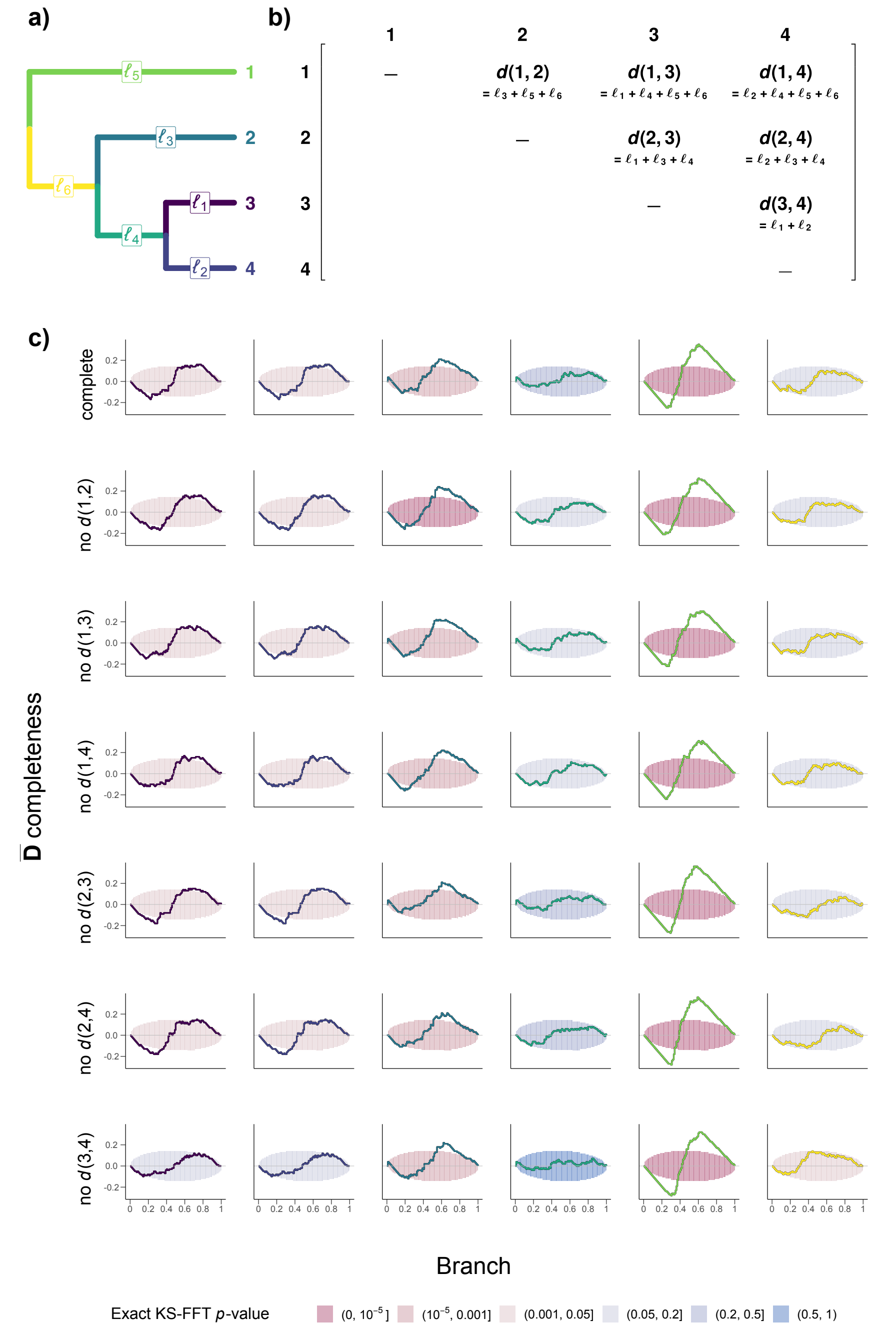

**Figure S2:** The 4-tip pectinate tree with labeled branches (a) and its distance matrix, with observed distances expressed as sums of the underlying branch durations (b). Panel (c) shows the difference between the empirical cumulative distribution function (ECDF) and the discrete uniform expectation (stepwise identity function) with 95% simultaneous confidence bands for individual branch durations and different average distance matrix ($\bar{\mathbf{D}}$) completeness scenarios. The confidence bands are colored according to the $p$-values from uniformity tests conducted using the exact Kolmogorov-Smirnov fast Fourier transform (Exact KS-FFT) method.

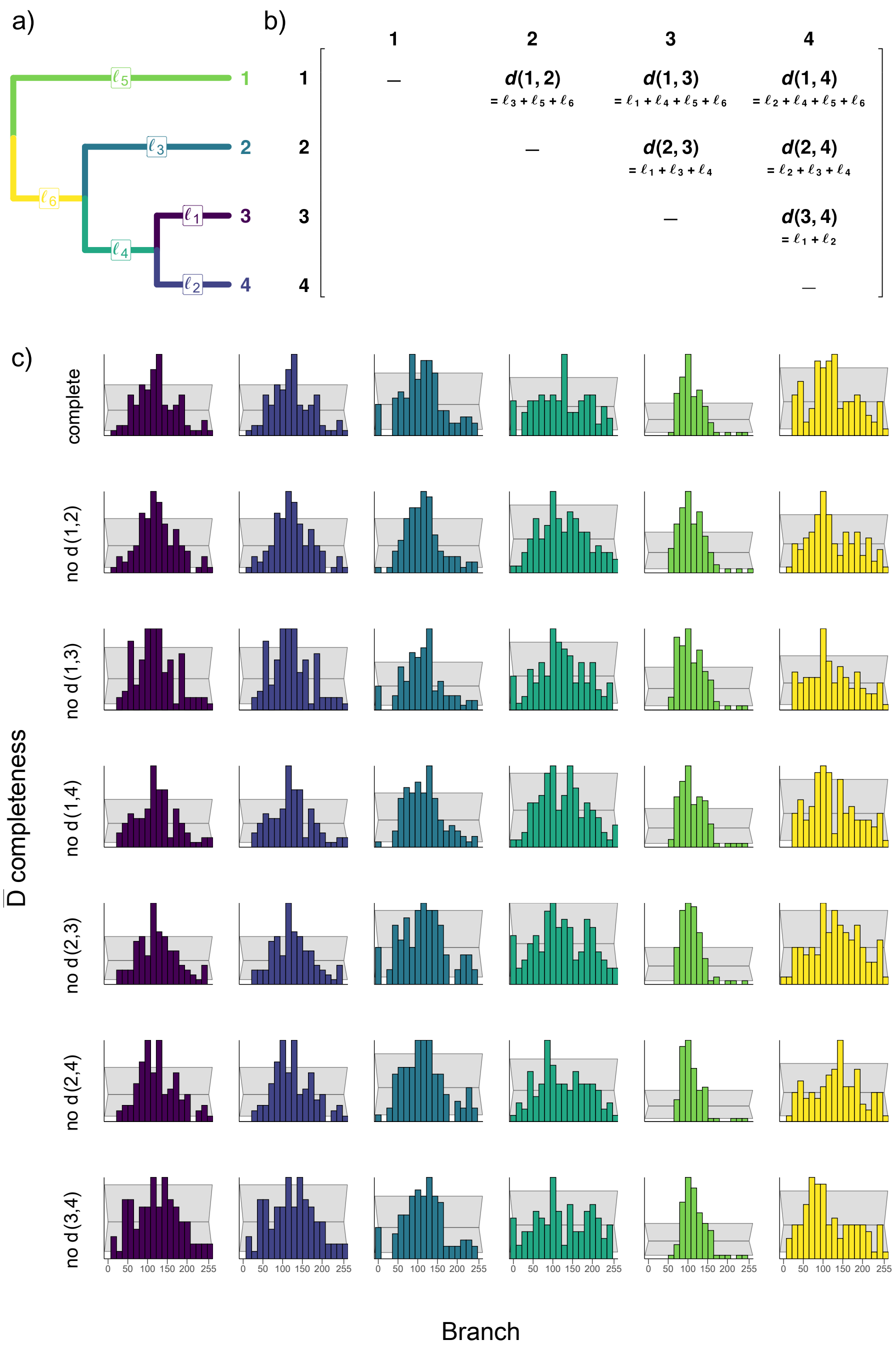

**Figure S3:** Results of simulation-based calibration checking for the 4-tip pectinate tree, visualized as rank histograms. Panel descriptions as in Figure S1.

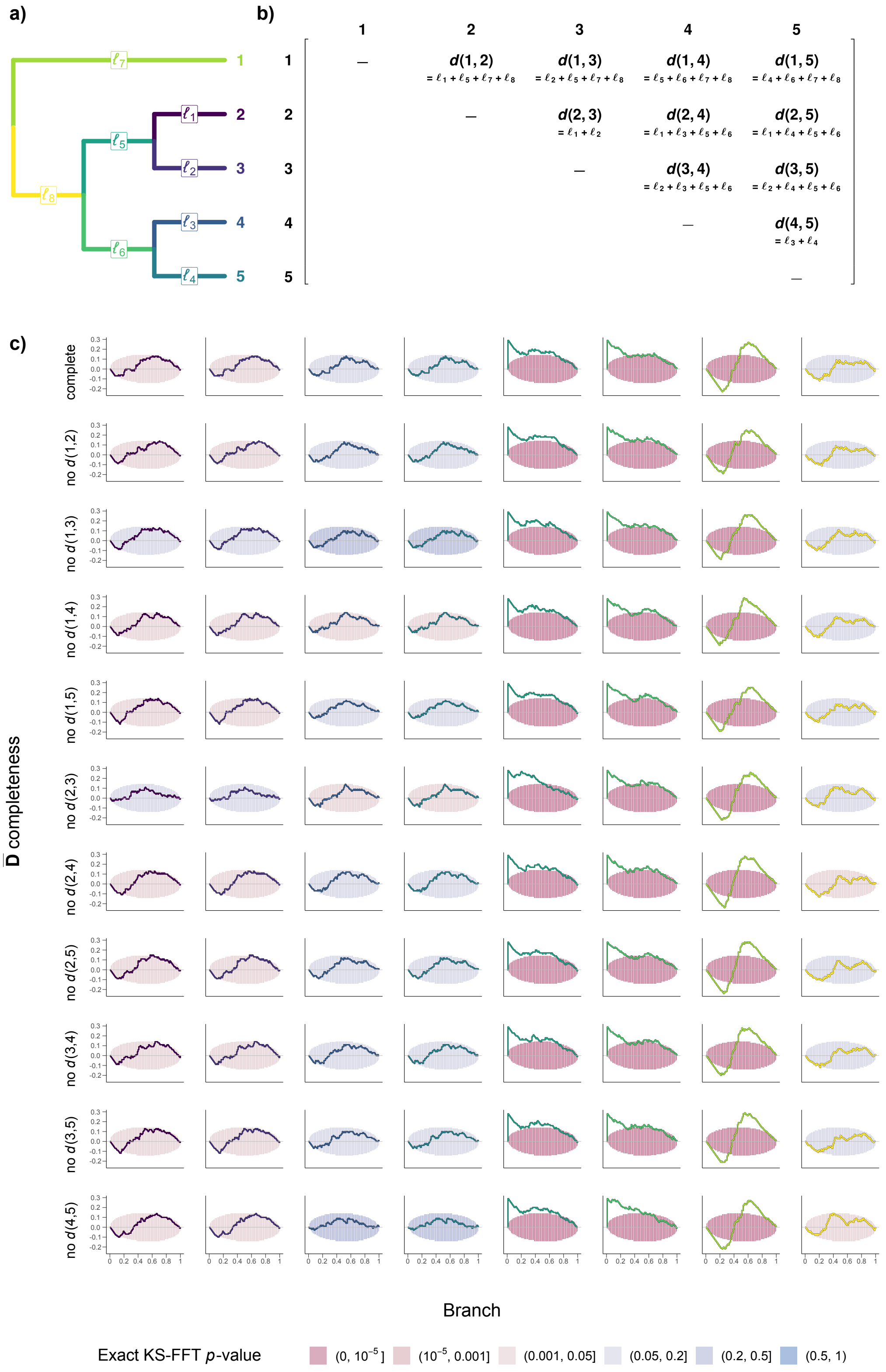

**Figure S4:** Results of simulation-based calibration checking for the 5-tip tree with a single tip sister to a balanced subtree, visualized as ECDF difference plots. Panel descriptions as in Figure S2.

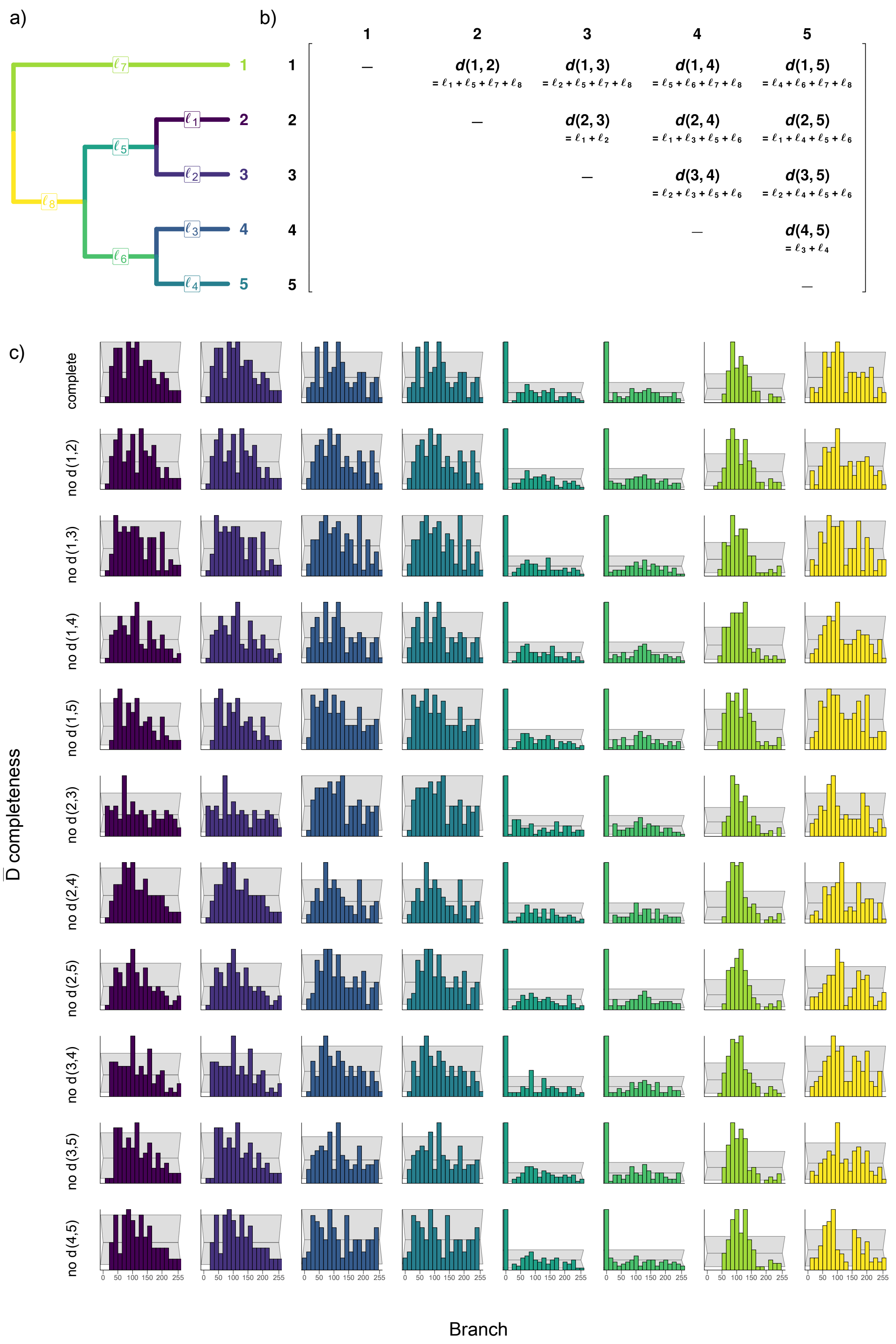

**Figure S5:** Results of simulation-based calibration checking for the 5-tip tree with a single tip sister to a balanced subtree, visualized as rank histograms. Panel descriptions as in Figure S1.

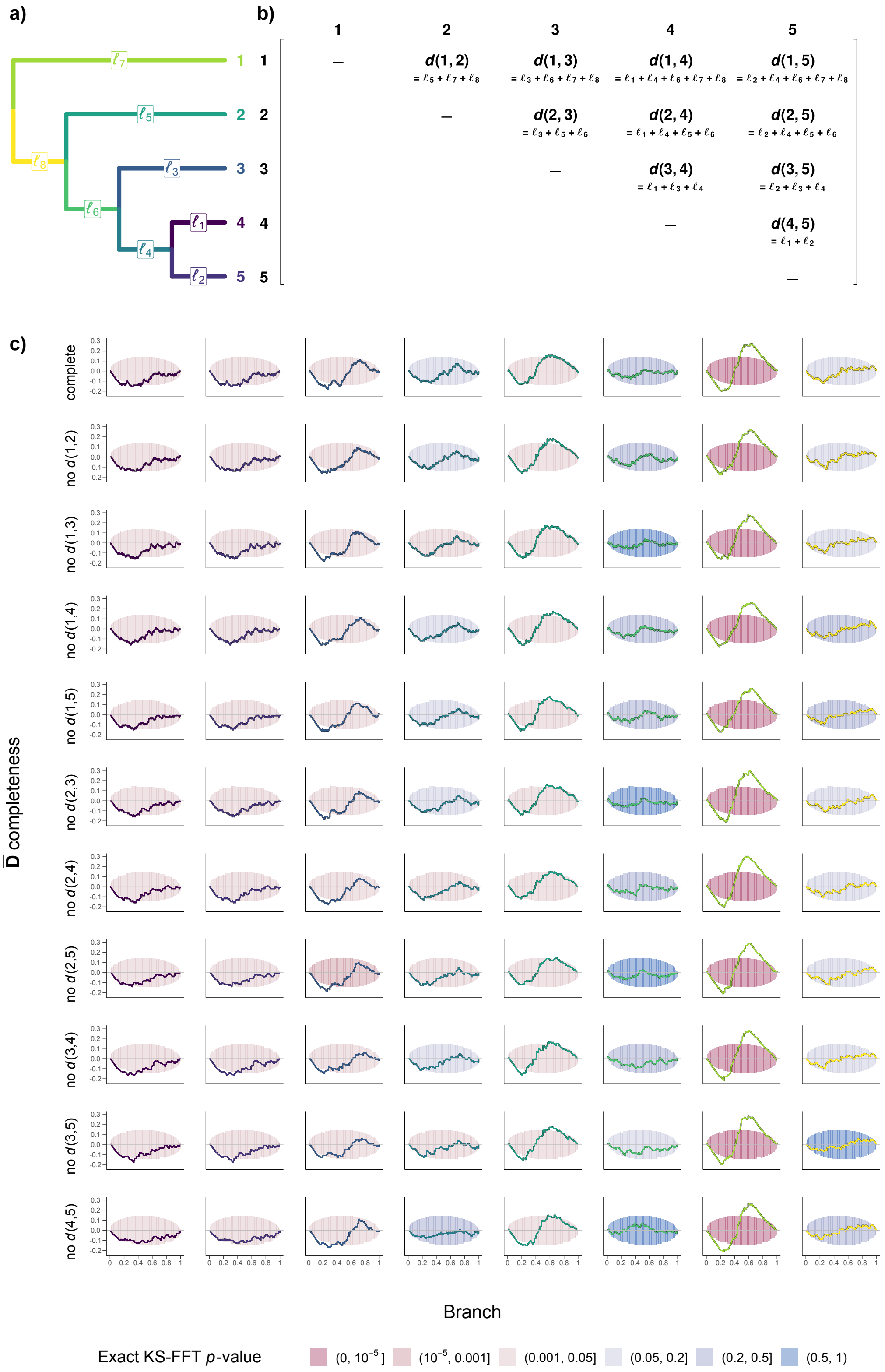

**Figure S6:** Results of simulation-based calibration checking for the fully pectinate 5-tip tree, visualized as ECDF difference plots. Panel descriptions as in Figure S2.

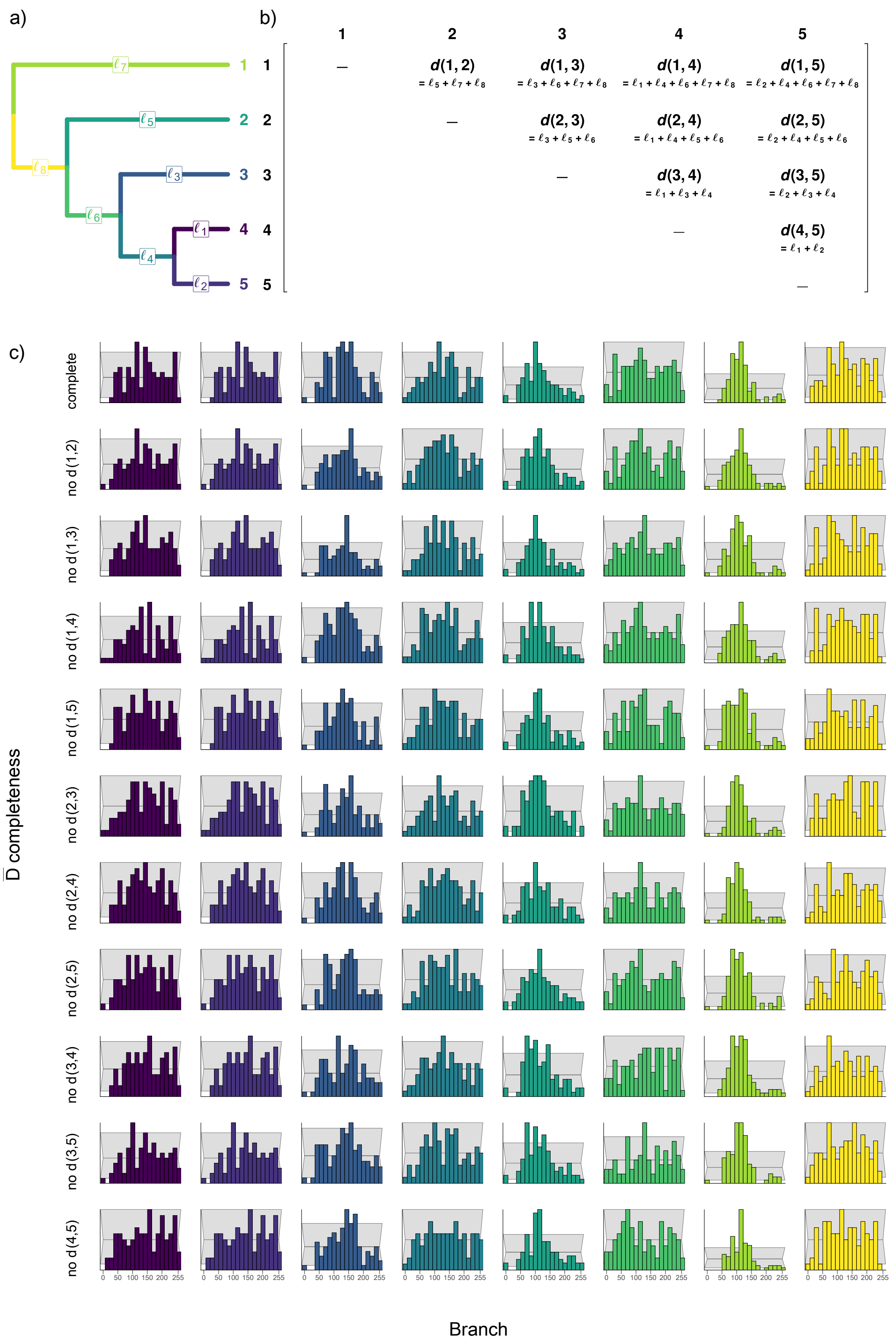

**Figure S7:** Results of simulation-based calibration checking for the fully pectinate 5-tip tree, visualized as rank histograms. Panel descriptions as in Figure S1.

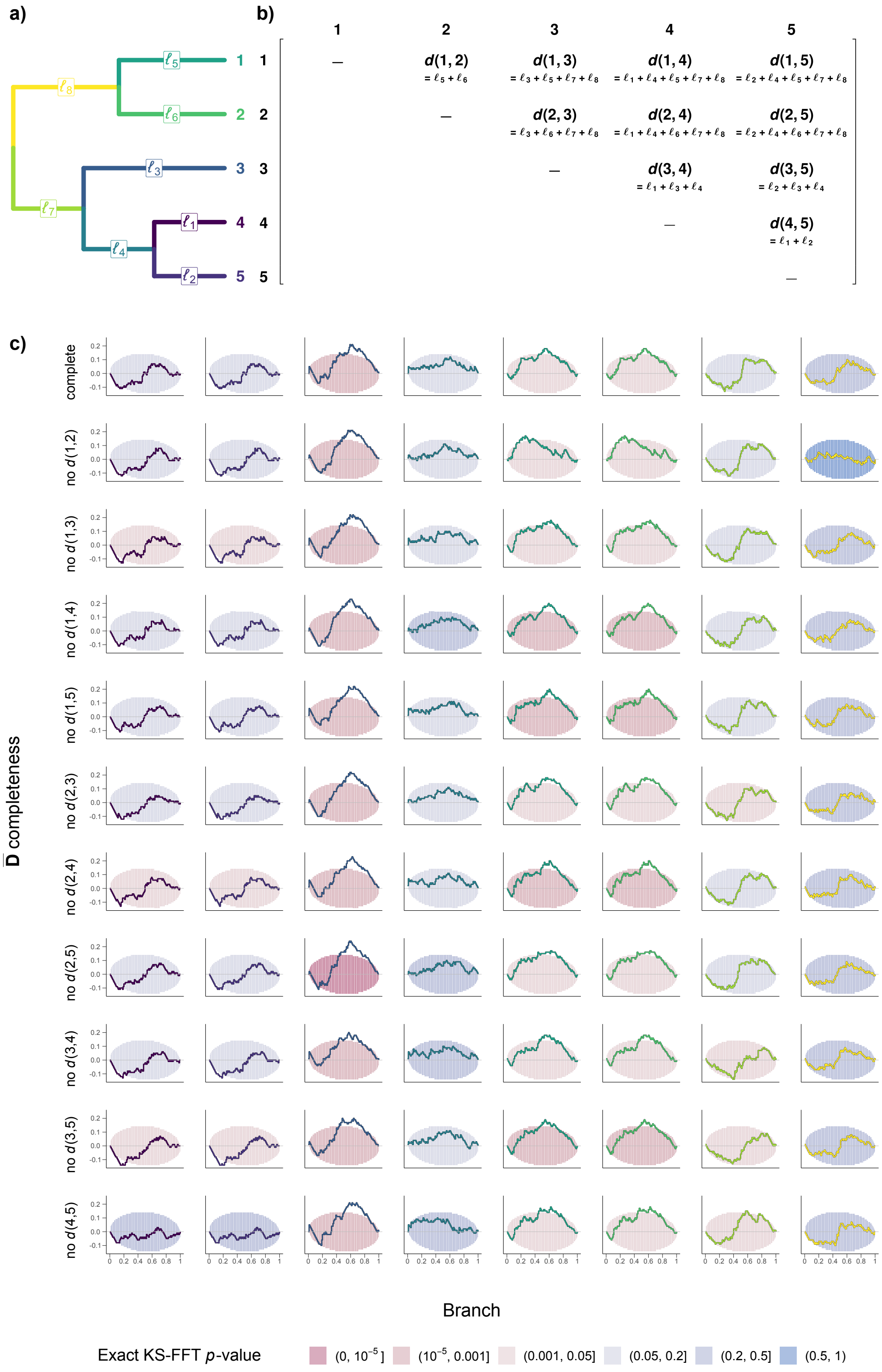

**Figure S8:** Results of simulation-based calibration checking for the 5-tip tree with a 2-tip clade sister to an imbalanced subtree, visualized as ECDF difference plots. Panel descriptions as in Figure S2.

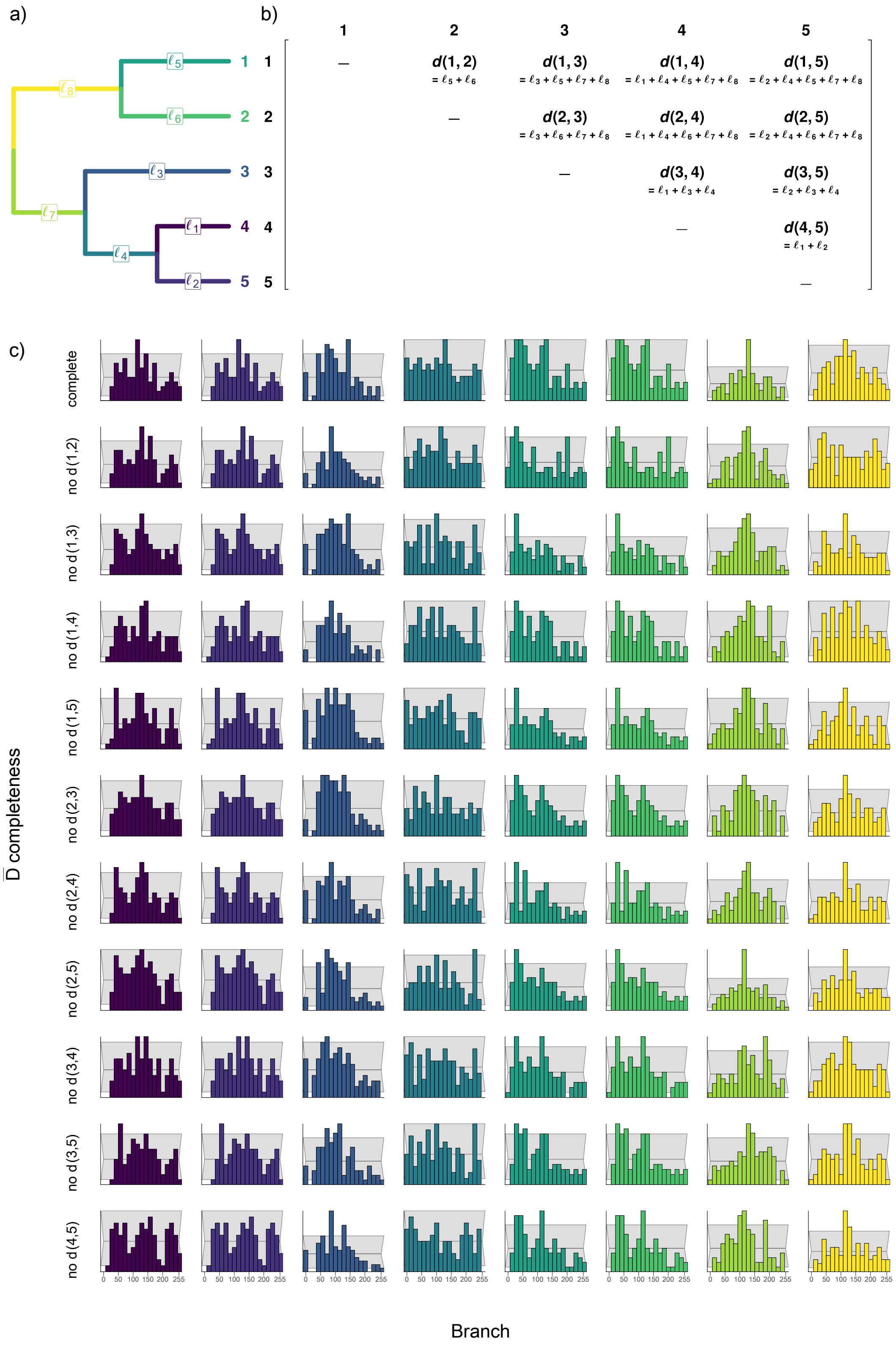

**Figure S9:** Results of simulation-based calibration checking for the 5-tip tree with a 2-tip clade sister to an imbalanced subtree, visualized as rank histograms. Panel descriptions as in Figure S1.

#### 1.2 200-tip simulations

**Table S1:** Properties of the 200-tip simulated trees and of the average distance matrices ($\bar{\mathbf{D}}$) obtained by breaking them up into subtrees under the backbone-and-patch (BP) and random selection (RS) tip sampling schemes. Root height is given in arbitrary units.

| Tree | Root height | Colless index | Subtree number (BP) | $\bar{\mathbf{D}}$ completeness (BP) | Subtree number (RS) | $\bar{\mathbf{D}}$ completeness (RS) |
|---|---|---|---|---|---|---|
| 1 | 191.66 | 0.0324 | 4 | 0.3577 | 18 | 0.5172 |
| 2 | 146.87 | 0.0521 | 9 | 0.3972 | 28 | 0.6863 |
| 3 | 11.68 | 0.0306 | 9 | 0.1958 | 24 | 0.6992 |
| 4 | 67.68 | 0.0586 | 4 | 0.5046 | 28 | 0.6763 |
| 5 | 193.82 | 0.0453 | 8 | 0.375 | 36 | 0.7446 |
| 6 | 128.73 | 0.0333 | 8 | 0.1614 | 32 | 0.7561 |
| 7 | 47.7 | 0.0404 | 27 | 0.297 | 23 | 0.6455 |
| 8 | 10.06 | 0.0283 | 32 | 0.1119 | 32 | 0.7732 |
| 9 | 95.48 | 0.0548 | 7 | 0.38 | 31 | 0.6664 |
| 10 | 157.06 | 0.0423 | 12 | 0.1402 | 25 | 0.6581 |
| 11 | 104.62 | 0.0337 | 4 | 0.3703 | 29 | 0.6789 |
| 12 | 64.79 | 0.045 | 8 | 0.2582 | 27 | 0.7264 |
| 13 | 131.73 | 0.041 | 10 | 0.1516 | 33 | 0.8163 |
| 14 | 180.26 | 0.0389 | 4 | 0.3666 | 26 | 0.6462 |
| 15 | 5.94 | 0.0385 | 9 | 0.1666 | 21 | 0.6196 |
| 16 | 59.27 | 0.0475 | 15 | 0.0969 | 30 | 0.703 |
| 17 | 43.63 | 0.0457 | 18 | 0.1009 | 28 | 0.6227 |
| 18 | 105.94 | 0.0325 | 7 | 0.2483 | 30 | 0.6952 |
| 19 | 113.23 | 0.0343 | 4 | 0.4293 | 29 | 0.7608 |
| 20 | 56 | 0.0373 | 12 | 0.1674 | 28 | 0.7493 |
| 21 | 157.67 | 0.0358 | 8 | 0.2257 | 17 | 0.5768 |
| 22 | 133.89 | 0.0338 | 4 | 0.3354 | 26 | 0.6963 |
| 23 | 172.42 | 0.0393 | 4 | 0.4478 | 35 | 0.7478 |
| 24 | 187.71 | 0.0358 | 4 | 0.4626 | 23 | 0.6354 |
| 25 | 126.88 | 0.0545 | 4 | 0.6037 | 25 | 0.651 |
| 26 | 195.83 | 0.0663 | 4 | 0.59 | 19 | 0.6432 |
| 27 | 197.24 | 0.0397 | 8 | 0.261 | 23 | 0.5939 |
| 28 | 64.9 | 0.0456 | 10 | 0.2152 | 49 | 0.8902 |
| 29 | 2.1 | 0.0338 | 4 | 0.3748 | 24 | 0.6284 |
| 30 | 65 | 0.0588 | 11 | 0.1437 | 24 | 0.6086 |
| 31 | 166.48 | 0.0399 | 4 | 0.4546 | 16 | 0.6012 |
| 32 | 98.35 | 0.0378 | 4 | 0.7575 | 27 | 0.6368 |
| 33 | 86 | 0.0436 | 7 | 0.2862 | 25 | 0.6538 |
| 34 | 197.84 | 0.0352 | 4 | 0.3558 | 17 | 0.6776 |
| 35 | 189.31 | 0.0504 | 7 | 0.2372 | 27 | 0.6922 |
| 36 | 84.75 | 0.0585 | 8 | 0.3275 | 32 | 0.7789 |
| 37 | 142.13 | 0.0357 | 4 | 0.5095 | 31 | 0.7152 |
| 38 | 196.95 | 0.0442 | 5 | 0.3114 | 24 | 0.6911 |
| 39 | 58.75 | 0.0638 | 7 | 0.3404 | 23 | 0.5986 |
| 40 | 50.14 | 0.0442 | 10 | 0.1794 | 35 | 0.8174 |
| 41 | 190.81 | 0.0325 | 4 | 0.3439 | 25 | 0.6463 |
| 42 | 27.57 | 0.0385 | 7 | 0.2193 | 20 | 0.5936 |
| 43 | 44.03 | 0.046 | 5 | 0.476 | 29 | 0.7102 |
| 44 | 181.96 | 0.0508 | 4 | 0.4982 | 16 | 0.5988 |
| 45 | 45.42 | 0.0378 | 8 | 0.195 | 22 | 0.6847 |

| | | | | | | |
|---|---|---|---|---|---|---|
| 46 | 47.41 | 0.0593 | 8 | 0.2322 | 22 | 0.5376 |
| 47 | 178.36 | 0.0375 | 6 | 0.2498 | 26 | 0.7404 |
| 48 | 154.12 | 0.0371 | 4 | 0.4703 | 23 | 0.5978 |
| 49 | 134.95 | 0.0497 | 20 | 0.0877 | 28 | 0.7398 |
| 50 | 22.96 | 0.0269 | 4 | 0.3546 | 21 | 0.6536 |
| 51 | 29.12 | 0.0406 | 11 | 0.1568 | 25 | 0.6989 |
| 52 | 54.08 | 0.0456 | 8 | 0.2628 | 29 | 0.7327 |
| 53 | 161.31 | 0.0387 | 9 | 0.1953 | 25 | 0.7458 |
| 54 | 153.85 | 0.0523 | 4 | 0.7781 | 40 | 0.8354 |
| 55 | 50.26 | 0.0441 | 4 | 0.4357 | 38 | 0.847 |
| 56 | 129.34 | 0.0411 | 31 | 0.0908 | 38 | 0.8336 |
| 57 | 69.06 | 0.0387 | 6 | 0.391 | 28 | 0.6944 |
| 58 | 86.19 | 0.0333 | 13 | 0.1057 | 20 | 0.6369 |
| 59 | 161.55 | 0.0467 | 4 | 0.4137 | 29 | 0.7272 |
| 60 | 105.57 | 0.0421 | 7 | 0.2388 | 20 | 0.642 |
| 61 | 128.38 | 0.0479 | 5 | 0.4062 | 25 | 0.7028 |
| 62 | 88.76 | 0.0518 | 10 | 0.1624 | 22 | 0.607 |
| 63 | 178.6 | 0.0367 | 6 | 0.266 | 20 | 0.6359 |
| 64 | 127.09 | 0.0397 | 15 | 0.0989 | 34 | 0.8026 |
| 65 | 62.6 | 0.0354 | 10 | 0.1388 | 25 | 0.6769 |
| 66 | 71.45 | 0.0505 | 12 | 0.1719 | 19 | 0.5826 |
| 67 | 124.11 | 0.0331 | 5 | 0.2652 | 39 | 0.8078 |
| 68 | 58.03 | 0.0385 | 10 | 0.1706 | 23 | 0.5929 |
| 69 | 83.93 | 0.0395 | 6 | 0.2462 | 29 | 0.7422 |
| 70 | 131.84 | 0.0568 | 7 | 0.3083 | 25 | 0.7079 |
| 71 | 188.87 | 0.0475 | 9 | 0.1832 | 38 | 0.7819 |
| 72 | 187.32 | 0.0374 | 7 | 0.2492 | 27 | 0.6246 |
| 73 | 159.59 | 0.0384 | 7 | 0.2052 | 25 | 0.7172 |
| 74 | 25.25 | 0.0407 | 7 | 0.2021 | 27 | 0.6826 |
| 75 | 69.58 | 0.0447 | 4 | 0.6073 | 24 | 0.6563 |
| 76 | 83.82 | 0.0497 | 5 | 0.4503 | 29 | 0.6496 |
| 77 | 67.01 | 0.0472 | 4 | 0.4024 | 27 | 0.7412 |
| 78 | 153.93 | 0.0381 | 7 | 0.241 | 18 | 0.5868 |
| 79 | 43.29 | 0.0349 | 9 | 0.1941 | 34 | 0.7364 |
| 80 | 29.3 | 0.0622 | 4 | 0.6328 | 22 | 0.6323 |
| 81 | 119.91 | 0.0548 | 10 | 0.1846 | 22 | 0.64 |
| 82 | 45.55 | 0.038 | 6 | 0.3384 | 19 | 0.592 |
| 83 | 111.18 | 0.0388 | 12 | 0.1696 | 27 | 0.741 |
| 84 | 185.11 | 0.0354 | 8 | 0.1871 | 22 | 0.6239 |
| 85 | 109.34 | 0.0522 | 4 | 0.3822 | 25 | 0.6541 |
| 86 | 105.81 | 0.0421 | 8 | 0.1864 | 25 | 0.74 |
| 87 | 162.96 | 0.0608 | 5 | 0.4956 | 36 | 0.7817 |
| 88 | 162.29 | 0.032 | 8 | 0.2058 | 26 | 0.6335 |
| 89 | 185.93 | 0.035 | 6 | 0.2757 | 28 | 0.6888 |
| 90 | 14.24 | 0.0363 | 4 | 0.4427 | 29 | 0.6824 |
| 91 | 135.74 | 0.0356 | 5 | 0.339 | 34 | 0.7338 |
| 92 | 172.67 | 0.0517 | 4 | 0.5664 | 18 | 0.5885 |
| 93 | 176.52 | 0.0368 | 5 | 0.3308 | 22 | 0.5901 |
| 94 | 174.7 | 0.0346 | 7 | 0.2299 | 26 | 0.6276 |
| 95 | 105.21 | 0.0481 | 4 | 0.412 | 26 | 0.6704 |
| 96 | 125.04 | 0.0388 | 4 | 0.4562 | 42 | 0.8293 |
| 97 | 53.68 | 0.0388 | 5 | 0.4122 | 24 | 0.655 |
| 98 | 30.97 | 0.0486 | 7 | 0.3959 | 27 | 0.6348 |
| 99 | 140.2 | 0.0356 | 4 | 0.4786 | 27 | 0.7048 |
| 100 | 32.01 | 0.0465 | 25 | 0.0692 | 29 | 0.7518 |

**Table S2:** Topological and divergence time accuracy of the maximum *a posteriori* (MAP) and maximum clade credibility (MCC) summary supertrees inferred under $\lambda_e = 0.1$. For each of the $n$ replicates that reached convergence, the posterior summary was compared to the corresponding true tree in terms of Robinson-Foulds (RF) distance and normalized root age difference ($\Delta$ root age = $|a_e - a_t| \; / \; a_t$, where $a_t$ is the true root age and $a_e$ is the posterior mean). Every statistic is reported as the mean ± standard deviation over all $n$ replicates.

| | | MAP tree | | MCC tree | |
|---|---|---|---|---|---|
| **Scenario** | ***n*** | **RF dist.** | **Δ root age** | **RF dist.** | **Δ root age** |
| no error, backbone and patch | 21 | 0.4643 ± 0.1798 | 0.1970 ± 0.1314 | 0.4462 ± 0.1856 | 0.1970 ± 0.1314 |
| no error, random selection | 87 | 0.2166 ± 0.1181 | 0.0004 ± 0.0032 | 0.1718 ± 0.1216 | 0.0004 ± 0.0032 |
| no error, backbone and patch, imputed | 98 | 0.1141 ± 0.1066 | 0.0003 ± 0.0025 | 0.0709 ± 0.1054 | 0.0003 ± 0.0025 |
| no error, random selection, imputed | 98 | 0.2454 ± 0.0844 | 0 ± 0.0001 | 0.2188 ± 0.0819 | 0 ± 0.0001 |
| top error, backbone and patch | 31 | 0.5618 ± 0.1410 | 1.1117 ± 3.0300 | 0.5359 ± 0.1440 | 1.1117 ± 3.0300 |
| top error, random selection | 43 | 0.2624 ± 0.0795 | 0.5964 ± 1.4833 | 0.2347 ± 0.0911 | 0.5964 ± 1.4833 |
| top error, backbone and patch, imputed | 45 | 0.7533 ± 0.0641 | 0.6819 ± 1.0366 | 0.7489 ± 0.0623 | 0.6819 ± 1.0366 |
| top error, random selection, imputed | 97 | 0.9651 ± 0.0159 | 0.6516 ± 1.6782 | 0.9645 ± 0.0159 | 0.6516 ± 1.6782 |

**Table S3:** Topological and divergence time accuracy of the MAP and MCC summary supertrees inferred under $\lambda_e = 0.33$. Abbreviations as in Table S2.

| | | MAP tree | | MCC tree | |
|---|---|---|---|---|---|
| **Scenario** | ***n*** | **RF dist.** | **Δ root age** | **RF dist.** | **Δ root age** |
| no error, backbone and patch | 13 | 0.4205 ± 0.2046 | 0.1508 ± 0.1434 | 0.4034 ± 0.2046 | 0.1508 ± 0.1434 |
| no error, random selection | 67 | 0.1590 ± 0.0664 | 0 ± 0 | 0.1278 ± 0.0573 | 0 ± 0 |
| no error, backbone and patch, imputed | 95 | 0.0690 ± 0.0741 | 0 ± 0.0001 | 0.0349 ± 0.0685 | 0 ± 0.0001 |
| no error, random selection, imputed | 92 | 0.2240 ± 0.0914 | 0 ± 0.0001 | 0.1993 ± 0.0939 | 0 ± 0.0001 |
| top error, backbone and patch | 29 | 0.5351 ± 0.1425 | 1.0427 ± 2.9626 | 0.5137 ± 0.1417 | 1.0427 ± 2.9626 |
| top error, random selection | 42 | 0.2499 ± 0.1248 | 1.2046 ± 3.3431 | 0.2350 ± 0.1297 | 1.2046 ± 3.3431 |
| top error, backbone and patch, imputed | 41 | 0.7558 ± 0.0526 | 0.7407 ± 1.0791 | 0.7555 ± 0.0525 | 0.7407 ± 1.0791 |
| top error, random selection, imputed | 97 | 0.9655 ± 0.0162 | 0.6484 ± 1.6782 | 0.9654 ± 0.0159 | 0.6484 ± 1.6782 |

**Table S4:** Topological and divergence time accuracy of the MAP and MCC summary supertrees inferred under $\lambda_e = 1$. Abbreviations as in Table S2.

| | | MAP tree | | MCC tree | |
|---|---|---|---|---|---|
| **Scenario** | ***n*** | **RF dist.** | **Δ root age** | **RF dist.** | **Δ root age** |
| no error, backbone and patch | 7 | 0.3561 ± 0.1366 | 0.0611 ± 0.0760 | 0.3387 ± 0.1450 | 0.0611 ± 0.0760 |
| no error, random selection | 75 | 0.1340 ± 0.0779 | 0 ± 0 | 0.1076 ± 0.0752 | 0 ± 0 |
| no error, backbone and patch, imputed | 86 | 0.0301 ± 0.0278 | 0 ± 0 | 0.0116 ± 0.0169 | 0 ± 0 |
| no error, random selection, imputed | 77 | 0.1993 ± 0.0844 | 0 ± 0 | 0.1775 ± 0.0803 | 0 ± 0 |
| top error, backbone and patch | 26 | 0.5203 ± 0.1348 | 1.2330 ± 3.1857 | 0.5027 ± 0.1312 | 1.2330 ± 3.1857 |
| top error, random selection | 28 | 0.2808 ± 0.1096 | 1.1226 ± 1.9651 | 0.2745 ± 0.1098 | 1.1226 ± 1.9651 |
| top error, backbone and patch, imputed | 40 | 0.7490 ± 0.0547 | 0.7205 ± 1.0890 | 0.7490 ± 0.0547 | 0.7205 ± 1.0890 |
| top error, random selection, imputed | 97 | 0.9651 ± 0.0161 | 0.6511 ± 1.6788 | 0.9653 ± 0.0160 | 0.6511 ± 1.6788 |

**Table S5:** Accuracy of the MCC summary supertrees inferred under $\lambda_e = 0.1$. For each of the $n$ replicates that reached convergence, the posterior summary was compared to the corresponding true tree in terms of Kuhner-Felsenstein (KF), Billera-Holmes-Vogtmann (BHV), and matching split distances. Every statistic is reported as the mean and 95% interpercentile range over all $n$ replicates.

| Scenario | $n$ | KF distance | BHV distance | Matching split dist. |
|---|---|---|---|---|
| no error, backbone and patch | 21 | 175.7 [10.25, 276.29] | 222.14 [12.22, 356.5] | 978.38 [263.5, 1839.5] |
| no error, random selection | 87 | 74.6 [21.01, 163.07] | 83.91 [23.2, 182.42] | 113.06 [35.45, 320.5] |
| no error, backbone and patch, imputed | 98 | 7.06 [5.57, 11.65] | 7.61 [5.92, 12.99] | 62 [10, 213.35] |
| no error, random selection, imputed | 98 | 165.34 [35.28, 387.57] | 178.4 [37.79, 440.19] | 154.92 [53.27, 312.2] |
| top error, backbone and patch | 31 | 240.16 [136.06, 344.01] | 296 [147.11, 448.14] | 1180.87 [477.5, 1806] |
| top error, random selection | 43 | 156.69 [65.56, 251.32] | 168.22 [71.35, 280.02] | 247.53 [90.6, 640.5] |
| top error, backbone and patch, imputed | 45 | 381.99 [193.69, 508.57] | 460.11 [210.62, 654.69] | 869.31 [669.4, 1158.1] |
| top error, random selection, imputed | 97 | 448.5 [274.35, 592.01] | 531.88 [286.96, 727.7] | 1394.81 [1028.2, 1961.8] |

**Table S6:** Accuracy of the MCC summary supertrees inferred under $\lambda_e = 0.33$. Abbreviations as in Table S5.

| Scenario | $n$ | KF distance | BHV distance | Matching split dist. |
|---|---|---|---|---|
| no error, backbone and patch | 13 | 153.14, [25.92, 257.11] | 191.57, [32.28, 329.21] | 880.23, [108, 1549.5] |
| no error, random selection | 67 | 72.91, [19.9, 157.58] | 80.62, [21.96, 176.86] | 73.57, [19, 173.1] |
| no error, backbone and patch, imputed | 95 | 3.01, [2.24, 4.35] | 3.18, [2.36, 4.76] | 27.92, [0.7, 104.3] |
| no error, random selection, imputed | 92 | 165.09, [25.06, 391.58] | 178.8, [27, 432.65] | 141.77, [40.83, 299.25] |
| top error, backbone and patch | 29 | 236.24, [138.59, 346.89] | 291.95, [148.16, 441.64] | 1158.76, [579, 1821.8] |
| top error, random selection | 42 | 156.53, [65.42, 246.92] | 165.38, [70.9, 267.03] | 277.12, [67.88, 820.73] |
| top error, backbone and patch, imputed | 41 | 377.32, [191.47, 519.57] | 454.93, [208.67, 661.57] | 870.39, [685, 1078] |
| top error, random selection, imputed | 97 | 449.84, [265.72, 612.32] | 533.64, [278.85, 738.85] | 1414.72, [1030.2, 2068] |

**Table S7:** Accuracy of the MCC summary supertrees inferred under $\lambda_e = 1$. Abbreviations as in Table S5.

| Scenario | $n$ | KF distance | BHV distance | Matching split dist. |
|---|---|---|---|---|
| no error, backbone and patch | 7 | 121.72, [19.87, 235.45] | 156.84, [24.98, 301.41] | 790.29, [217.85, 1353.2] |
| no error, random selection | 75 | 63.94, [16.04, 135.94] | 71.1, [17.43, 150.62] | 58.97, [11.95, 210.25] |
| no error, backbone and patch, imputed | 86 | 1.3, [0.96, 1.8] | 1.34, [0.96, 2.01] | 11.17, [0, 96.25] |
| no error, random selection, imputed | 77 | 145.22, [21.82, 295.24] | 155.9, [23.01, 324.07] | 125.47, [28.6, 277] |
| top error, backbone and patch | 26 | 229.44, [138.34, 337.68] | 276.1, [145.44, 420.67] | 1178.08, [531.25, 1819] |
| top error, random selection | 28 | 189.18, [116.11, 263.13] | 199.53, [121.3, 273.98] | 325.04, [135.4, 704.52] |
| top error, backbone and patch, imputed | 40 | 378.64, [191.23, 509.49] | 454.9, [208.13, 655.52] | 864.72, [676.78, 1082.88] |
| top error, random selection, imputed | 97 | 449.35, [271.36, 590.53] | 532.95, [279.87, 726.98] | 1418.04, [1028.4, 2087.4] |

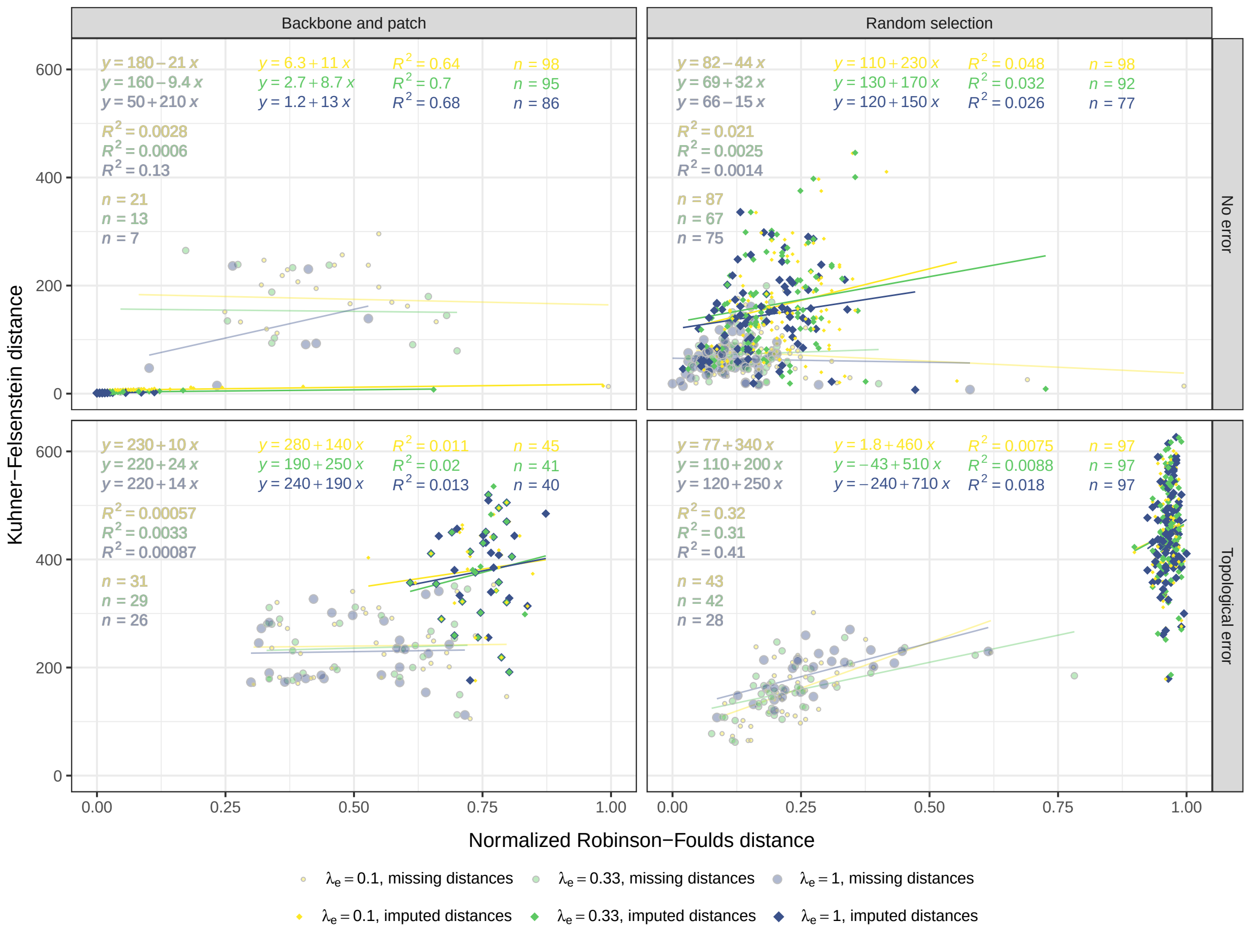

**Figure S10:** Scenario-specific relationships of the Robinson-Foulds ($x$) and Kuhner-Felsenstein ($y$) distances between the true (simulated) and estimated (BLeSS MCC) supertrees, computed from all $n$ replicates that reached convergence. These two accuracy metrics were used as response variables in the Bayesian additive regression tree (BART) analyses of the 200-tip simulation results. While exhibiting a strong and positive correlation globally (i.e., across scenarios: $y = 28 + 430x$; $R^2 = 0.8$), they show no or even an inverse relationship in several simulation scenarios (e.g., no error, random selection, missing distances), motivating the use of both metrics.

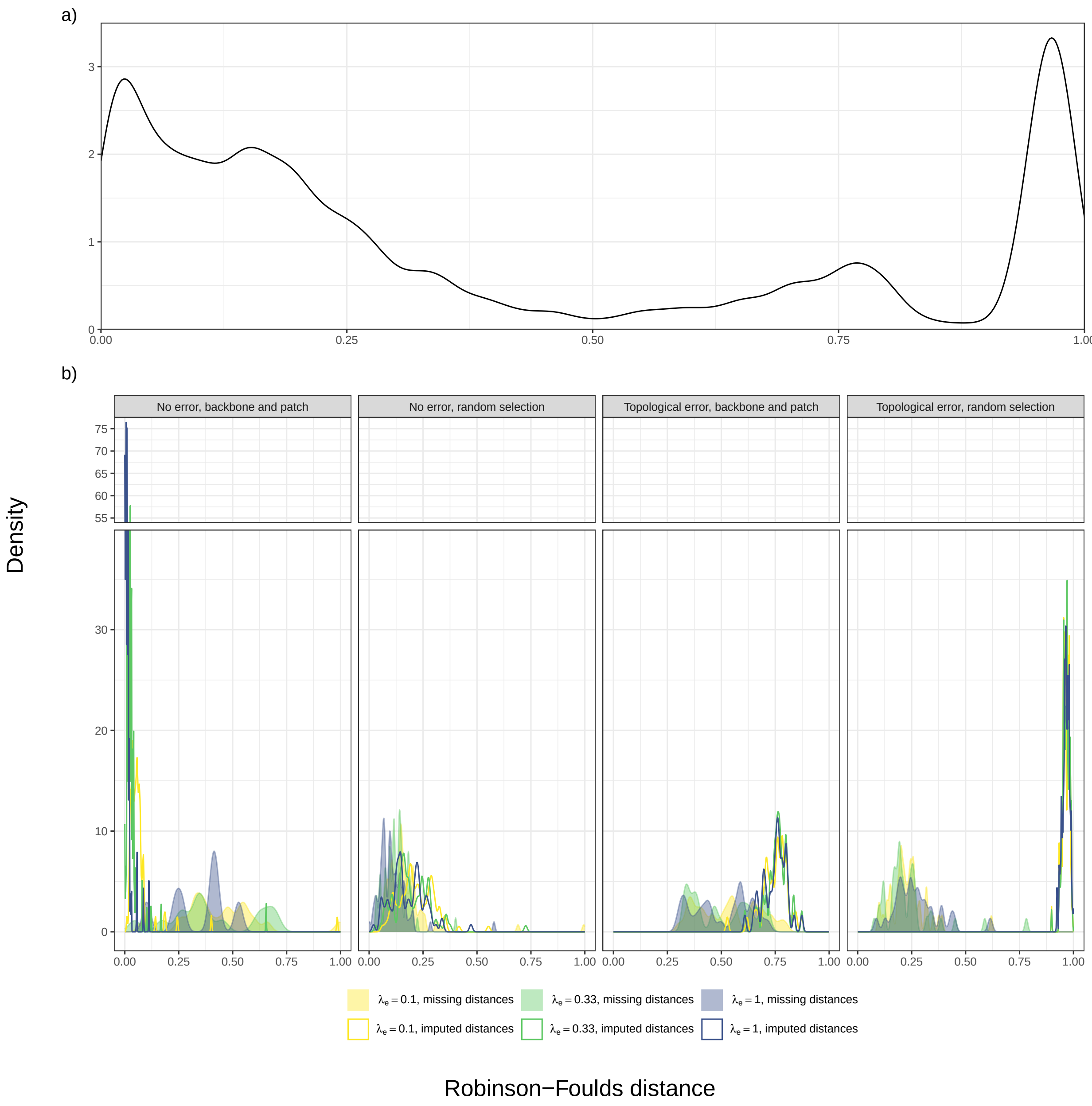

**Figure S11:** Global (a) and scenario-specific (b) distributions of the Robinson-Foulds distance between the true (simulated) and estimated (BLeSS MCC) supertree, one of the accuracy metrics used as response variables in the BART analyses of the 200-tip simulation results. Note the $y$-axis break from 38 to 55 in panel (b).

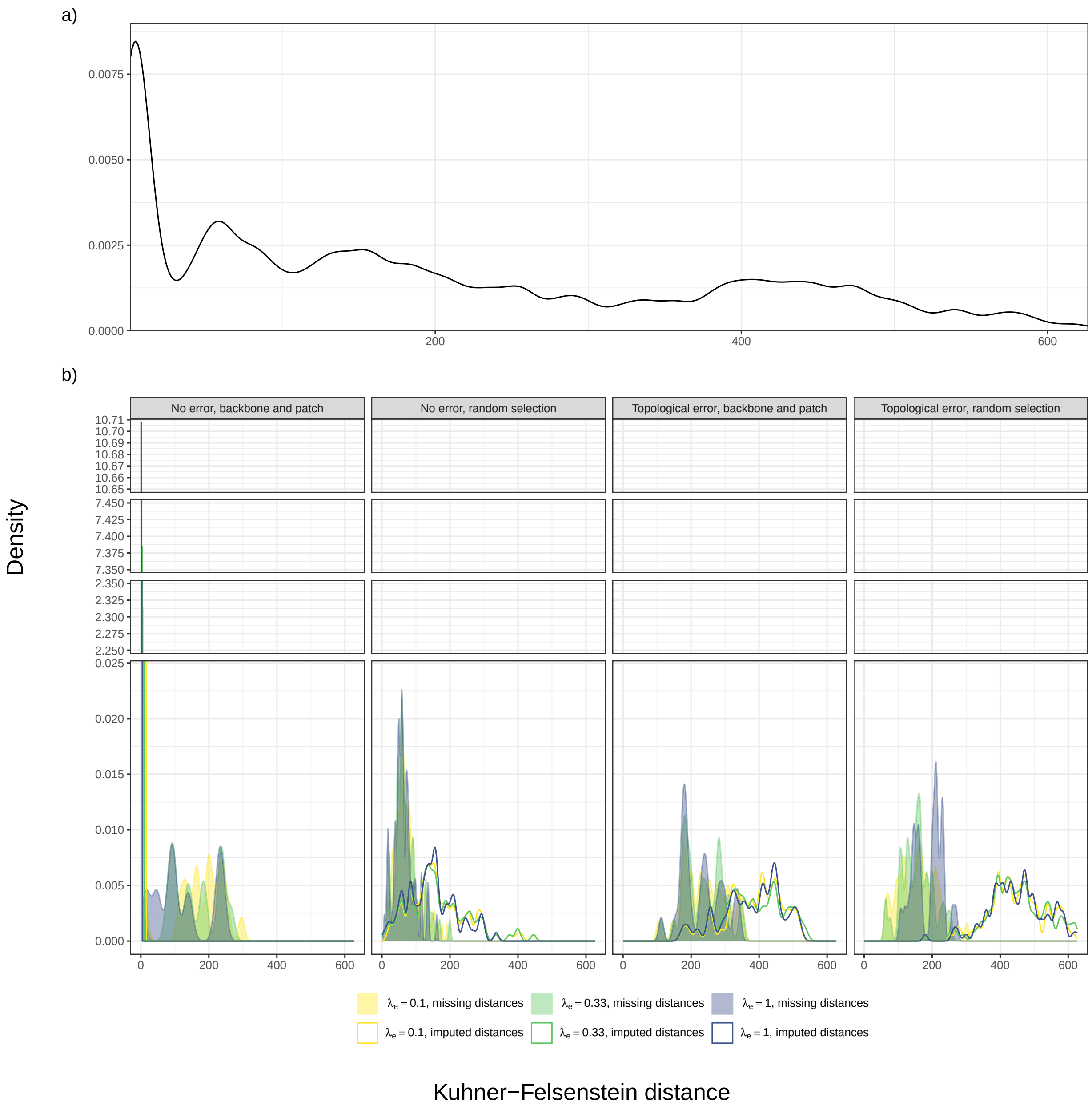

**Figure S12:** Global (a) and scenario-specific (b) distributions of the Kuhner-Felsenstein distance between the true (simulated) and estimated (BLeSS MCC) supertree, one of the accuracy metrics used as response variables in the BART analyses of the 200-tip simulation results. Note the $y$-axis breaks from 0.024 to 2.25, from 2.35 to 7.35, and from 7.45 to 10.65 in panel (b).

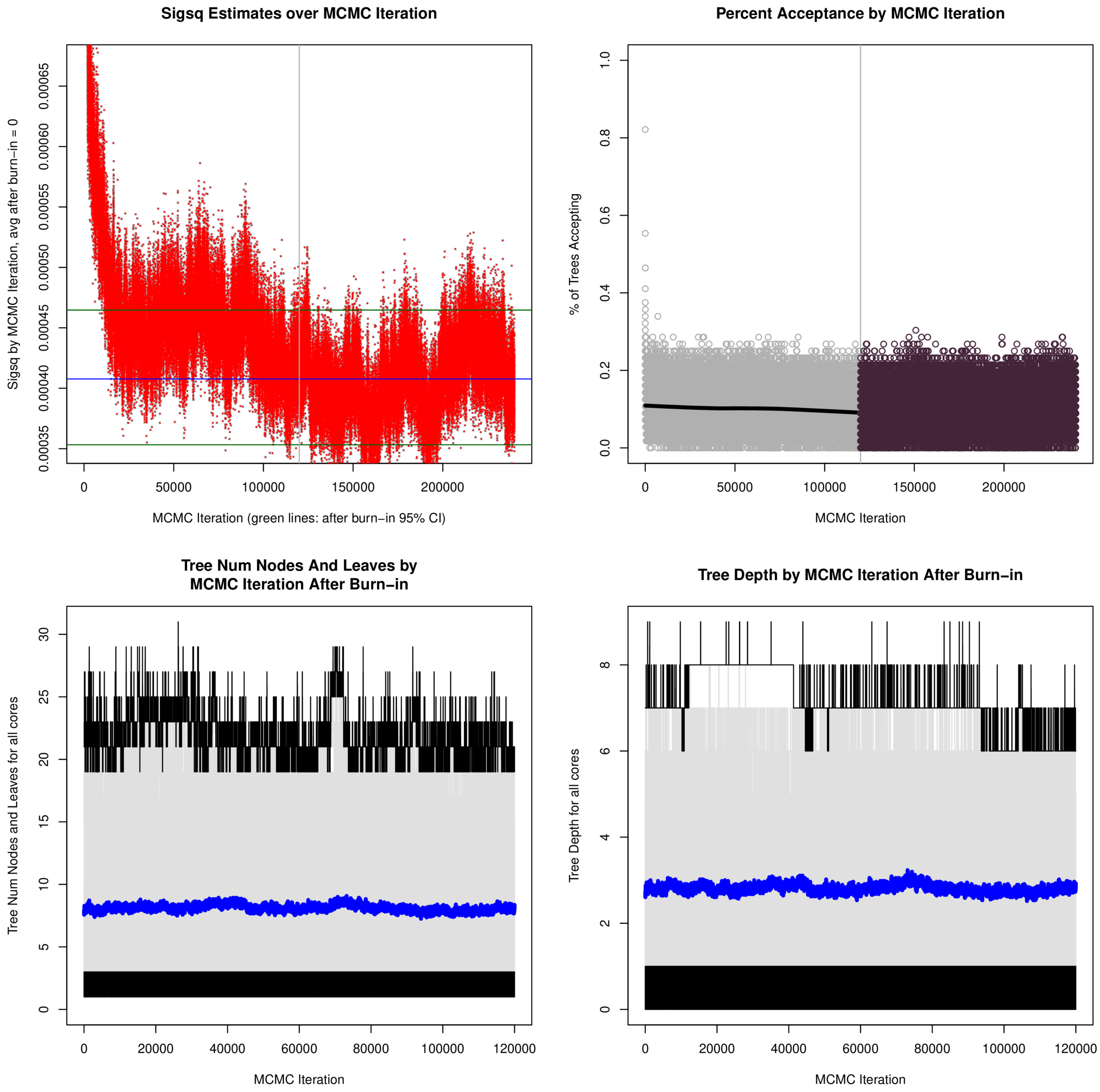

**Figure S13:** Convergence diagnostics for the BART analysis of the 200-tip simulation results with the RF distance between the true (simulated) and estimated (BLeSS MCC) supertree as the response variable. Top left: posterior estimate of $\sigma^2$ (error variance; "sigsq") by MCMC iteration. Note that due to the Metropolis-within-Gibbs sampling employed by BART, each MCMC iteration corresponds to one Gibbs sample but $2m+1$ Metropolis-Hastings (MH) steps, where $m$ is the number of trees (56 for the analysis of RF distances). The blue and green lines indicate the post-burnin mean and 95% credible interval (CI), respectively. Top right: proportion of MH steps accepted for each Gibbs sample, calculated as the number of accepted steps divided by $m$. The gray vertical line separates the pre-burnin and post-burnin iterations; the thick black line shows the LOESS smoothing results. Bottom left: number of nodes (including terminal nodes, i.e., leaves) per tree in each post-burnin Gibbs sample. Gray lines correspond to individual trees, black lines denote the trees with the minimum and maximum number of nodes, and the thick blue line shows the mean number of nodes over all $m = 56$ trees. Bottom right: tree depth (i.e., number of edges between the root and the most distant leaf) per tree in each post-burnin Gibbs sample. Gray lines correspond to individual trees, black lines denote the trees with the minimum and maximum depth, and the thick blue line shows the mean depth over all $m = 56$ trees. Note that the analysis was conducted on a single core to ensure deterministic output.

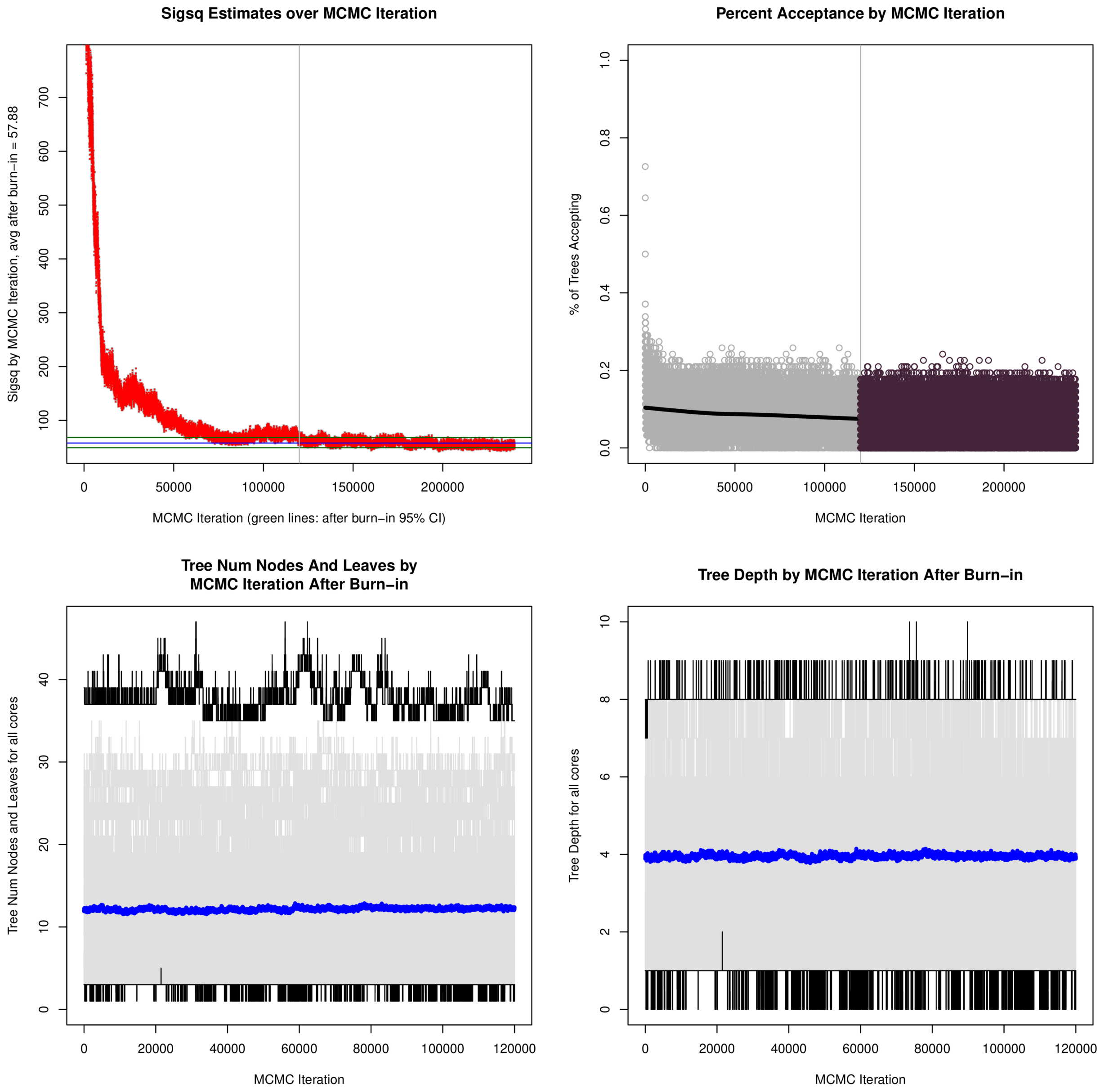

**Figure S14:** Convergence diagnostics for the BART analysis of the 200-tip simulation results with the KF distance between the true (simulated) and estimated (BLeSS MCC) supertree as the response variable. Panel descriptions as in Figure S13. The analysis employed $m = 62$ trees and was conducted on a single core to ensure deterministic output.

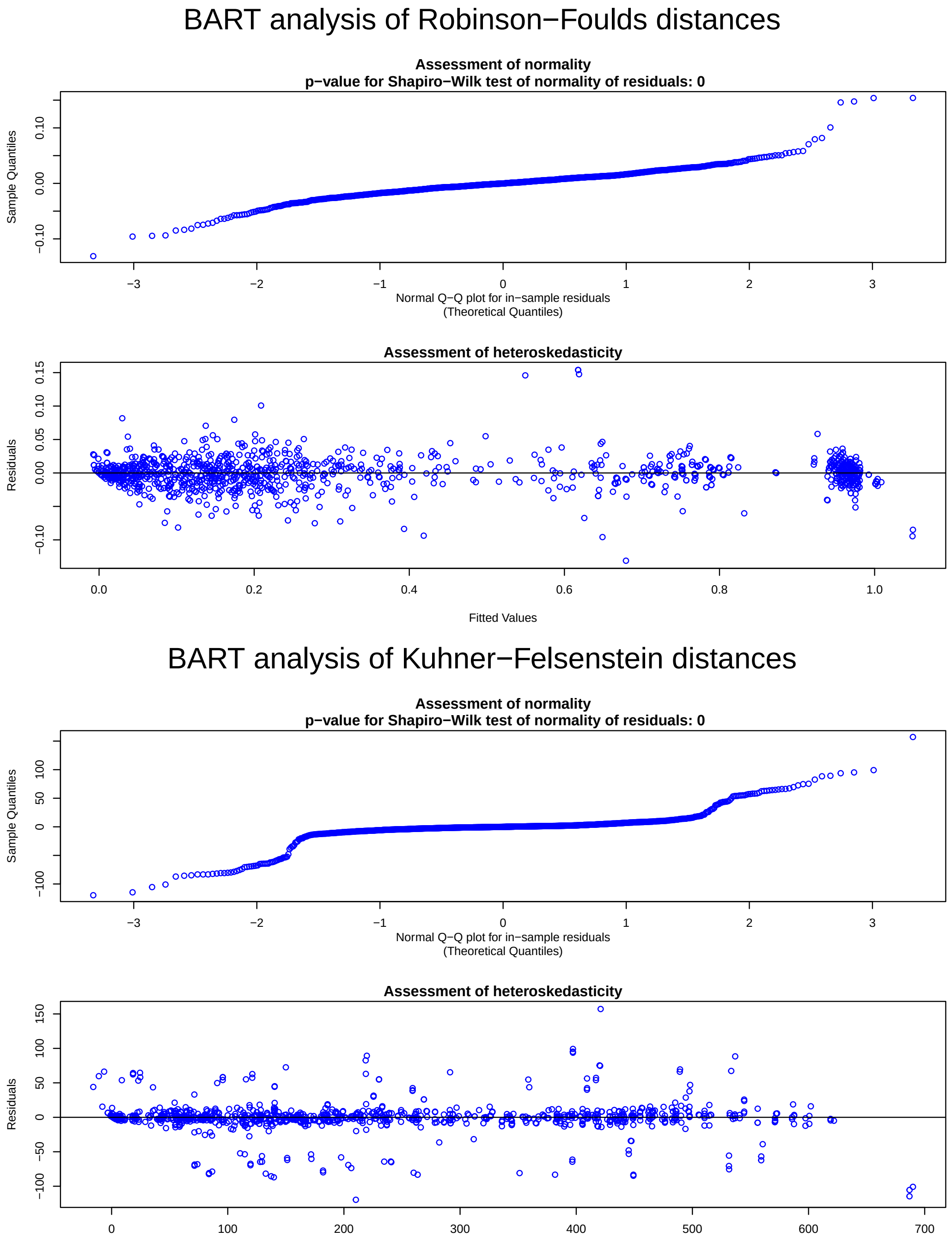

**Figure S15:** Error assumption checks for the BART analyses of the RF (top) and KF (bottom) distances between the true (simulated) and estimated (BLeSS MCC) supertrees. The model assumes that the errors are normally distributed and homoskedastic; the former is assessed using the Shapiro-Wilk test and a quantile-quantile plot of the actual residual distribution against the normal expectation, while the latter is tested by plotting the residuals against the fitted values of the response. In both analyses, the normality assumption is violated.

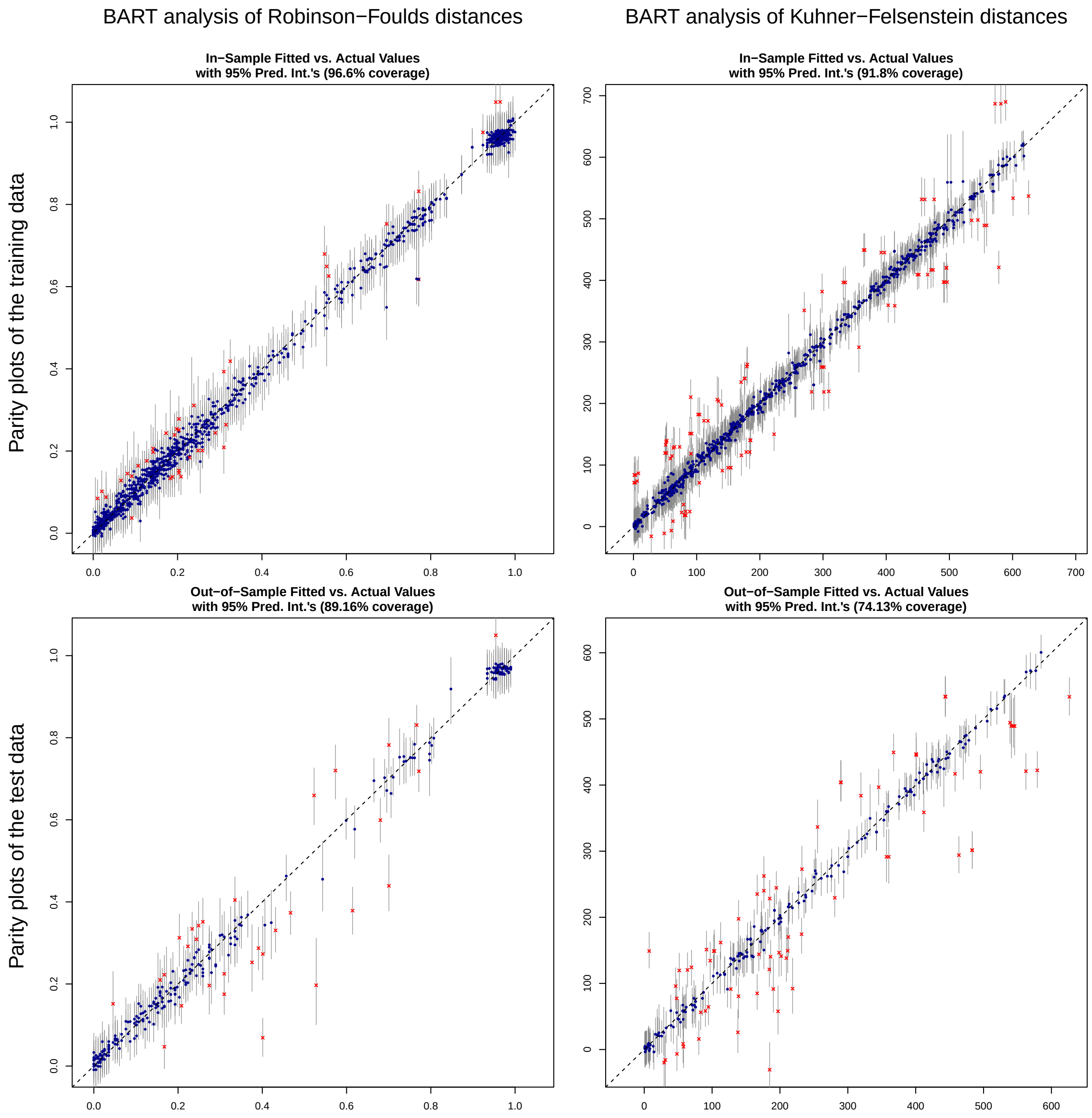

**Figure S16:** Parity plots of the fitted vs. actual response values for the training (top) and testing (bottom) sets from the BART analyses of the RF (left) and KF (right) distances between the true (simulated) and estimated (BLeSS MCC) supertrees. The gray vertical segments represent prediction intervals; blue dots and red crosses indicate actual response values that do and do not fall within the corresponding prediction interval, respectively.

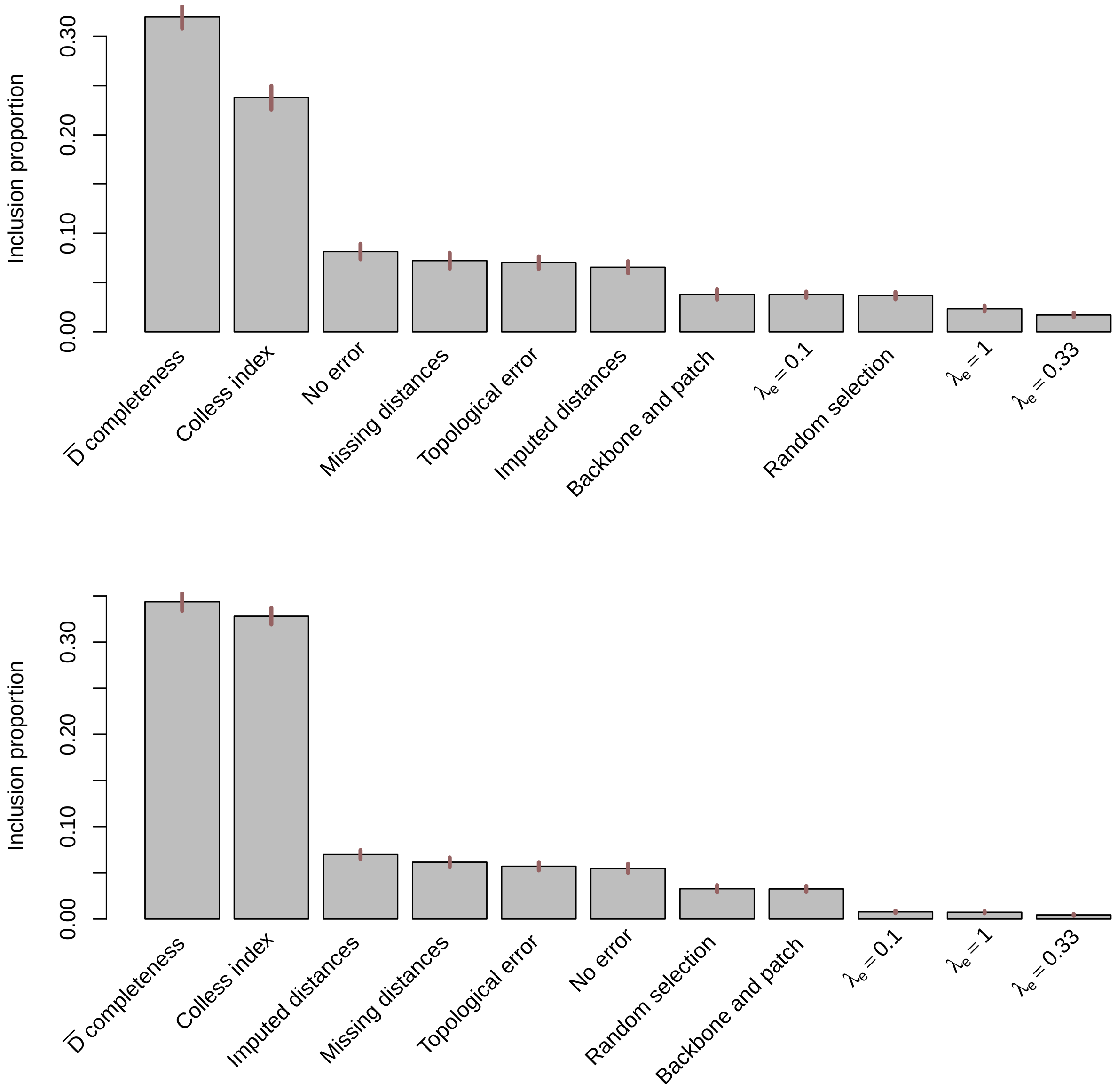

**Figure S17:** Predictor importance in the BART analyses of the RF (top) and KF (bottom) distances between the true (simulated) and estimated (BLeSS MCC) supertrees, as quantified by the number of splits that use a given predictor as a splitting variable divided by the total number of splits in the ensemble ("inclusion proportion"). This proportion is averaged across all $m$ trees (here, $m = 20$ to force the predictors to compete for inclusion), all post-burnin MCMC iterations (Gibbs samples), and 20 runs to ensure the sampler does not get stuck in a local maximum. The red segments represent 95% confidence intervals.

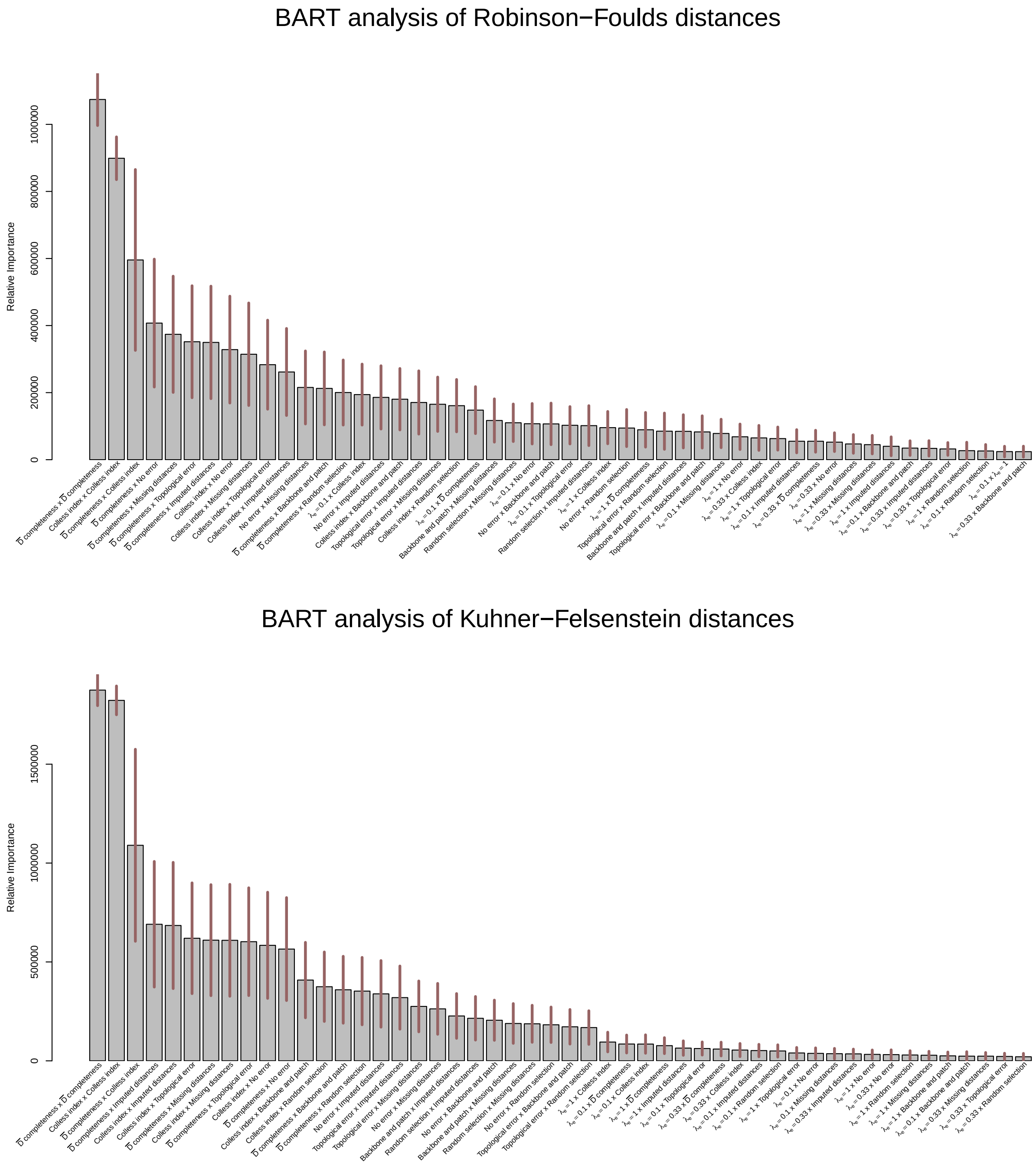

**Figure S18:** The top 50 predictor interactions in the BART analyses of the RF (top) and KF (bottom) distances between the true (simulated) and estimated (BLeSS MCC) supertrees. Two predictors are considered to interact if splits that use them as splitting variables appear in the same root-to-leaf path. The strength of interaction ("relative importance") is quantified by the number of times this occurs, summed over all $m$ trees (here, $m = 20$ to force the predictors to compete for inclusion), all post-burnin MCMC iterations (Gibbs samples), and 20 runs to ensure the sampler does not get stuck in a local maximum. The red segments represent 95% confidence intervals. Note that a predictor can interact with itself, such as when adjacent splits involve different thresholds for the same continuous variable.

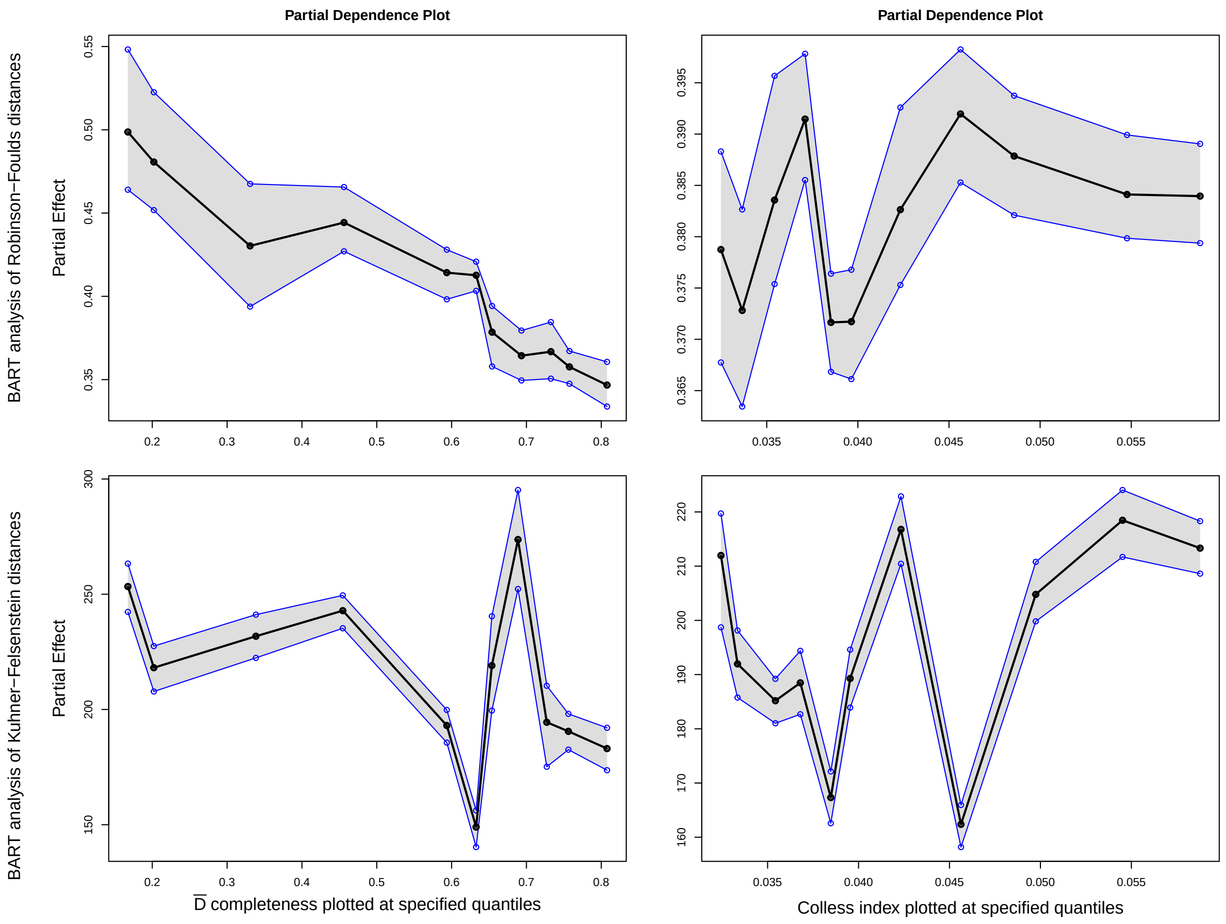

**Figure S19:** Partial dependence plots for the two continuous predictors in the BART analyses of the RF (top) and KF (bottom) distances between the true (simulated) and estimated (BLeSS MCC) supertrees. The partial effect of a given predictor represents the average contribution to the predicted response after fixing the predictor to a given value and letting the remaining predictors vary over their marginal distribution. The mean estimate of the partial dependence function is plotted in black at the 5th, 10th, . . ., 90th, and 95th percentiles, with 95% credible bands in gray and blue.

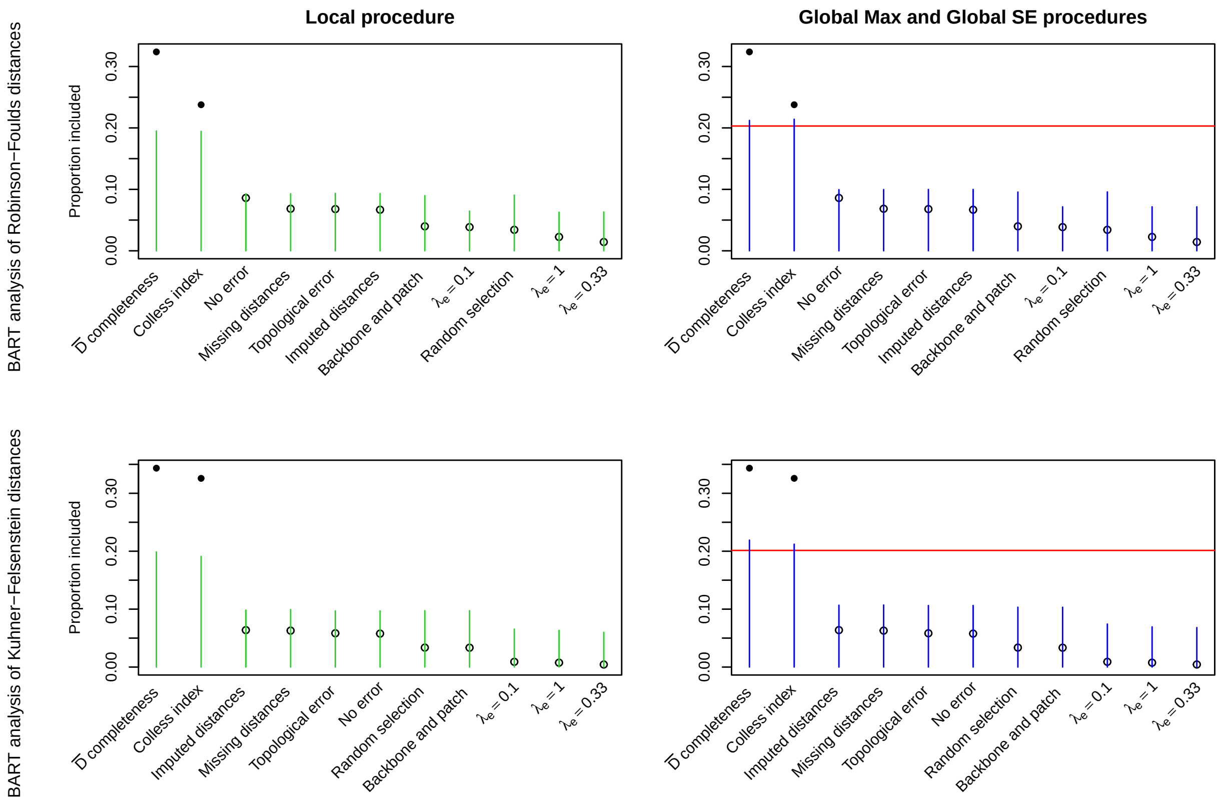

**Figure S20:** Results of three variable selection procedures performed on the BART analyses of the RF (top) and KF (bottom) distances between the true (simulated) and estimated (BLeSS MCC) supertrees. Left: results of the Local procedure, with green segments representing the permutation-derived inclusion proportion (see Figure S17) thresholds that a predictor has to exceed to be selected. Solid and open dots denote predictors that were and were not selected for inclusion in the model, respectively. Right: results of both the Global Max and Global SE procedures. The red line represents the Global Max cutoff, while the blue segments indicate the Global SE inclusion proportion thresholds. Solid dots denote predictors that exceeded both thresholds; open dots denote predictors that were not selected by either procedure. All thresholds were computed at $\alpha = 0.05$ using 100 permutations of the response and 25 BART runs with $m = 20$ trees each (to force the predictors to compete for inclusion). Note that all three procedures agree on the variables to be selected.

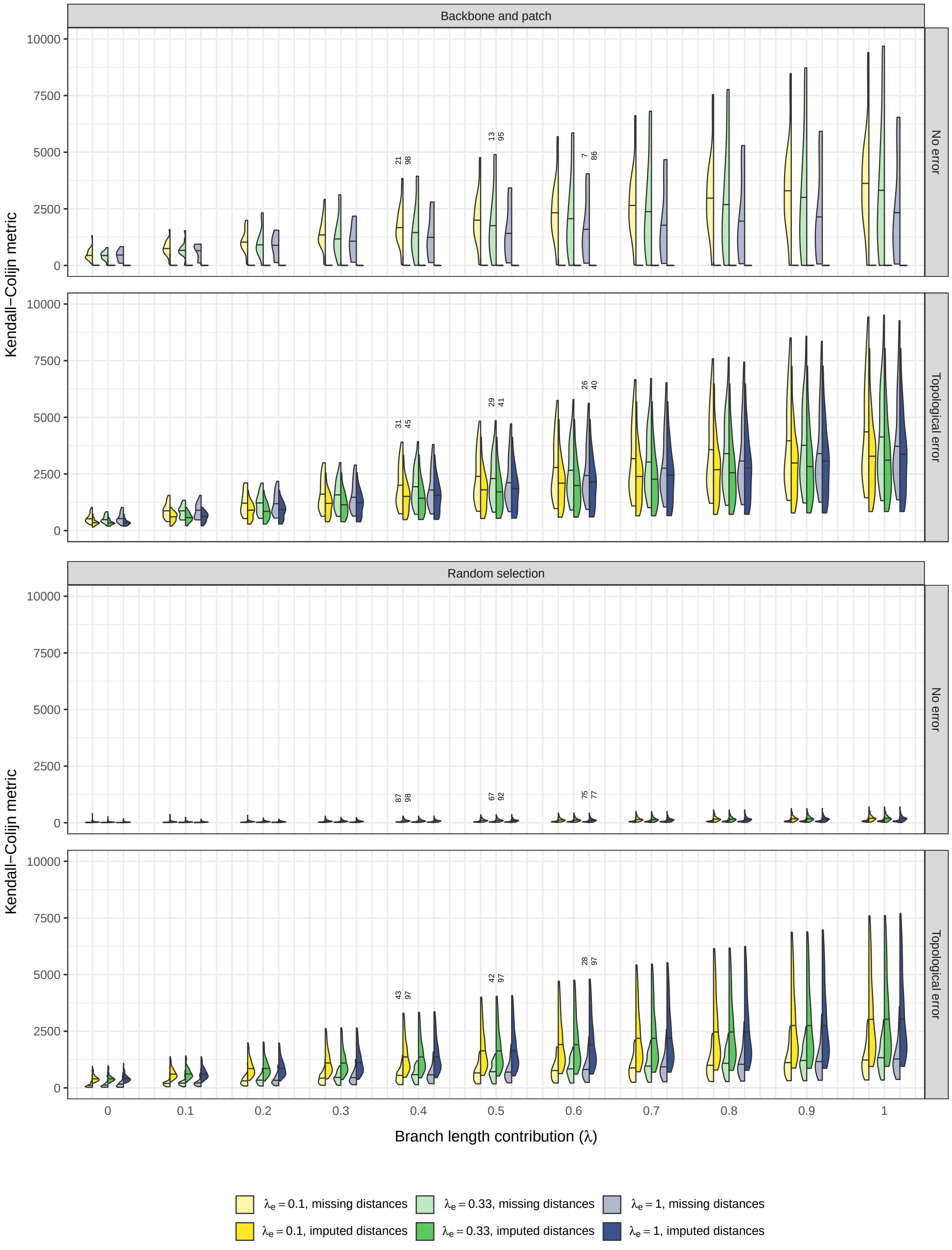

**Figure S21:** Distributions of Kendall-Colijn (KC) distances between the true (simulated) and estimated (BLeSS MCC) supertrees, plotted for different simulation scenarios and different values of the parameter $\lambda$, which determines the relative contribution of branch lengths to the KC metric ($\lambda = 0$: topology only, $\lambda = 1$: branch lengths only). The values above the violin plots denote the number of replicates that reached convergence for a given simulation scenario.

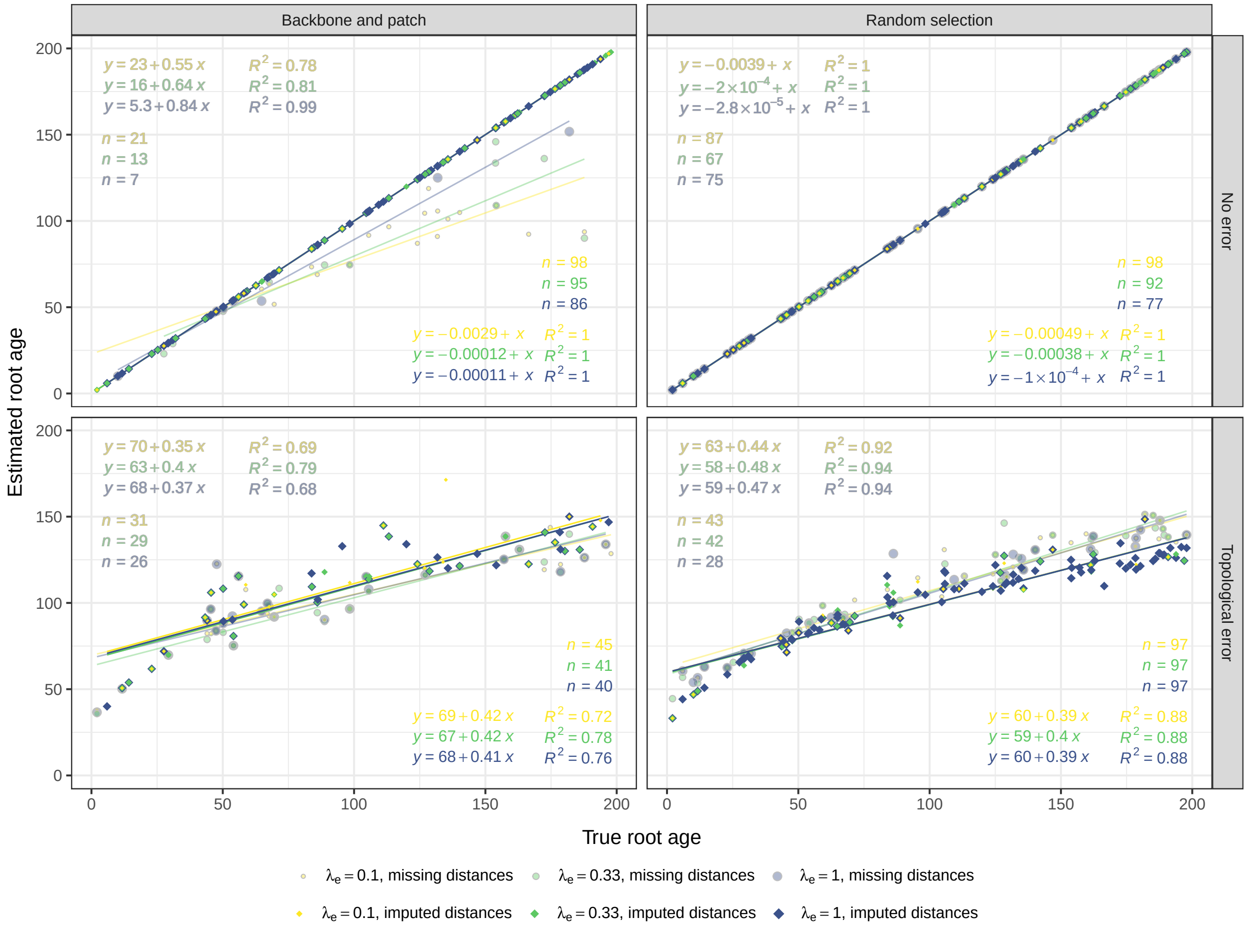

**Figure S22:** Scenario-specific relationships of the true (simulated; $x$) and estimated (BLeSS posterior mean; $y$) root ages, computed from all $n$ replicates that reached convergence. As in individual simulation scenarios, the two sets of ages also exhibit a strong and positive correlation globally (i.e., across scenarios: $y = 28 + 0.73x$; $R^2 = 0.83$).

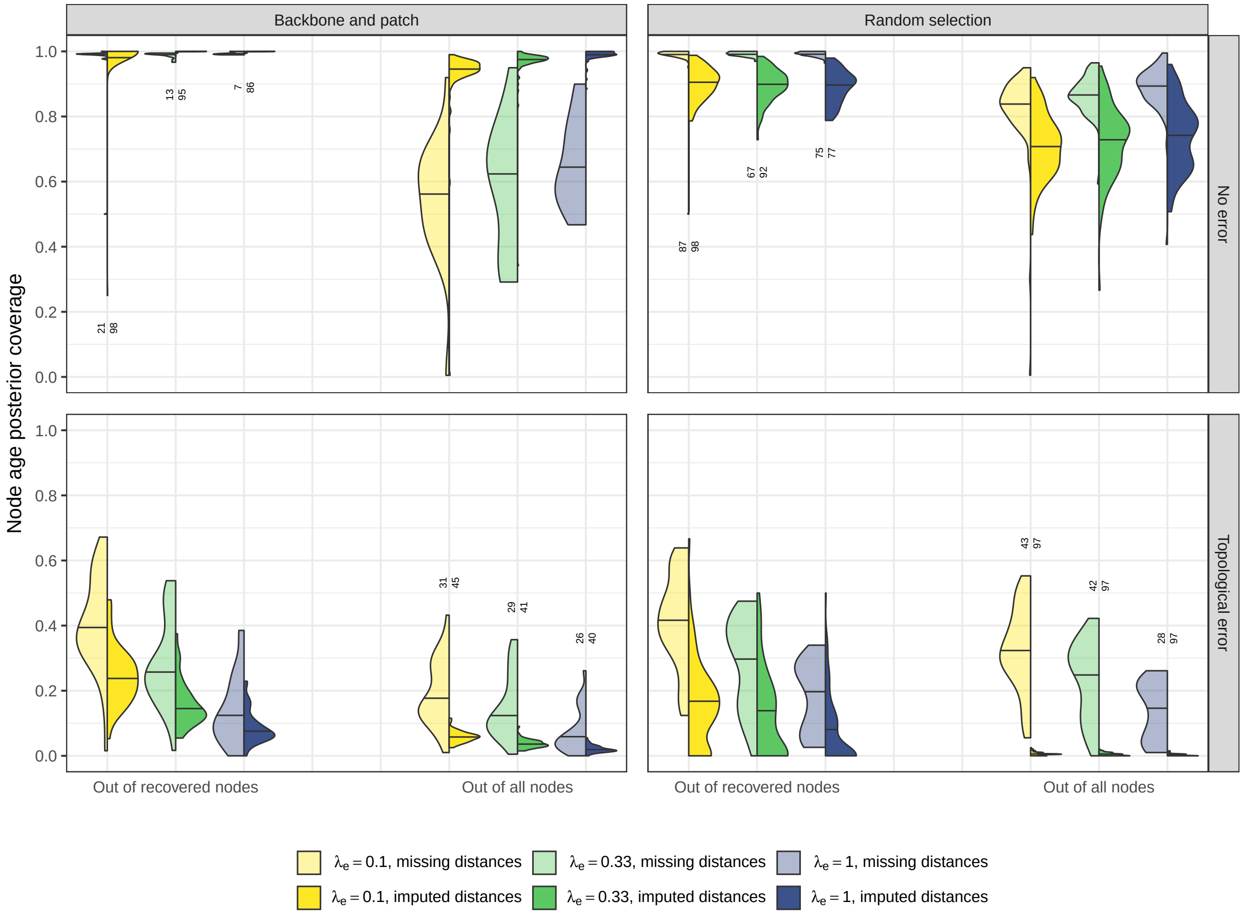

**Figure S23:** The proportion of nodes in a given tree whose true (simulated) age falls within the corresponding 95% highest posterior density interval yielded by BLeSS, computed for different simulation scenarios and either out of only those nodes that were present both in the true tree and the BLeSS supertree (left group of violin plots), or all 199 nodes (right group of violin plots). The values above the violin plots denote the number of replicates that reached convergence for a given simulation scenario. Note that the resulting proportions should not be strictly interpreted as posterior coverages, because the ages of individual nodes are not mutually independent.

#### 1.3 Scalability tests

**Table S8:** Properties of the large-scale simulated trees and their corresponding average distance matrices ($\bar{\mathbf{D}}$). Root height is given in arbitrary units.

| Tip number | Root height | Colless index | Subtree number | $\bar{\mathbf{D}}$ completeness |
|---|---|---|---|---|
| 500 | 552.15 | 0.0208 | 11 | 0.8893 |
| 1000 | 621.46 | 0.012 | 17 | 0.8578 |
| 5000 | 782.40 | 0.0031 | 18 | 0.9125 |
| 10,000 | 851.72 | 0.0016 | 22 | 0.9081 |

**Table S9:** Model and MCMC settings used for the large-scale tree analyses. Sampling freq. = frequency at which samples of scalar parameters and trees are extracted from the chain and logged; Printing freq. = frequency at which elapsed time and select parameter values are written to the standard output. Chain lengths and frequencies are given in MCMC moves.

| Tip number | $\lambda_e$ | Target chain length | Sampling freq. | Printing freq. | Checkpointing freq. |
|---|---|---|---|---|---|
| 500 | 1 | $5 \times 10^6$ | $2.5 \times 10^3$ | 200 | $5 \times 10^4$ |
| 1000 | 1 | $5 \times 10^6$ | 500 | 50 | $2.5 \times 10^4$ |
| 5000 | 1 | $5 \times 10^6$ | 100 | 4 | $1.6 \times 10^4$ |
| 10,000 | 1 | $5 \times 10^6$ | 50 | 2 | $8 \times 10^3$ |

**Table S10:** Wall-clock time spent on individual operations underlying BLeSS as a function of tree size and missing data treatment. All values are averaged over 10 replicates and measured with millisecond precision for the MPI version of `RevBayes` v1.2.2 (commit b083532) run with 32 threads per replicate on a compute cluster consisting of Intel Xeon Gold 6248R processors.

| | Time (s); no imputation | | | |
|---|---|---|---|---|
| Operation | 500 tips | 1000 tips | 5000 tips | 10,000 tips |
| Convert source trees to distance matrices | 0.118 | 0.225 | 0.868 | 2.341 |
| Compute the average distance matrix $\bar{\mathbf{D}}$ | 0.021 | 0.087 | 1.919 | 7.765 |
| Draw a tree $\hat{\mathcal{T}}$ from a birth-death process | 0.032 | 0.125 | 3.100 | 12.566 |
| Convert the proposed tree $\hat{\mathcal{T}}$ to a distance matrix $\hat{\mathbf{D}}$ | 0.033 | 0.139 | 4.156 | 18.872 |
| Draw from the exp. error distribution conditional on $\hat{\mathbf{D}}$ | 0.071 | 0.293 | 8.022 | 33.942 |
| Clamp the distribution to $\bar{\mathbf{D}}$ | 0.016 | 0.072 | 1.630 | 6.867 |
| Compute the log probability density | 0.003 | 0.006 | 0.143 | 0.612 |
| | Time (s); imputation | | | |
| Read in an imputed distance matrix | 0.108 | 0.486 | 38.935 | 291.262 |
| Convert the imputed matrix to type `AverageDistanceMatrix` | 0.032 | 0.141 | 4.486 | 19.944 |
| Draw a tree $\hat{\mathcal{T}}$ from a birth-death process | 0.032 | 0.124 | 3.131 | 12.630 |
| Convert the proposed tree $\hat{\mathcal{T}}$ to a distance matrix $\hat{\mathbf{D}}$ | 0.033 | 0.140 | 4.156 | 18.874 |
| Draw from the exp. error distribution conditional on $\hat{\mathbf{D}}$ | 0.071 | 0.291 | 7.972 | 34.345 |
| Clamp the distribution to $\bar{\mathbf{D}}$ | 0.028 | 0.126 | 4.220 | 20.428 |
| Compute the log probability density | 0.003 | 0.011 | 0.249 | 1.452 |

**Table S11:** Wall-clock runtime per 1 million MCMC iterations (under `moveschedule="single"`, i.e., with a single Metropolis-Hastings move proposed per iteration) as a function of tree size and missing data treatment. All values are averaged over 5 replicates and were calculated for the single-core version of RevBayes v1.2.2 (commit ed422ba) on a compute cluster consisting of Intel Xeon Gold 6248R processors. See also Figure S24.

| | Time (s) | |
|---|---|---|
| Tree size | No imputation | Imputation |
| 500 tips | 26,482 s ≈ 7.4 hours | 26,097 s ≈ 7.2 hours |
| 1,000 tips | 110,954 s ≈ 30.8 hours | 108,426 s ≈ 30.1 hours |
| 5,000 tips | $3.251 \times 10^6$ s ≈ 5.4 weeks (projected) | $3.231 \times 10^6$ s ≈ 5.3 weeks (projected) |
| 10,000 tips | $1.384 \times 10^7$ s ≈ 5.3 months (projected) | $1.464 \times 10^7$ s ≈ 5.6 months (projected) |

**Table S12:** Acceptance rates of different MCMC moves applied during BLeSS inference either directly to the supertree (`tree`) or to the hyperparameters of the birth-death prior from which the supertrees are drawn (`speciation`, `extinction`, `tree_height`), tabulated as a function of tree size and missing data treatment. The rates were calculated from the following total numbers of proposed moves: $5 \times 10^6$ (500 tips), $1.15 \times 10^6$ (1000 tips), 40,000 (5000 tips), and 8000 (10,000 tips). The weight of a particular move represents the relative frequency at which it is proposed (with the actual frequency obtained by dividing its weight by the sum of all move weights).

| | | | Acceptance rate (no imputation) | | | |
|---|---|---|---|---|---|---|
| Move | Parameter | Weight | 500 tips | 1000 tips | 5000 tips | 10,000 tips |
| `Scaling` | `speciation` | 3 | 0.2131 | 0.1646 | 0.0955 | 0.0533 |
| `Scaling` | `extinction` | 3 | 0.2185 | 0.2241 | 0.9428 | 0.9203 |
| `Scaling` | `tree_height` | 2 | 0.0007 | 0.0025 | 0.0113 | 0.0392 |
| `NarrowExchange` | `tree` | 10 | 0.1509 | 0.2285 | 0.4870 | 0.4949 |
| `NNI` | `tree` | 10 | 0.1178 | 0.1836 | 0.4522 | 0.4581 |
| `FNPR` | `tree` | 10 | 0.0034 | 0.0235 | 0.4238 | 0.4974 |
| `SubtreeScale` | `tree` | 10 | 0.2404 | 0.3157 | 0.5624 | 0.5458 |
| `NodeTimeSlideUniform` | `tree` | 15 | 0.3145 | 0.4152 | 0.7511 | 0.7533 |
| | | | Acceptance rate (imputation) | | | |
| `Scaling` | `speciation` | 3 | 0.4271 | 0.4003 | 0.1039 | 0.0545 |
| `Scaling` | `extinction` | 3 | 0.4279 | 0.4115 | 0.9580 | 0.9212 |
| `Scaling` | `tree_height` | 2 | 0.0001 | 0.0004 | 0.0054 | 0.0183 |
| `NarrowExchange` | `tree` | 10 | 0.0331 | 0.0866 | 0.2904 | 0.3604 |
| `NNI` | `tree` | 10 | 0.0185 | 0.0496 | 0.2591 | 0.3263 |
| `FNPR` | `tree` | 10 | 0.0046 | 0.0264 | 0.4715 | 0.4912 |
| `SubtreeScale` | `tree` | 10 | 0.0756 | 0.1016 | 0.3120 | 0.3579 |
| `NodeTimeSlideUniform` | `tree` | 15 | 0.0938 | 0.1714 | 0.4999 | 0.5598 |

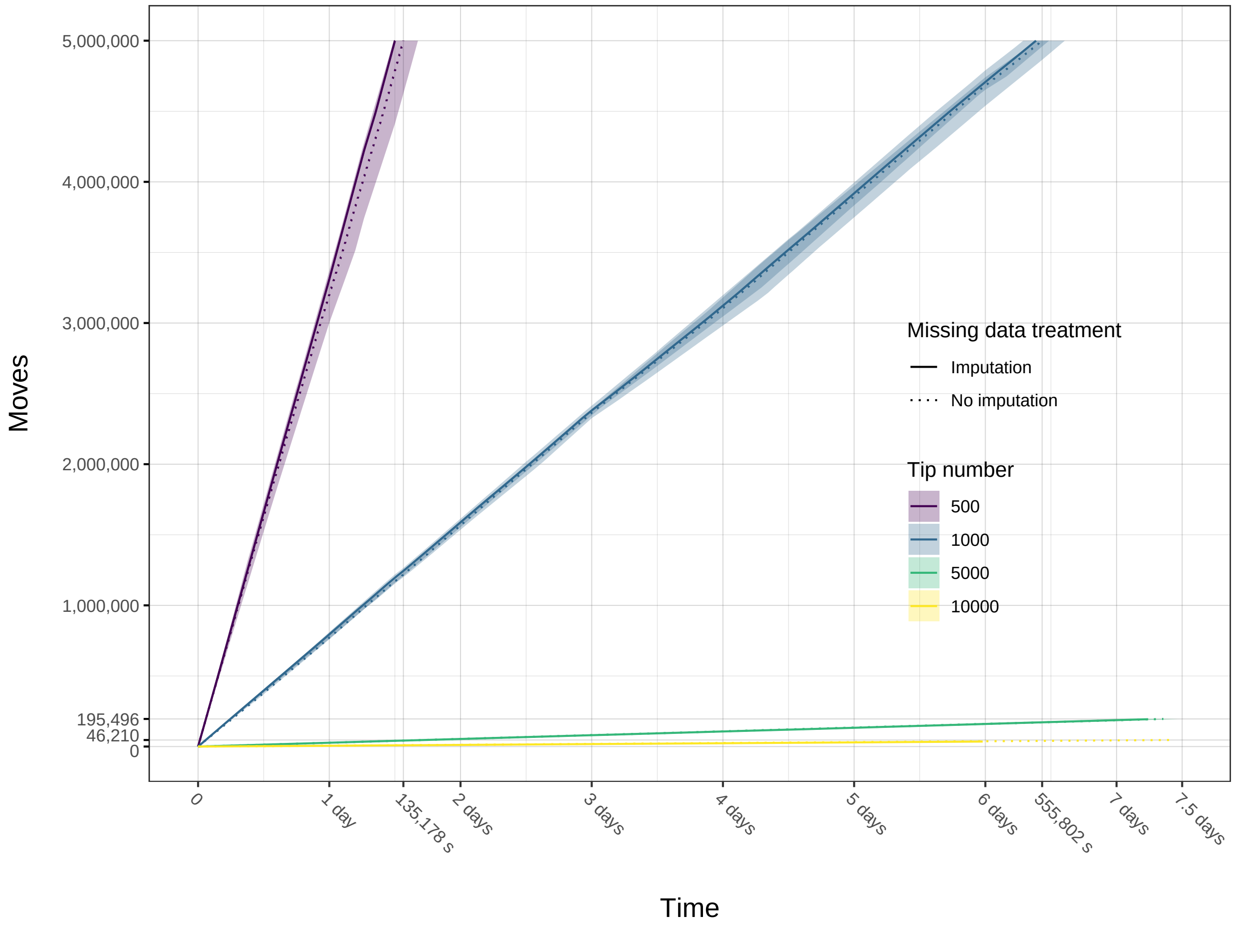

**Figure S24:** Number of MCMC iterations (under `moveschedule="single"`, i.e., with a single Metropolis-Hastings move proposed per iteration) as a function of wall-clock time, plotted for different tree sizes and missing data treatments. Lines denote mean values across 5 replicates (performed on the same simulated dataset but under different random seeds); shaded bands indicate the corresponding range. The maximum runtime of the analyses that reached the target chain length of 5 million iterations within the time limit of 7.5 days (500 tips, 1000 tips) is recorded along the $x$-axis. For the analyses that failed to do so (5000 tips, 10,000 tips), the maximum number of iterations completed before the time limit was reached is recorded along the $y$-axis. When restarting analyses from a checkpoint file, the iterations performed after the last saved state but before a timeout were counted toward the total.

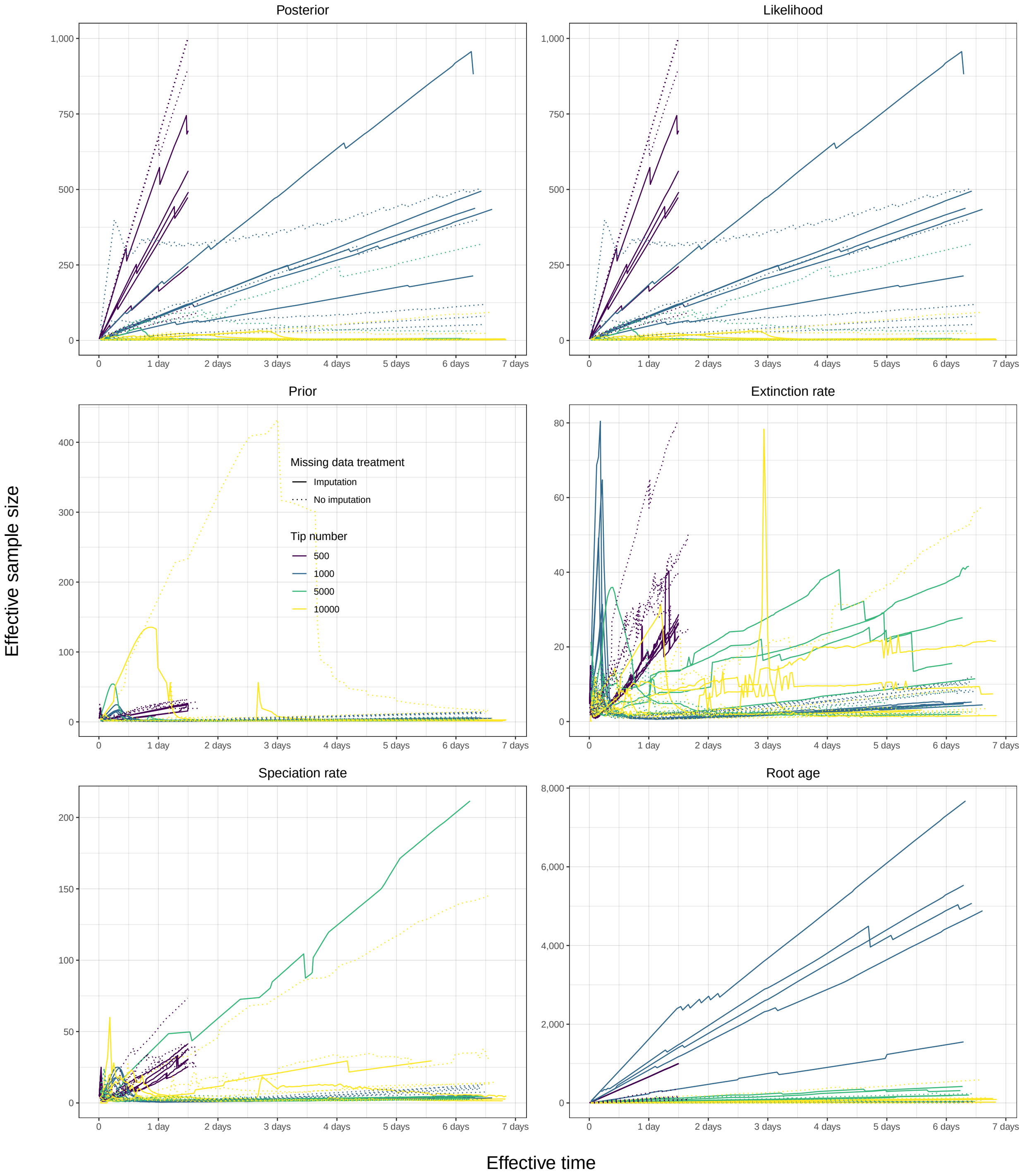

**Figure S25:** Effective sample sizes of scalar parameters as a function of wall-clock time, plotted for different tree sizes and missing data treatments. Each of the 5 replicates (performed on the same simulated dataset but under different random seeds) is plotted as a separate line. When restarting analyses from a checkpoint file, the iterations performed after the last saved state but before a timeout were not counted toward the total; as a result, the effective time was less than the total runtime.

### 2 Empirical Application to the Order Carnivora

Our literature search yielded 41 candidate studies that carried out divergence time estimation for the order as a whole or various subclades thereof. We evaluated these studies for their suitability as BLeSS input, which was largely determined either by the availability of the resulting time tree(s) in a machine-parsable format (Newick or Nexus) or by the possibility of assembling such a time tree from other information provided (e.g., a figure illustrating the topology of the tree plus a table of divergence times). Below, we provide a brief description of each study, including its focal clade and (if applicable) the number of source trees extracted from it. Included studies are indicated by an asterisk (*).

**Bagatharia et al. (2013)*** *Felidae.* Estimated divergence times for a small sample of felids using whole mitochondrial genomes and a relaxed uncorrelated molecular clock in BEAST v1.7.1. They used a single calibration point at 10.78 Ma for Felidae. Although no tree file is available, we could reconstruct the topology and divergence times from the paper.

**Barrett et al. (2021)** *Carnivora.* Used BEAST v2.6.3 to infer a total-evidence tip-dated tree of carnivorans using 223 morphological characters, three nuclear, and 13 mitochondrial protein-coding loci (14,624 bp) for 13 extant and 32 extinct ingroup species as well as 3 outgroups. They did not provide a summary tree but did provide a posterior sample of trees in the supplement that were used for macroevolutionary analyses. Within this sample, extant tips are found to be non-contemporaneous because of the use of stratigraphic ranges for taxa with fossil records. Rather than extending tip branches or re-inferring the tree with these tips treated as extant, we opted not to use this dataset.

**Bon et al. (2008)*** *Ursidae.* Used BEAST to infer phylogeny and divergence times for extant ursids and the extinct cave bear from mitochondrial genomes and two fossil calibrations under an uncorrelated lognormal relaxed clock. No tree file is available and the tree is reconstructed from Figure 5, with divergence times taken from the associated figure legend.

**Davis et al. (2010)*** *Felidae.* Generated novel sequence data and employed Bayesian species tree analysis, followed by time scaling in PAML/multidivtime, for the big cats *Panthera* plus one outgroup. They used three minimum age calibrations and investigated the effects of moving each. As results are similar, we use the divergence times from analysis with the complete set of priors here. A tree file is not provided, so we used their Figure 8 to reconstruct the chronogram.

**dos Reis et al. (2012)** *Carnivora.* Used mitochondrial and nuclear loci to infer a dated phylogeny of 274 placental mammal species, including a number of carnivorans. However, no tree file or table of divergence times is available, with a figure showing topology being all that is available in the supplement.

**Eizirik et al. (2010)*** *Carnivora.* Sampled 14 genes across Carnivora and inferred the time scale of evolution using the Thorne-Kishino autocorrelated-rates clock model in divtime5b and a relaxed uncorrelated clock in BEAST v1.5.2, with 25 fossil calibrations.

No tree files are available but we were able to manually digitize the tree and annotated it with mean ages across the divtime and BEAST analyses, where both were provided (their Table 6). For nodes not represented in Table 6, we used the divtime dates reported in Table S1 that were obtained from analyses with a 55 Ma root prior.

**Fulton and Strobeck (2010)*** *Phocidae.* Used fifteen nuclear genes and mitochondrial genomes and 8 fossil calibrations to infer topology and branch lengths in BEAST v1.4.8 for all extant phocid species as well as the walrus, three otariids and seven carnivore outgroups (two felids, two canids, three mustelids). We transcribed their time tree from Figure 2 and Table 4.

**Gaubert and Begg (2007)*** *Viverridae.* Employed two mitochondrial loci (cytochrome *b* and 5' domain of the control region) and one Y-linked (terminal exon of zinc-finger) gene to infer the phylogeny of the genus *Genetta* using likelihood and Bayesian methods. They then time-scaled the CytB tree under a relaxed clock using PAML/multidivtime with a single calibration point assigned to the root. We transcribed their time tree from Figure 2, omitting duplicated individuals within a single species and treating the divergence of *G. maculata* and *G. tigrina* as the older of the two ages.

**Gaubert and Cordeiro-Estrela (2006)*** *Viverridae.* Used penalized likelihood, as implemented in r8s v1.7, to time-scale a tree of feliform carnivorans, with sampling focused on Viverrinae, inferred from 2 nuclear and one mitochondrial gene. No tree file is available, so we digitized the tree plotted in Figure 4, which is described as a linearized tree showing minimum divergence time estimates. Several taxa (*Cryptoprocta ferox*, some *Genetta* species) were pruned from the tree as divergence times were not provided.

**Hassanin et al. (2021)** *Carnivora.* Added 51 new mitogenomes from 13 families to previously sequenced mitogenomes for an alignment of 220 taxa. They time-calibrated their tree using BEAST v2.4.7 with 21 fossil calibration points. Due to the completeness of this data set it is not used here for assessing the performance of our supertree method.

**Helgen et al. (2013)** *Procyonidae.* Used sequence data analyzed in BEAST to infer a time-calibrated phylogeny of Procyonidae while also describing a new species of olingo *Bassaricyon neblina*. However, no tree file is available and the descriptions of divergence times in the text are insufficient to reconstruct the chronogram manually. This dataset is therefore excluded.

**Higdon et al. (2007)*** *Pinnipedia.* Assembled a supertree of pinnipeds, plus a canid and ursid outgroup, from a weighted matrix representation with parsimony (MRP) analysis of 50 gene trees inferred under maximum likelihood, and time-scaled the resulting topology using a perl script to calibrate relative branch lengths from the gene trees to a set of fossils. We took the topology from Figure 1 and the divergence time estimates from Table 2.

**Johnson et al. (2006)*** *Felidae.* Used 22,789 base pairs of nuclear and mitochondrial data from 37 extant felid species along with several outgroups to infer a maximum likelihood phylogeny using PAUP*. They then employed 16 fossil calibrations to timescale their tree using the nuclear loci alone in the program estbranches. A tree file is not available but we were able to reconstruct the tree topology and branch lengths from their Figure 1 and Table 1.

**Koepfli et al. (2006)*** *Hyaenidae.* Used 7 nuclear loci and 1 mitochondrial gene to infer a maximum likelihood phylogeny for the four extant species of Hyaenidae and a set of feliform outgroups, and then time-scaled this topology using a relaxed molecular clock as implemented in divtime5b based on 9 fossil constraints. We manually digitized the tree topology from Figure 2 and added mean node ages from Table 6.

**Koepfli et al. (2007)*** *Procyonidae.* Used data from 9 nuclear and 2 mitochondrial genes to infer the phylogeny of Procyonidae and estimated divergence times under a relaxed molecular clock using divtime5b with a variable set of fossil calibrations. We manually reconstructed the maximum likelihood topology from Figure 3 and sampled all three sets of divergence times from Table 4.

**Koepfli et al. (2008)*** *Mustelidae.* Used 22 gene segments for 44 mustelids plus 2 procyonid outgroups to infer a time-scaled phylogeny using BEAST v1.4.2. They employed several calibration strategies, reporting the mean and 95% highest posterior density (HPD) intervals for each. We used the mean of the mean ages for each node. We note that divergence times were not provided for the splits between *Bassariscus astutus* and *Procyon lotor*, *Enhydra lutris* and *Hydrictis maculicollis*, and *Martes americana* and *Martes melampus*. In each case, we deleted the first named taxon from the tree.

**Krause et al. (2008)*** *Ursidae.* Also used mitochondrial sequences for extant bears, plus two extinct taxa (short faced and cave bears) and inferred phylogeny and branch lengths using BEAST v1.4.4 under a strict molecular clock. The tree is manually reconstructed from Figure 1 of their paper.

**Kutschera et al. (2014)*** *Ursidae.* Used *BEAST to jointly estimate gene trees for 14 autosomal introns and 9 Y-chromosome genes, and the species tree for all 8 extant bear species under a multilocus coalescence approach. The topology with highest posterior probability (their Figure 2A) is used here, with posterior mean divergence times taken from Table 1.

**Law et al. (2018)*** *Musteloidea.* Used a 46-gene dataset in combination with 74 fossil ages to infer a time-scaled phylogeny for 75 species of musteloid carnivores under a relaxed uncorrelated lognormal molecular clock and fossilized birth-death tree prior in BEAST v2.3.2. We use the maximum clade credibility tree from this study.

**Li et al. (2016)*** *Felidae.* Used phylogenomic data to investigate phylogeny, divergence times and introgression in felids. No tree files are provided, so we used the topology and mean branch lengths derived from multidivtime analyses performed on the biparental nuclear DNA genome partition, with different partitions and constraints, as reconstructed from Supplemental Figure S5A and mean ages from Supplemental Table S3.

**Lindblad-Toh et al. (2005)** *Canidae.* Presented a time-scaled phylogeny of canids based on ∼15 kbp of exon and intron sequences. However, the figure does not include all node ages and a tree file is not available. This dataset is not included here.

**Lopes et al. (2021)*** *Otariidae.* Used whole-genome sequencing for 14 of 15 extant species of Otariidae and inferred time-calibrated species trees using ASTRAL and StarBeast 2.

**Lynch (2020)*** *Mustelidae.* Used 12S, 16S, cytochrome b, and D-loop sequences for *Martes americana*, *M. caurina*, and *M. foina*, combined with three fossil calibrations, to infer the timescale of divergence using BEAST v1.10.4. The tree was manually transcribed from Figure 3 with mean divergence times recorded from the text.

**McDonough et al. (2022)*** *Mephitidae.* Used a multilocus nuclear and mitochondrial dataset to recognize seven distinct species of *Spilogale*. They estimated divergence times in BEAST v2.6.1 based on 13 protein-coding genes from the mitochondrial genome and calibrated the molecular clock using one fossil calibration and one molecular date for the origin of American mephitids.

**Meredith et al. (2011)*** *Carnivora.* Used a molecular dataset comprising 26 genes with a variety of treatments and fossil calibration schemes to infer the timescale of mammalian diversification. We extracted the carnivoran portion of their trees and scaled branches to mean divergence times reported over calibration schemes in the supplementary materials. The topology of the tree differed in resolution at the base of Arctoidea between analysis of amino acids vs nucleotides (ursids as sister to pinnipeds in the former vs. ursids as sister to pinnipeds plus musteloids in the latter), so we included two versions of the tree, with divergence times taken from the appropriate dataset.

**Mitra et al. (2019)*** *Feliformia.* Used protein coding loci from complete mitochondrial genomes and BEAST v1.7 to infer the timescale of feliform evolution from 24 species, plus a canid outgroup (*Cuon*). We transcribed their maximum clade credibility tree from Figure 2 of their paper.

**Nascimento et al. (2021)*** *Felidae.* Compiled previously published mitochondrial sequences covering ATP8, control region, CytB, and ND5 for for the pampas cat complex and inferred topology and divergence times under a relaxed clock in BEAST v1.8.2 using 2 calibrations. Based on the resulting clades, combined with morphological features, they recognized five species of pampas cat. A tree file was not available for this study but the topology and branch lengths could be manually reconstructed based on the mean ages labeled in Figure 8a of their paper.

**Nigenda-Morales et al. (2019)** *Procyonidae.* Used three mitochondrial loci genotypes from 11 microsatellite loci and BEAST v2.3.1 to examine the phylogeographic history of the white-nosed coati *Nasua narica* but also included samples from other procyonid species. However, no supplementary tree files are available and divergence times are not listed or described in the text. This dataset therefore cannot be used.

**Ogilvie et al. (2022)*** *Canidae*. Introduced a new model, the FBD-MSC, that combines the multispecies coalescent (MSC) and the fossilized birth–death (FBD) processes. They used the approach to infer the phylogeny and divergence times of all extant Canidae using a combined dataset of 58 nuclear genes and 230 morphological characters. Following their methods, we combined the sampled trees from four posterior chains (sb2-extant-only) provided in their online materials after removing the first 2% of each as burnin and produced a maximum clade credibility tree from the resulting output, annotated with mean divergence times.

**Paijmans et al. (2017)*** *Felidae*. Used mitogenomes to investigate the phylogeny and divergence times of sabertoothed cats (Machairodontinae) with respect to a sample of extant felids and feliform carnivores. They do not provide a tree file and, although the time-scaled topology is figured in their Figure 1, they do not provide a table of divergence times. We were able to extract the topology for Felidae that encompassed the two genera of sabertoothed cats and two felines spanning the root node.

**Patou et al. (2008)*** *Viverridae*. Used two mitochondrial and two nuclear loci, combined with a relaxed molecular clock to infer the phylogeny and divergence times of a set of hemigaline and paradoxurine viverrids. No tree file is available but we were able to manually reconstruct the tree topology and branch lengths using the divergence times estimated from the complete dataset. Note that the authors omitted the otter civet *Cynogale bennetti* from their combined analyses.

**Patou et al. (2009)** *Herpestidae*. Used 2 mitochondrial (CytB and ND2) and 2 nuclear ($\beta$-fibrinogen intron 7 and transthyretin intron 1) loci from almost all of the recognized mongoose species to produce a phylogeny of the Herpestidae. Divergence ages were estimated using a relaxed molecular clock approach using PAML/multidivtime. TTRi1 was excluded from divergence time estimation as it was missing for too many taxa. The tree is not available online and the figure is insufficient to reconstruct branch lengths. Therefore, this study is not included.

**Patou et al. (2010)** *Viverridae*. Used mitochondrial sequence data to estimate the phylogeny and divergence times of the genus *Paradoxurus*, including the identification of putative distinct, species-level OTUs within *P. hermaphroditus*. The did not provide a tree file, and while a time tree figure is shown in the paper, insufficient detail is provided to transcribe it. We therefore omitted this dataset.

**Perri et al. (2021)*** *Canidae*. Time-scaled a maximum likelihood topology inferred for 10 extant canids plus the dire wolf *Aenoycon dirus* using a nuclear supermatrix in PAML/MCMCTree. A time tree file is not available, so we manually transcribed the tree and branch lengths from Figure 2a. We retain the dire wolf here as its terminal age is within the rounding error of the branch durations.

**Püschel et al. (2020)** *Ursidae*. Used a combination of molecular and morphological data for extant and extinct ursids, as well as a canid outgroup to infer branch lengths under an independent gamma rates (IGR) relaxed clock in MrBayes. Although tree files are available, the resulting branching time estimates are unusually young, and are therefore excluded here.

**Rao et al. (2020)** *Hyaenidae.* Used proteomic data from 3 extant species of hyena as well as the extinct cave hyena (Eurasian *Crocuta crocuta*) to infer the timescale of hyaenid evolution using BEAST v2.6.2. Although XML files are available and a time-scaled phylogeny is plotted (their Figure 4), no tree file or description of divergence times are provided, and we therefore exclude this dataset.

**Scheel et al. (2014)*** *Phocidae.* Added CytB sequence data for the Neotropical monk seal *Neomonachus tropicalis* to the alignment of Fulton and Strobeck (2010) and jointly inferred the tree topology and divergence times using BEAST v1.7.4. They do not provide a tree file or a set of divergence time estimates for most nodes, but they do provide the latter for the node uniting the monk seals and the node uniting the two New World species. Accordingly, we assembled a 3-taxon tree from these two nodes.

**Trindade et al. (2021)*** *Felidae.* Generated a maximum likelihood tree from 60,931 SNPs to investigate the relationships of South American felids. They time-scaled this tree using PAML/MCMCTree under an autocorrelated-rates clock with a single calibration applied to the root node. There was no tree file available with the paper but we were able to manually digitize the topology and branch lengths present in Figure 1B of their paper.

**Tseng et al. (2014)** *Felidae.* Combined 43 kbp of molecular data with morphological data for 10 extant and recently extinct felid species with morphological data coded for those taxa plus an additional two fossil species and inferred a total-evidence tip-dated tree using MrBayes under a uniform tree prior and IGR clock. Although a MrBayes input file is available on MorphoBank (project number P898) there is no tree file and the divergence times are not provided in the paper. We therefore opted not to use this dataset.

**Westbury et al. (2021)*** *Hyaenidae.* Used ASTRAL to infer a species tree for the four extant species of Hyaenidae from maximum likelihood gene trees estimated using IQ-TREE. They then dated this tree using 3 fossil calibrations with soft bounds in the software PAML/MCMCTree. We took the tree topology from Figure 1 of their paper and posterior mean divergence times from the text. Outgroups were not included as dates are not available in the paper.

**Wolsan and Sato (2020)*** *Pinnipedia.* Used three taste receptor genes and six fossil constraints to sample time-calibrated phylogenies for 16 pinnipeds and 8 caniform outgroups. We transcribed the maximum likelihood tree shown in Supplementary Figure 5 and the posterior mean divergence time estimates from Supplementary Table 8.

**Table S13:** Composition of the 53 benchmark clades used to evaluate the performance of the empirical BLeSS analyses, ordered by taxonomic rank from most inclusive to least inclusive.

| Clade | Rank | Included species |
|---|---|---|
| Caniformia | Suborder | *Aenocyon dirus*, *Ailuropoda melanoleuca*, *Ailurus fulgens*, *Aonyx capensis*, *Aonyx cinerea*, *Arctocephalus australis*, *Arctocephalus forsteri*, *Arctocephalus galapagoensis*, *Arctocephalus gazella*, *Arctocephalus philippii*, *Arctocephalus pusillus*, *Arctocephalus townsendi*, *Arctocephalus tropicalis*, *Arctodus simus*, *Arctonyx collaris*, *Atelocynus microtis*, *Bassaricyon alleni*, *Bassaricyon gabbii*, *Bassaricyon medius*, *Bassaricyon neblina*, *Bassariscus astutus*, *Bassariscus sumichrasti*, *Callorhinus ursinus*, *Canis aureus*, *Canis latrans*, *Canis lupaster*, *Canis lupus*, *Canis rufus*, *Canis simensis*, *Cerdocyon thous*, *Chrysocyon brachyurus*, *Conepatus chinga*, *Conepatus leuconotus*, *Conepatus semistriatus*, *Cuon alpinus*, *Cystophora cristata*, *Dusicyon australis*, *Eira barbara*, *Enhydra lutris*, *Erignathus barbatus*, *Eumetopias jubatus*, *Galictis cuja*, *Galictis vittata*, *Gulo gulo*, *Halichoerus grypus*, *Helarctos malayanus*, *Histriophoca fasciata*, *Hydrictis maculicollis*, *Hydrurga leptonyx*, *Ictonyx libyca*, *Ictonyx striatus*, *Leptonychotes weddellii*, *Lobodon carcinophaga*, *Lontra canadensis*, *Lontra felina*, *Lontra longicaudis*, *Lontra provocax*, *Lupulella adusta*, *Lupulella mesomelas*, *Lutra lutra*, *Lutra sumatrana*, *Lutrogale perspicillata*, *Lycalopex culpaeus*, *Lycalopex fulvipes*, *Lycalopex griseus*, *Lycalopex gymnocercus*, *Lycalopex sechurae*, *Lycalopex vetulus*, *Lycaon pictus*, *Lyncodon patagonicus*, *Martes americana*, *Martes caurina*, *Martes flavigula*, *Martes foina*, *Martes martes*, *Martes melampus*, *Martes zibellina*, *Meles anakuma*, *Meles leucurus*, *Meles meles*, *Mellivora capensis*, *Melogale cucphuongensis*, *Melogale moschata*, *Melogale personata*, *Melursus ursinus*, *Mephitis macroura*, *Mephitis mephitis*, *Mirounga angustirostris*, *Mirounga leonina*, *Monachus monachus*, *Mustela altaica*, *Mustela erminea*, *Mustela eversmanni*, *Mustela itatsi*, *Mustela kathiah*, *Mustela lutreola*, *Mustela nigripes*, *Mustela nivalis*, *Mustela nudipes*, *Mustela putorius*, *Mustela sibirica*, *Mustela strigidorsa*, *Mydaus javanensis*, *Mydaus marchei*, *Nasua narica*, *Nasua nasua*, *Nasuella meridensis*, *Nasuella olivacea*, *Neogale africana*, *Neogale felipei*, *Neogale frenata*, *Neogale vison*, *Neomonachus schauinslandi*, *Neomonachus tropicalis*, *Neophoca cinerea*, *Nyctereutes procyonoides*, *Odobenus rosmarus*, *Ommatophoca rossii*, *Otaria byronia*, *Otaria flavescens*, *Otocyon megalotis*, *Pagophilus groenlandicus*, *Pekania pennanti*, *Phoca largha*, *Phoca vitulina*, *Phocarctos hookeri*, *Poecilogale albinucha*, *Potos flavus*, *Procyon cancrivorus*, *Procyon lotor*, *Pteronura brasiliensis*, *Pusa caspica*, *Pusa hispida*, *Pusa sibirica*, *Speothos venaticus*, *Spilogale angustifrons*, *Spilogale gracilis*, *Spilogale interrupta*, *Spilogale leucoparia*, *Spilogale putorius*, *Spilogale pygmaea*, *Spilogale* sp., *Spilogale yucatanensis*, *Taxidea taxus*, *Tremarctos ornatus*, *Urocyon cinereoargenteus*, *Urocyon littoralis*, *Ursus americanus*, *Ursus arctos*, *Ursus maritimus*, *Ursus spelaeus*, *Ursus thibetanus*, *Vormela peregusna*, *Vulpes bengalensis*, *Vulpes cana*, *Vulpes chama*, *Vulpes corsac*, *Vulpes ferrilata*, *Vulpes lagopus*, *Vulpes macrotis*, *Vulpes pallida*, *Vulpes rueppellii*, *Vulpes velox*, *Vulpes vulpes*, *Vulpes zerda*, *Zalophus californianus*, *Zalophus wollebaeki* |

| | | |
|---|---|---|
| Feliformia | Suborder | *Acinonyx jubatus*, *Arctictis binturong*, *Arctogalidia trivirgata*, *Caracal aurata*, *Caracal caracal*, *Catopuma badia*, *Catopuma temminckii*, *Chrotogale owstoni*, *Civettictis civetta*, *Crocuta crocuta*, *Crossarchus obscurus*, *Cryptoprocta ferox*, *Cynictis pencillata*, *Cynogale bennetti*, *Felis bieti*, *Felis chaus*, *Felis lybica*, *Felis margarita*, *Felis nigripes*, *Felis silvestris*, *Fossa fossana*, *Galidia elegans*, *Genetta angolensis*, *Genetta bourloni*, *Genetta felina*, *Genetta genetta*, *Genetta johnstoni*, *Genetta maculata*, *Genetta pardina*, *Genetta piscivora*, *Genetta poensis*, *Genetta schoutedeni*, *Genetta servalina*, *Genetta thierryi*, *Genetta tigrina*, *Genetta victoriae*, *Helogale parvula*, *Hemigalus derbyanus*, *Herpailurus yagouaroundi*, *Herpestes edwardsii*, *Herpestes ichneumon*, *Herpestes javanicus*, *Homotherium latidens*, *Hyaena hyaena*, *Ichneumia albicauda*, *Leopardus braccatus*, *Leopardus colocola*, *Leopardus garleppi*, *Leopardus geoffroyi*, *Leopardus guigna*, *Leopardus guttulus*, *Leopardus jacobita*, *Leopardus munoai*, *Leopardus pajeros*, *Leopardus pardalis*, *Leopardus tigrinus*, *Leopardus wiedii*, *Leptailurus serval*, *Lynx canadensis*, *Lynx lynx*, *Lynx pardinus*, *Lynx rufus*, *Mungos mungo*, *Mungotictis decemlineata*, *Nandinia binotata*, *Neofelis diardi*, *Neofelis nebulosa*, *Otocolobus manul*, *Paguma larvata*, *Panthera leo*, *Panthera onca*, *Panthera pardus*, *Panthera tigris*, *Panthera uncia*, *Paradoxurus hermaphroditus*, *Paradoxurus jerdoni*, *Parahyaena brunnea*, *Pardofelis marmorata*, *Poiana richardsonii*, *Prionailurus bengalensis*, *Prionailurus planiceps*, *Prionailurus rubiginosus*, *Prionailurus viverrinus*, *Prionodon linsang*, *Prionodon pardicolor*, *Proteles cristatus*, *Puma concolor*, *Rhynchogale melleri*, *Smilodon populator*, *Suricata suricatta*, *Viverra megaspila*, *Viverra tangalunga*, *Viverra zibetha*, *Viverricula indica* |
| Arctoidea | Infraorder | *Ailuropoda melanoleuca*, *Ailurus fulgens*, *Aonyx capensis*, *Aonyx cinerea*, *Arctocephalus australis*, *Arctocephalus forsteri*, *Arctocephalus galapagoensis*, *Arctocephalus gazella*, *Arctocephalus philippii*, *Arctocephalus pusillus*, *Arctocephalus townsendi*, *Arctocephalus tropicalis*, *Arctodus simus*, *Arctonyx collaris*, *Bassaricyon alleni*, *Bassaricyon gabbii*, *Bassaricyon medius*, *Bassaricyon neblina*, *Bassariscus astutus*, *Bassariscus sumichrasti*, *Callorhinus ursinus*, *Conepatus chinga*, *Conepatus leuconotus*, *Conepatus semistriatus*, *Cystophora cristata*, *Eira barbara*, *Enhydra lutris*, *Erignathus barbatus*, *Eumetopias jubatus*, *Galictis cuja*, *Galictis vittata*, *Gulo gulo*, *Halichoerus grypus*, *Helarctos malayanus*, *Histriophoca fasciata*, *Hydrictis maculicollis*, *Hydrurga leptonyx*, *Ictonyx libyca*, *Ictonyx striatus*, *Leptonychotes weddellii*, *Lobodon carcinophaga*, *Lontra canadensis*, *Lontra felina*, *Lontra longicaudis*, *Lontra provocax*, *Lutra lutra*, *Lutra sumatrana*, *Lutrogale perspicillata*, *Lyncodon patagonicus*, *Martes americana*, *Martes caurina*, *Martes flavigula*, *Martes foina*, *Martes martes*, *Martes melampus*, *Martes zibellina*, *Meles anakuma*, *Meles leucurus*, *Meles meles*, *Mellivora capensis*, *Melogale cucphuongensis*, *Melogale moschata*, *Melogale personata*, *Melursus ursinus*, *Mephitis macroura*, *Mephitis mephitis*, *Mirounga angustirostris*, *Mirounga leonina*, *Monachus monachus*, *Mustela altaica*, *Mustela erminea*, *Mustela eversmanni*, *Mustela itatsi*, *Mustela kathiah*, *Mustela lutreola*, *Mustela nigripes*, *Mustela nivalis*, *Mustela nudipes*, *Mustela putorius*, *Mustela sibirica*, *Mustela strigidorsa*, *Mydaus javanensis*, *Mydaus marchei*, *Nasua narica*, *Nasua nasua*, *Nasuella meridensis*, *Nasuella olivacea*, *Neogale africana*, *Neogale felipei*, *Neogale frenata*, *Neogale vison*, *Neomonachus schauinslandi*, *Neomonachus tropicalis*, *Neophoca cinerea*, *Odobenus rosmarus*, *Ommatophoca rossii*, *Otaria byronia*, *Otaria flavescens*, |

| | | |
|---|---|---|
| | | *Pagophilus groenlandicus*, *Pekania pennanti*, *Phoca largha*, *Phoca vitulina*, *Phocarctos hookeri*, *Poecilogale albinucha*, *Potos flavus*, *Procyon cancrivorus*, *Procyon lotor*, *Pteronura brasiliensis*, *Pusa caspica*, *Pusa hispida*, *Pusa sibirica*, *Spilogale angustifrons*, *Spilogale gracilis*, *Spilogale interrupta*, *Spilogale leucoparia*, *Spilogale putorius*, *Spilogale pygmaea*, *Spilogale* sp., *Spilogale yucatanensis*, *Taxidea taxus*, *Tremarctos ornatus*, *Ursus americanus*, *Ursus arctos*, *Ursus maritimus*, *Ursus spelaeus*, *Ursus thibetanus*, *Vormela peregusna*, *Zalophus californianus*, *Zalophus wollebaeki* |
| Viverroidea | Infraorder | *Arctictis binturong*, *Arctogalidia trivirgata*, *Chrotogale owstoni*, *Civettictis civetta*, *Crocuta crocuta*, *Crossarchus obscurus*, *Cryptoprocta ferox*, *Cynictis pencillata*, *Cynogale bennetti*, *Fossa fossana*, *Galidia elegans*, *Genetta angolensis*, *Genetta bourloni*, *Genetta felina*, *Genetta genetta*, *Genetta johnstoni*, *Genetta maculata*, *Genetta pardina*, *Genetta piscivora*, *Genetta poensis*, *Genetta schoutedeni*, *Genetta servalina*, *Genetta thierryi*, *Genetta tigrina*, *Genetta victoriae*, *Helogale parvula*, *Hemigalus derbyanus*, *Herpestes edwardsii*, *Herpestes ichneumon*, *Herpestes javanicus*, *Hyaena hyaena*, *Ichneumia albicauda*, *Mungos mungo*, *Mungotictis decemlineata*, *Paguma larvata*, *Paradoxurus hermaphroditus*, *Paradoxurus jerdoni*, *Parahyaena brunnea*, *Poiana richardsonii*, *Proteles cristatus*, *Rhynchogale melleri*, *Suricata suricatta*, *Viverra megaspila*, *Viverra tangalunga*, *Viverra zibetha*, *Viverricula indica* |
| Feloidea | Superfamily | *Acinonyx jubatus*, *Caracal aurata*, *Caracal caracal*, *Catopuma badia*, *Catopuma temminckii*, *Felis bieti*, *Felis chaus*, *Felis lybica*, *Felis margarita*, *Felis nigripes*, *Felis silvestris*, *Herpailurus yagouaroundi*, *Homotherium latidens*, *Leopardus braccatus*, *Leopardus colocola*, *Leopardus garleppi*, *Leopardus geoffroyi*, *Leopardus guigna*, *Leopardus guttulus*, *Leopardus jacobita*, *Leopardus munoai*, *Leopardus pajeros*, *Leopardus pardalis*, *Leopardus tigrinus*, *Leopardus wiedii*, *Leptailurus serval*, *Lynx canadensis*, *Lynx lynx*, *Lynx pardinus*, *Lynx rufus*, *Neofelis diardi*, *Neofelis nebulosa*, *Otocolobus manul*, *Panthera leo*, *Panthera onca*, *Panthera pardus*, *Panthera tigris*, *Panthera uncia*, *Pardofelis marmorata*, *Prionailurus bengalensis*, *Prionailurus planiceps*, *Prionailurus rubiginosus*, *Prionailurus viverrinus*, *Prionodon linsang*, *Prionodon pardicolor*, *Puma concolor*, *Smilodon populator* |
| Herpestoidea | Superfamily | *Crocuta crocuta*, *Crossarchus obscurus*, *Cryptoprocta ferox*, *Cynictis pencillata*, *Fossa fossana*, *Galidia elegans*, *Helogale parvula*, *Herpestes edwardsii*, *Herpestes ichneumon*, *Herpestes javanicus*, *Hyaena hyaena*, *Ichneumia albicauda*, *Mungos mungo*, *Mungotictis decemlineata*, *Parahyaena brunnea*, *Proteles cristatus*, *Rhynchogale melleri*, *Suricata suricatta* |
| Musteloidea | Superfamily | *Ailurus fulgens*, *Aonyx capensis*, *Aonyx cinerea*, *Arctonyx collaris*, *Bassaricyon alleni*, *Bassaricyon gabbii*, *Bassaricyon medius*, *Bassaricyon neblina*, *Bassariscus astutus*, *Bassariscus sumichrasti*, *Conepatus chinga*, *Conepatus leuconotus*, *Conepatus semistriatus*, *Eira barbara*, *Enhydra lutris*, *Galictis cuja*, *Galictis vittata*, *Gulo gulo*, *Hydrictis maculicollis*, *Ictonyx libyca*, *Ictonyx striatus*, *Lontra canadensis*, *Lontra felina*, *Lontra longicaudis*, *Lontra provocax*, *Lutra lutra*, *Lutra sumatrana*, *Lutrogale perspicillata*, *Lyncodon patagonicus*, *Martes americana*, *Martes caurina*, *Martes flavigula*, *Martes foina*, *Martes martes*, *Martes melampus*, *Martes zibellina*, *Meles anakuma*, *Meles leucurus*, *Meles meles*, *Mellivora capensis*, *Melogale cucphuongensis*, *Melogale moschata*, *Melogale personata*, *Mephitis macroura*, *Mephitis mephitis*, *Mustela altaica*, |

| | | |
|---|---|---|
| | | *Mustela erminea*, *Mustela eversmanni*, *Mustela itatsi*, *Mustela kathiah*, *Mustela lutreola*, *Mustela nigripes*, *Mustela nivalis*, *Mustela nudipes*, *Mustela putorius*, *Mustela sibirica*, *Mustela strigidorsa*, *Mydaus javanensis*, *Mydaus marchei*, *Nasua narica*, *Nasua nasua*, *Nasuella meridensis*, *Nasuella olivacea*, *Neogale africana*, *Neogale felipei*, *Neogale frenata*, *Neogale vison*, *Pekania pennanti*, *Poecilogale albinucha*, *Potos flavus*, *Procyon cancrivorus*, *Procyon lotor*, *Pteronura brasiliensis*, *Spilogale angustifrons*, *Spilogale gracilis*, *Spilogale interrupta*, *Spilogale leucoparia*, *Spilogale putorius*, *Spilogale pygmaea*, *Spilogale* sp., *Spilogale yucatanensis*, *Taxidea taxus*, *Vormela peregusna* |
| Pinnipedia | Superfamily | *Arctocephalus australis*, *Arctocephalus forsteri*, *Arctocephalus galapagoensis*, *Arctocephalus gazella*, *Arctocephalus philippii*, *Arctocephalus pusillus*, *Arctocephalus townsendi*, *Arctocephalus tropicalis*, *Callorhinus ursinus*, *Cystophora cristata*, *Erignathus barbatus*, *Eumetopias jubatus*, *Halichoerus grypus*, *Histriophoca fasciata*, *Hydrurga leptonyx*, *Leptonychotes weddellii*, *Lobodon carcinophaga*, *Mirounga angustirostris*, *Mirounga leonina*, *Monachus monachus*, *Neomonachus schauinslandi*, *Neomonachus tropicalis*, *Neophoca cinerea*, *Odobenus rosmarus*, *Ommatophoca rossii*, *Otaria byronia*, *Otaria flavescens*, *Pagophilus groenlandicus*, *Phoca largha*, *Phoca vitulina*, *Phocarctos hookeri*, *Pusa caspica*, *Pusa hispida*, *Pusa sibirica*, *Zalophus californianus*, *Zalophus wollebaeki* |
| Canidae | Family | *Aenocyon dirus*, *Atelocynus microtis*, *Canis aureus*, *Canis latrans*, *Canis lupaster*, *Canis lupus*, *Canis rufus*, *Canis simensis*, *Cerdocyon thous*, *Chrysocyon brachyurus*, *Cuon alpinus*, *Dusicyon australis*, *Lupulella adusta*, *Lupulella mesomelas*, *Lycalopex culpaeus*, *Lycalopex fulvipes*, *Lycalopex griseus*, *Lycalopex gymnocercus*, *Lycalopex sechurae*, *Lycalopex vetulus*, *Lycaon pictus*, *Nyctereutes procyonoides*, *Otocyon megalotis*, *Speothos venaticus*, *Urocyon cinereoargenteus*, *Urocyon littoralis*, *Vulpes bengalensis*, *Vulpes cana*, *Vulpes chama*, *Vulpes corsac*, *Vulpes ferrilata*, *Vulpes lagopus*, *Vulpes macrotis*, *Vulpes pallida*, *Vulpes rueppellii*, *Vulpes velox*, *Vulpes vulpes*, *Vulpes zerda* |
| Eupleridae | Family | *Cryptoprocta ferox*, *Fossa fossana*, *Galidia elegans*, *Mungotictis decemlineata* |
| Felidae | Family | *Acinonyx jubatus*, *Caracal aurata*, *Caracal caracal*, *Catopuma badia*, *Catopuma temminckii*, *Felis bieti*, *Felis chaus*, *Felis lybica*, *Felis margarita*, *Felis nigripes*, *Felis silvestris*, *Herpailurus yagouaroundi*, *Homotherium latidens*, *Leopardus braccatus*, *Leopardus colocola*, *Leopardus garleppi*, *Leopardus geoffroyi*, *Leopardus guigna*, *Leopardus guttulus*, *Leopardus jacobita*, *Leopardus munoai*, *Leopardus pajeros*, *Leopardus pardalis*, *Leopardus tigrinus*, *Leopardus wiedii*, *Leptailurus serval*, *Lynx canadensis*, *Lynx lynx*, *Lynx pardinus*, *Lynx rufus*, *Neofelis diardi*, *Neofelis nebulosa*, *Otocolobus manul*, *Panthera leo*, *Panthera onca*, *Panthera pardus*, *Panthera tigris*, *Panthera uncia*, *Pardofelis marmorata*, *Prionailurus bengalensis*, *Prionailurus planiceps*, *Prionailurus rubiginosus*, *Prionailurus viverrinus*, *Puma concolor*, *Smilodon populator* |
| Herpestidae | Family | *Crossarchus obscurus*, *Cynictis pencillata*, *Helogale parvula*, *Herpestes edwardsii*, *Herpestes ichneumon*, *Herpestes javanicus*, *Ichneumia albicauda*, *Mungos mungo*, *Rhynchogale melleri*, *Suricata suricatta* |
| Hyaenidae | Family | *Crocuta crocuta*, *Hyaena hyaena*, *Parahyaena brunnea*, *Proteles cristatus* |
| Mephitidae | Family | *Conepatus chinga*, *Conepatus leuconotus*, *Conepatus semistriatus*, *Mephitis macroura*, *Mephitis mephitis*, *Mydaus javanensis*, *Mydaus marchei*, *Spilogale angustifrons*, *Spilogale gracilis*, *Spilogale interrupta*, *Spilogale leucoparia*, *Spilogale putorius*, *Spilogale pygmaea*, *Spilogale* sp., *Spilogale yucatanensis* |

| | | |
|---|---|---|
| Mustelidae | Family | *Aonyx capensis*, *Aonyx cinerea*, *Arctonyx collaris*, *Eira barbara*, *Enhydra lutris*, *Galictis cuja*, *Galictis vittata*, *Gulo gulo*, *Hydrictis maculicollis*, *Ictonyx libyca*, *Ictonyx striatus*, *Lontra canadensis*, *Lontra felina*, *Lontra longicaudis*, *Lontra provocax*, *Lutra lutra*, *Lutra sumatrana*, *Lutrogale perspicillata*, *Lyncodon patagonicus*, *Martes americana*, *Martes caurina*, *Martes flavigula*, *Martes foina*, *Martes martes*, *Martes melampus*, *Martes zibellina*, *Meles anakuma*, *Meles leucurus*, *Meles meles*, *Mellivora capensis*, *Melogale cucphuongensis*, *Melogale moschata*, *Melogale personata*, *Mustela altaica*, *Mustela erminea*, *Mustela eversmanni*, *Mustela itatsi*, *Mustela kathiah*, *Mustela lutreola*, *Mustela nigripes*, *Mustela nivalis*, *Mustela nudipes*, *Mustela putorius*, *Mustela sibirica*, *Mustela strigidorsa*, *Neogale africana*, *Neogale felipei*, *Neogale frenata*, *Neogale vison*, *Pekania pennanti*, *Poecilogale albinucha*, *Pteronura brasiliensis*, *Taxidea taxus*, *Vormela peregusna* |
| Otariidae | Family | *Arctocephalus australis*, *Arctocephalus forsteri*, *Arctocephalus galapagoensis*, *Arctocephalus gazella*, *Arctocephalus philippii*, *Arctocephalus pusillus*, *Arctocephalus townsendi*, *Arctocephalus tropicalis*, *Callorhinus ursinus*, *Eumetopias jubatus*, *Neophoca cinerea*, *Otaria byronia*, *Otaria flavescens*, *Phocarctos hookeri*, *Zalophus californianus*, *Zalophus wollebaeki* |
| Phocidae | Family | *Cystophora cristata*, *Erignathus barbatus*, *Halichoerus grypus*, *Histriophoca fasciata*, *Hydrurga leptonyx*, *Leptonychotes weddellii*, *Lobodon carcinophaga*, *Mirounga angustirostris*, *Mirounga leonina*, *Monachus monachus*, *Neomonachus schauinslandi*, *Neomonachus tropicalis*, *Ommatophoca rossii*, *Pagophilus groenlandicus*, *Phoca largha*, *Phoca vitulina*, *Pusa caspica*, *Pusa hispida*, *Pusa sibirica* |
| Prionodontidae | Family | *Prionodon linsang*, *Prionodon pardicolor* |
| Procyonidae | Family | *Bassaricyon alleni*, *Bassaricyon gabbii*, *Bassaricyon medius*, *Bassaricyon neblina*, *Bassariscus astutus*, *Bassariscus sumichrasti*, *Nasua narica*, *Nasua nasua*, *Nasuella meridensis*, *Nasuella olivacea*, *Potos flavus*, *Procyon cancrivorus*, *Procyon lotor* |
| Ursidae | Family | *Ailuropoda melanoleuca*, *Arctodus simus*, *Helarctos malayanus*, *Melursus ursinus*, *Tremarctos ornatus*, *Ursus americanus*, *Ursus arctos*, *Ursus maritimus*, *Ursus spelaeus*, *Ursus thibetanus* |
| Viverridae | Family | *Arctictis binturong*, *Arctogalidia trivirgata*, *Chrotogale owstoni*, *Civettictis civetta*, *Cynogale bennetti*, *Genetta angolensis*, *Genetta bourloni*, *Genetta felina*, *Genetta genetta*, *Genetta johnstoni*, *Genetta maculata*, *Genetta pardina*, *Genetta piscivora*, *Genetta poensis*, *Genetta schoutedeni*, *Genetta servalina*, *Genetta thierryi*, *Genetta tigrina*, *Genetta victoriae*, *Hemigalus derbyanus*, *Paguma larvata*, *Paradoxurus hermaphroditus*, *Paradoxurus jerdoni*, *Poiana richardsonii*, *Viverra megaspila*, *Viverra tangalunga*, *Viverra zibetha*, *Viverricula indica* |
| Caninae | Subfamily | *Aenocyon dirus*, *Atelocynus microtis*, *Canis aureus*, *Canis latrans*, *Canis lupaster*, *Canis lupus*, *Canis rufus*, *Canis simensis*, *Cerdocyon thous*, *Chrysocyon brachyurus*, *Cuon alpinus*, *Dusicyon australis*, *Lupulella adusta*, *Lupulella mesomelas*, *Lycalopex culpaeus*, *Lycalopex fulvipes*, *Lycalopex griseus*, *Lycalopex gymnocercus*, *Lycalopex sechurae*, *Lycalopex vetulus*, *Lycaon pictus*, *Nyctereutes procyonoides*, *Otocyon megalotis*, *Speothos venaticus*, *Urocyon cinereoargenteus*, *Urocyon littoralis*, *Vulpes bengalensis*, *Vulpes cana*, *Vulpes chama*, *Vulpes corsac*, *Vulpes ferrilata*, *Vulpes lagopus*, *Vulpes macrotis*, *Vulpes pallida*, *Vulpes rueppellii*, *Vulpes velox*, *Vulpes vulpes*, *Vulpes zerda* |
| Euplerinae | Subfamily | *Cryptoprocta ferox*, *Fossa fossana* |

| | | |
|---|---|---|
| Felinae | Subfamily | *Acinonyx jubatus*, *Caracal aurata*, *Caracal caracal*, *Catopuma badia*, *Catopuma temminckii*, *Felis bieti*, *Felis chaus*, *Felis lybica*, *Felis margarita*, *Felis nigripes*, *Felis silvestris*, *Herpailurus yagouaroundi*, *Leopardus braccatus*, *Leopardus colocola*, *Leopardus garleppi*, *Leopardus geoffroyi*, *Leopardus guigna*, *Leopardus guttulus*, *Leopardus jacobita*, *Leopardus munoai*, *Leopardus pajeros*, *Leopardus pardalis*, *Leopardus tigrinus*, *Leopardus wiedii*, *Leptailurus serval*, *Lynx canadensis*, *Lynx lynx*, *Lynx pardinus*, *Lynx rufus*, *Neofelis diardi*, *Neofelis nebulosa*, *Otocolobus manul*, *Panthera leo*, *Panthera onca*, *Panthera pardus*, *Panthera tigris*, *Panthera uncia*, *Pardofelis marmorata*, *Prionailurus bengalensis*, *Prionailurus planiceps*, *Prionailurus rubiginosus*, *Prionailurus viverrinus*, *Puma concolor* |
| Galidinae | Subfamily | *Galidia elegans*, *Mungotictis decemlineata* |
| Genettinae | Subfamily | *Genetta angolensis*, *Genetta bourloni*, *Genetta felina*, *Genetta genetta*, *Genetta johnstoni*, *Genetta maculata*, *Genetta pardina*, *Genetta piscivora*, *Genetta poensis*, *Genetta schoutedeni*, *Genetta servalina*, *Genetta thierryi*, *Genetta tigrina*, *Genetta victoriae*, *Poiana richardsonii* |
| Guloninae | Subfamily | *Eira barbara*, *Gulo gulo*, *Martes americana*, *Martes caurina*, *Martes flavigula*, *Martes foina*, *Martes martes*, *Martes melampus*, *Martes zibellina*, *Pekania pennanti* |
| Helictidinae | Subfamily | *Melogale cucphuongensis*, *Melogale moschata*, *Melogale personata* |
| Hemigalinae | Subfamily | *Chrotogale owstoni*, *Cynogale bennetti*, *Hemigalus derbyanus* |
| Herpestinae | Subfamily | *Cynictis pencillata*, *Herpestes edwardsii*, *Herpestes ichneumon*, *Herpestes javanicus*, *Ichneumia albicauda*, *Rhynchogale melleri* |
| Ictonychinae | Subfamily | *Galictis cuja*, *Galictis vittata*, *Ictonyx libyca*, *Ictonyx striatus*, *Lyncodon patagonicus*, *Poecilogale albinucha*, *Vormela peregusna* |
| Lutrinae | Subfamily | *Aonyx capensis*, *Aonyx cinerea*, *Enhydra lutris*, *Hydrictis maculicollis*, *Lontra canadensis*, *Lontra felina*, *Lontra longicaudis*, *Lontra provocax*, *Lutra lutra*, *Lutra sumatrana*, *Lutrogale perspicillata*, *Pteronura brasiliensis* |
| Machairodontinae | Subfamily | *Homotherium latidens*, *Smilodon populator* |
| Melinae | Subfamily | *Arctonyx collaris*, *Meles anakuma*, *Meles leucurus*, *Meles meles* |
| Monachinae | Subfamily | *Hydrurga leptonyx*, *Leptonychotes weddellii*, *Lobodon carcinophaga*, *Mirounga angustirostris*, *Mirounga leonina*, *Monachus monachus*, *Neomonachus schauinslandi*, *Neomonachus tropicalis*, *Ommatophoca rossii* |
| Mungotinae | Subfamily | *Crossarchus obscurus*, *Helogale parvula*, *Mungos mungo*, *Suricata suricatta* |
| Mustelinae | Subfamily | *Mustela altaica*, *Mustela erminea*, *Mustela eversmanni*, *Mustela itatsi*, *Mustela kathiah*, *Mustela lutreola*, *Mustela nigripes*, *Mustela nivalis*, *Mustela nudipes*, *Mustela putorius*, *Mustela sibirica*, *Mustela strigidorsa*, *Neogale africana*, *Neogale felipei*, *Neogale frenata*, *Neogale vison* |
| Paradoxurinae | Subfamily | *Arctictis binturong*, *Arctogalidia trivirgata*, *Paguma larvata*, *Paradoxurus hermaphroditus*, *Paradoxurus jerdoni* |
| Phocinae | Subfamily | *Cystophora cristata*, *Erignathus barbatus*, *Halichoerus grypus*, *Histriophoca fasciata*, *Pagophilus groenlandicus*, *Phoca largha*, *Phoca vitulina*, *Pusa caspica*, *Pusa hispida*, *Pusa sibirica* |
| Ursinae | Subfamily | *Ailuropoda melanoleuca*, *Arctodus simus*, *Helarctos malayanus*, *Melursus ursinus*, *Tremarctos ornatus*, *Ursus americanus*, *Ursus arctos*, *Ursus maritimus*, *Ursus spelaeus*, *Ursus thibetanus* |
| Viverrinae | Subfamily | *Civettictis civetta*, *Viverra megaspila*, *Viverra tangalunga*, *Viverra zibetha*, *Viverricula indica* |
| Arctotheriini | Tribe | *Arctodus simus*, *Tremarctos ornatus* |

| | | |
|---|---|---|
| Canini | Tribe | *Aenocyon dirus*, *Atelocynus microtis*, *Canis aureus*, *Canis latrans*, *Canis lupaster*, *Canis lupus*, *Canis rufus*, *Canis simensis*, *Cerdocyon thous*, *Chrysocyon brachyurus*, *Cuon alpinus*, *Dusicyon australis*, *Lupulella adusta*, *Lupulella mesomelas*, *Lycalopex culpaeus*, *Lycalopex fulvipes*, *Lycalopex griseus*, *Lycalopex gymnocercus*, *Lycalopex sechurae*, *Lycalopex vetulus*, *Lycaon pictus*, *Speothos venaticus* |
| Felini | Tribe | *Acinonyx jubatus*, *Caracal aurata*, *Caracal caracal*, *Catopuma badia*, *Catopuma temminckii*, *Felis bieti*, *Felis chaus*, *Felis lybica*, *Felis margarita*, *Felis nigripes*, *Felis silvestris*, *Herpailurus yagouaroundi*, *Leopardus braccatus*, *Leopardus colocola*, *Leopardus garleppi*, *Leopardus geoffroyi*, *Leopardus guigna*, *Leopardus guttulus*, *Leopardus jacobita*, *Leopardus munoai*, *Leopardus pajeros*, *Leopardus pardalis*, *Leopardus tigrinus*, *Leopardus wiedii*, *Leptailurus serval*, *Lynx canadensis*, *Lynx lynx*, *Lynx pardinus*, *Lynx rufus*, *Otocolobus manul*, *Pardofelis marmorata*, *Prionailurus bengalensis*, *Prionailurus planiceps*, *Prionailurus rubiginosus*, *Prionailurus viverrinus*, *Puma concolor* |
| Lobodontini | Tribe | *Hydrurga leptonyx*, *Leptonychotes weddellii*, *Lobodon carcinophaga*, *Ommatophoca rossii* |
| Miroungini | Tribe | *Mirounga angustirostris*, *Mirounga leonina* |
| Monachini | Tribe | *Monachus monachus*, *Neomonachus schauinslandi*, *Neomonachus tropicalis* |
| Pantherini | Tribe | *Neofelis diardi*, *Neofelis nebulosa*, *Panthera leo*, *Panthera onca*, *Panthera pardus*, *Panthera tigris*, *Panthera uncia* |
| Phocini | Tribe | *Halichoerus grypus*, *Histriophoca fasciata*, *Pagophilus groenlandicus*, *Phoca largha*, *Phoca vitulina*, *Pusa caspica*, *Pusa hispida*, *Pusa sibirica* |
| Ursini | Tribe | *Helarctos malayanus*, *Melursus ursinus*, *Ursus americanus*, *Ursus arctos*, *Ursus maritimus*, *Ursus spelaeus*, *Ursus thibetanus* |
| Vulpini | Tribe | *Nyctereutes procyonoides*, *Otocyon megalotis*, *Urocyon cinereoargenteus*, *Urocyon littoralis*, *Vulpes bengalensis*, *Vulpes cana*, *Vulpes chama*, *Vulpes corsac*, *Vulpes ferrilata*, *Vulpes lagopus*, *Vulpes macrotis*, *Vulpes pallida*, *Vulpes rueppellii*, *Vulpes velox*, *Vulpes vulpes*, *Vulpes zerda* |
| Canina | Subtribe | *Aenocyon dirus*, *Canis aureus*, *Canis latrans*, *Canis lupaster*, *Canis lupus*, *Canis rufus*, *Canis simensis*, *Cuon alpinus*, *Lupulella adusta*, *Lupulella mesomelas*, *Lycaon pictus* |
| Cerdocyonina | Subtribe | *Atelocynus microtis*, *Cerdocyon thous*, *Chrysocyon brachyurus*, *Dusicyon australis*, *Lycalopex culpaeus*, *Lycalopex fulvipes*, *Lycalopex griseus*, *Lycalopex gymnocercus*, *Lycalopex sechurae*, *Lycalopex vetulus*, *Speothos venaticus* |

**Table S14:** Mean posterior root ages and topological accuracy (as quantified by the number of recovered benchmark clades from Table S13) of the summary carnivoran supertrees inferred with BLeSS under different analytical settings, as well as of the time-scaled supertree of Nyakatura and Bininda-Emonds (2012) ("N&BE12"), the backbone-and-patch time tree of Upham et al. (2019) ("UEA19"), and the supermatrix-based time tree of Slater and Friscia (2019) ("S&F19"). Due to differences in taxon sampling, not all trees could be scored for the same set of benchmark clades. Downw. = downweighting of mtDNA-based source trees; Est. = estimated; MAP = maximum *a posteriori* tree; MCC = maximum clade credibility tree; HL run = MCMC run with the highest post-burnin likelihoods. Note that the MAP and MCC trees were only computed from the pooled posterior sample of all 4 MCMC runs if the average standard deviation of split frequencies did not exceed 0.01; otherwise, only the highest-likelihood run was summarized.

| Source | BLeSS settings | | | | Root age | Benchmark clades | | Success |
|---|---|---|---|---|---|---|---|---|
| | Imputation | Downw. | $\lambda_e$ | Summary tree | (Ma) | Recovered | Scored | rate |
| This study | No | No | 0.1 | MAP, all runs | 49.301 | 16 | 53 | 0.302 |
| This study | No | No | 0.1 | MCC, all runs | 49.301 | 16 | 53 | 0.302 |
| This study | No | No | 0.1 | MAP, HL run | 49.301 | 16 | 53 | 0.302 |
| This study | No | No | 0.1 | MCC, HL run | 49.301 | 17 | 53 | 0.321 |
| This study | No | Yes | 0.1 | MAP, all runs | 49.658 | 31 | 53 | 0.585 |
| This study | No | Yes | 0.1 | MCC, all runs | 49.658 | 19 | 53 | 0.358 |
| This study | No | Yes | 0.1 | MAP, HL run | 56.118 | 31 | 53 | 0.585 |
| This study | No | Yes | 0.1 | MCC, HL run | 56.118 | 31 | 53 | 0.585 |
| This study | Yes | No | 0.1 | MAP, HL run | 51.265 | 45 | 53 | 0.849 |
| This study | Yes | No | 0.1 | MCC, HL run | 51.265 | 45 | 53 | 0.849 |
| This study | Yes | Yes | 0.1 | MAP, HL run | 51.265 | 45 | 53 | 0.849 |
| This study | Yes | Yes | 0.1 | MCC, HL run | 51.265 | 46 | 53 | 0.868 |
| This study | Yes | Yes | Est. | MAP, all runs | 43.374 | 0 | 53 | 0 |
| This study | Yes | Yes | Est. | MCC, all runs | 43.374 | 0 | 53 | 0 |
| N&BE12 | N/A | N/A | N/A | N/A | 64.9 | 42 | 51 | 0.824 |
| UEA19 | N/A | N/A | N/A | N/A | 40.795 | 48 | 52 | 0.923 |
| S&F19 | N/A | N/A | N/A | N/A | 48.186 | 43 | 44 | 0.977 |

**Table S15:** Pairwise tree distances calculated for the carnivoran phylogenies of Nyakatura and Bininda-Emonds (2012) ("N&BE12") and Upham et al. (2019) ("UEA19") relative to the best BLeSS estimate (imputation, downweighting of mtDNA-based source trees, MCC supertree generated from the highest-likelihood run only) and the supermatrix-based phylogeny of Slater and Friscia (2019) ("S&F19"), which exhibited the highest proportion of recovered benchmark clades (Table S14). Prior to carrying out the comparison, each pair of trees was pruned down to the largest common subset of tips. Note that the N&BE12 tree contained polytomies, which were stochastically resolved to allow calculating those distance metrics that are only defined for binary trees. The Robinson-Foulds distance, path length difference, clustering information distance, and symmetric quartet divergence are normalized. $\lambda$ denotes the contribution of branch lengths (as opposed to topology) to the Kendall-Colijn metric.

| Metric | N&BE12 vs. BLeSS | UEA19 vs. BLeSS | S&F19 vs. BLeSS | N&BE12 vs. S&F19 | UEA19 vs. S&F19 |
|---|---|---|---|---|---|
| Shared tips | 237 | 236 | 206 | 196 | 198 |
| Robinson-Foulds | 0.4615 | 0.412 | 0.3547 | 0.3212 | 0.1846 |
| Kuhner-Felsenstein | 86.54 | 60.02 | 50.52 | 80.97 | 29.26 |
| Path length difference | 0.2376 | 0.2427 | 0.134 | 0.2854 | 0.1937 |
| Billera-Holmes-Vogtmann | 93.77 | 62.48 | 53.4 | 88.36 | 30.24 |
| Matching split | 565 | 497 | 411 | 253 | 149 |
| Clustering information | 0.8085 | 0.83 | 0.8401 | 0.8741 | 0.9214 |
| Symmetric quartet divergence | 0.9445 | 0.9442 | 0.9495 | 0.9881 | 0.9941 |
| Kendall-Colijn, $\lambda = 0$ | 157.11 | 150.05 | 114.48 | 74.89 | 40.37 |
| Kendall-Colijn, $\lambda = 0.1$ | 216.31 | 130.09 | 108.89 | 141.72 | 52.65 |
| Kendall-Colijn, $\lambda = 0.2$ | 295.11 | 137.61 | 128.49 | 235.79 | 81.91 |
| Kendall-Colijn, $\lambda = 0.3$ | 381.55 | 168.98 | 164.5 | 334.88 | 115.95 |
| Kendall-Colijn, $\lambda = 0.4$ | 471.45 | 213.96 | 208.59 | 435.58 | 151.57 |
| Kendall-Colijn, $\lambda = 0.5$ | 563.15 | 265.71 | 256.63 | 536.98 | 187.88 |
| Kendall-Colijn, $\lambda = 0.6$ | 655.9 | 320.99 | 306.76 | 638.75 | 224.55 |
| Kendall-Colijn, $\lambda = 0.7$ | 749.31 | 378.25 | 358.12 | 740.74 | 261.42 |
| Kendall-Colijn, $\lambda = 0.8$ | 843.15 | 436.7 | 410.23 | 842.86 | 298.42 |
| Kendall-Colijn, $\lambda = 0.9$ | 937.31 | 495.94 | 462.85 | 945.08 | 335.51 |
| Kendall-Colijn, $\lambda = 1$ | 1031.69 | 555.7 | 515.82 | 1047.36 | 372.65 |
| Root age difference (myr) | 13.63 | −10.47 | −3.08 | 16.71 | −7.39 |

**Table S16:** Pairwise tree distances calculated for the carnivoran phylogenies of Nyakatura and Bininda-Emonds (2012) ("N&BE12") and Upham et al. (2019) ("UEA19") relative to the best BLeSS estimate and the phylogeny of Slater and Friscia (2019) ("S&F19") after pruning each tree down to the largest subset of tips common to all four. All distances calculated and reported as in Table S15.

| Metric | N&BE12 vs. BLeSS | UEA19 vs. BLeSS | S&F19 vs. BLeSS | N&BE12 vs. S&F19 | UEA19 vs. S&F19 |
|---|---|---|---|---|---|
| Shared tips | 192 | 192 | 192 | 192 | 192 |
| Robinson-Foulds | 0.4603 | 0.3968 | 0.3439 | 0.3228 | 0.1905 |
| Kuhner-Felsenstein | 84.23 | 55.61 | 48.75 | 80.91 | 29.39 |
| Path length difference | 0.2459 | 0.2416 | 0.1334 | 0.2822 | 0.1926 |
| Billera-Holmes-Vogtmann | 91.34 | 57.46 | 51.14 | 88.3 | 30.36 |
| Matching split | 424 | 361 | 367 | 247 | 148 |
| Clustering information | 0.8113 | 0.8361 | 0.8407 | 0.8732 | 0.9192 |
| Symmetric quartet divergence | 0.9473 | 0.9488 | 0.9505 | 0.9877 | 0.9936 |
| Kendall-Colijn, $\lambda = 0$ | 112.48 | 103.46 | 103.04 | 74.03 | 40.3 |
| Kendall-Colijn, $\lambda = 0.1$ | 163.9 | 101.19 | 99.22 | 139.75 | 52.1 |
| Kendall-Colijn, $\lambda = 0.2$ | 234.55 | 124.49 | 119.85 | 232.46 | 80.53 |
| Kendall-Colijn, $\lambda = 0.3$ | 311.62 | 162.74 | 155.48 | 330.16 | 113.74 |
| Kendall-Colijn, $\lambda = 0.4$ | 391.33 | 207.83 | 198.18 | 429.45 | 148.55 |
| Kendall-Colijn, $\lambda = 0.5$ | 472.35 | 256.18 | 244.27 | 529.45 | 184.07 |
| Kendall-Colijn, $\lambda = 0.6$ | 554.1 | 306.24 | 292.14 | 629.81 | 219.94 |
| Kendall-Colijn, $\lambda = 0.7$ | 636.31 | 357.31 | 341.06 | 730.39 | 256.02 |
| Kendall-Colijn, $\lambda = 0.8$ | 718.8 | 408.99 | 390.62 | 831.1 | 292.23 |
| Kendall-Colijn, $\lambda = 0.9$ | 801.51 | 461.09 | 440.61 | 931.91 | 328.53 |
| Kendall-Colijn, $\lambda = 1$ | 884.36 | 513.48 | 490.9 | 1032.78 | 364.9 |
| Root age difference (myr) | 13.63 | −10.47 | −3.08 | 16.71 | −7.39 |

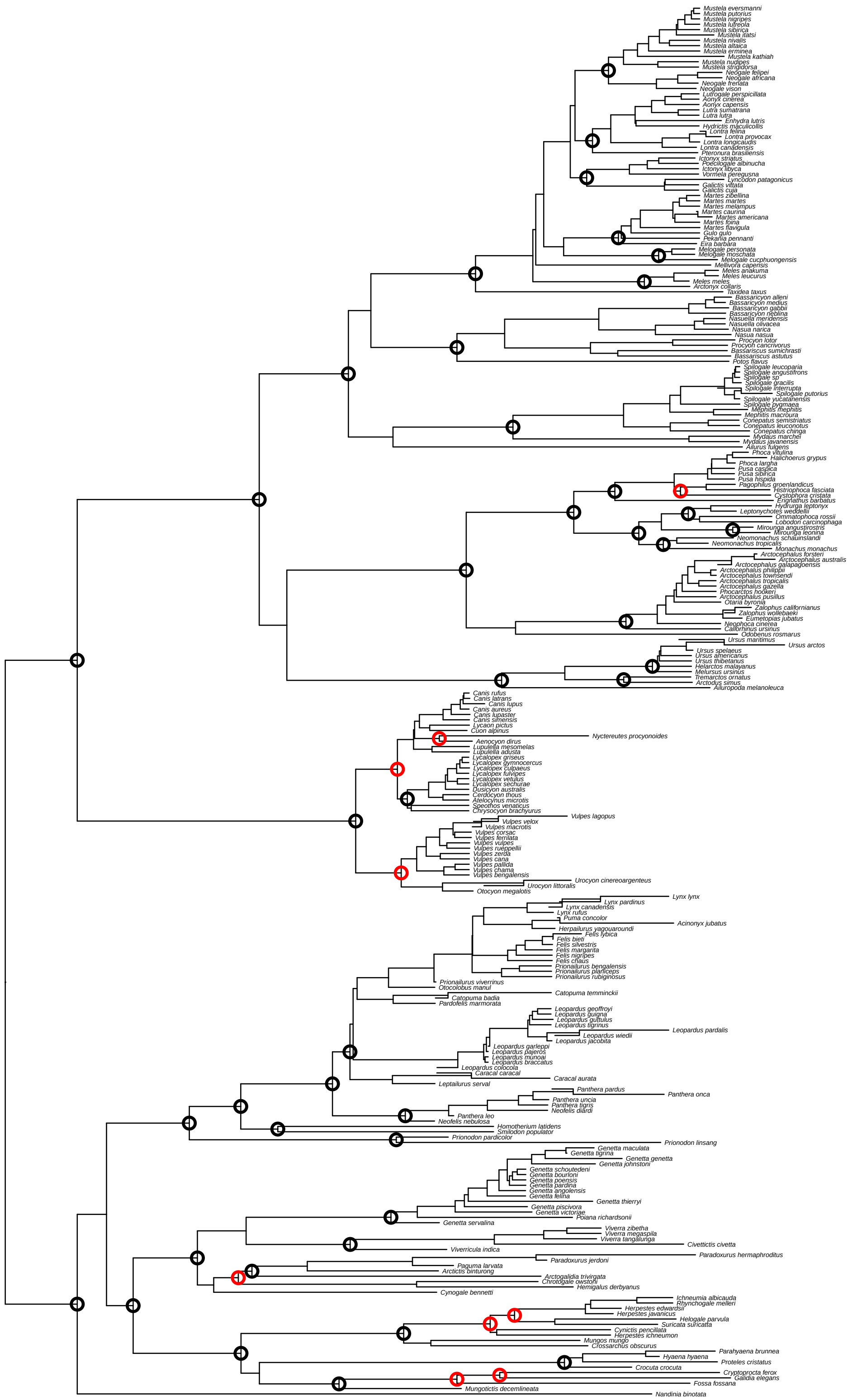

**Figure S26:** Neighbor-joining supertree of Carnivora inferred from an average distance matrix with imputed missing entries and all source trees weighted equally. The circles indicate nodes congruent with (black) or contradicting (red) the 53 predefined benchmark clades. Note that the tree is not ultrametric, rendering the branch lengths uninterpretable.

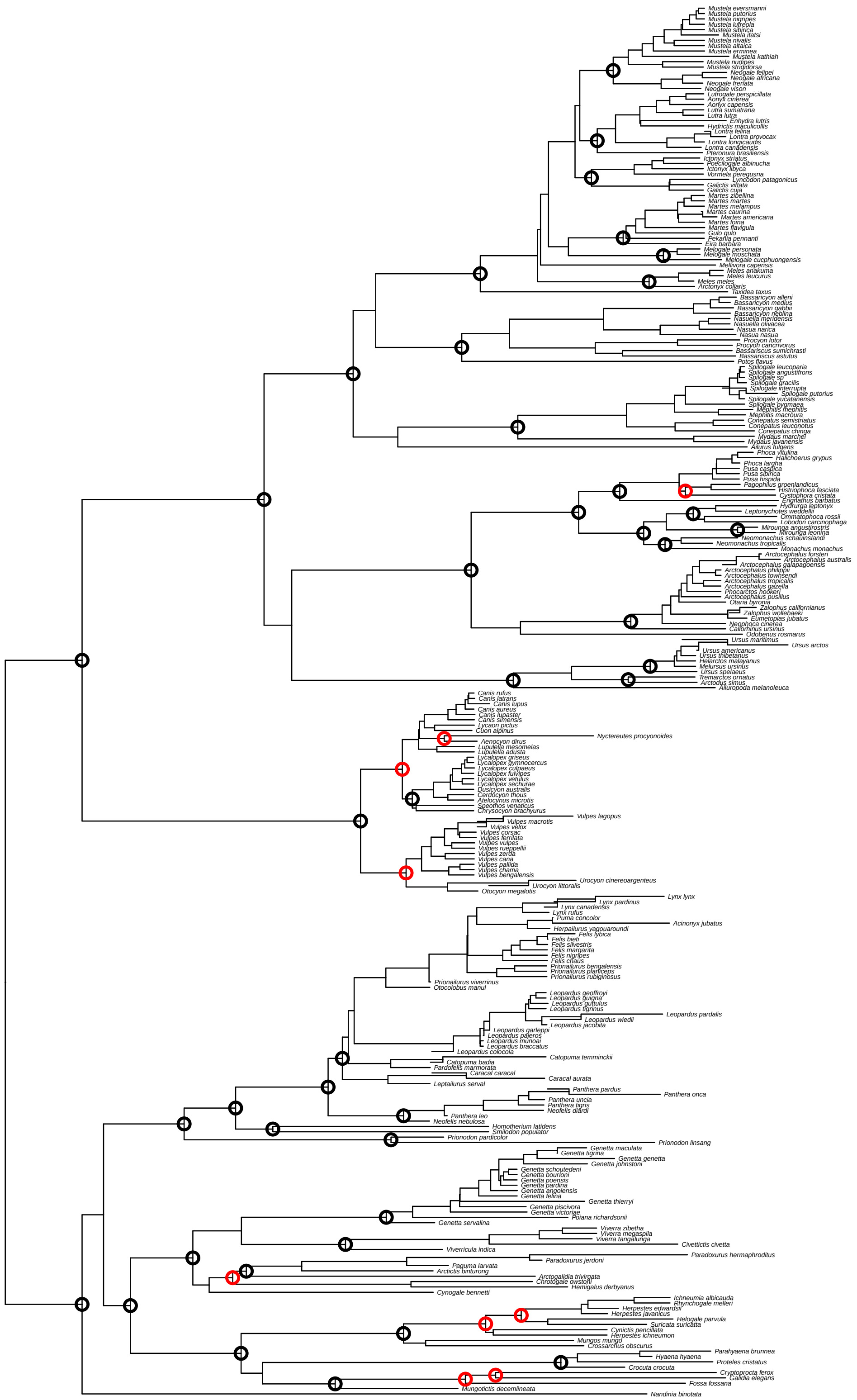

**Figure S27:** Neighbor-joining supertree of Carnivora inferred from an average distance matrix with imputed missing entries and with mtDNA-based source trees downweighted by a factor of 10. The circles indicate nodes congruent with (black) or contradicting (red) the 53 predefined benchmark clades. Note that the tree is not ultrametric, rendering the branch lengths uninterpretable.

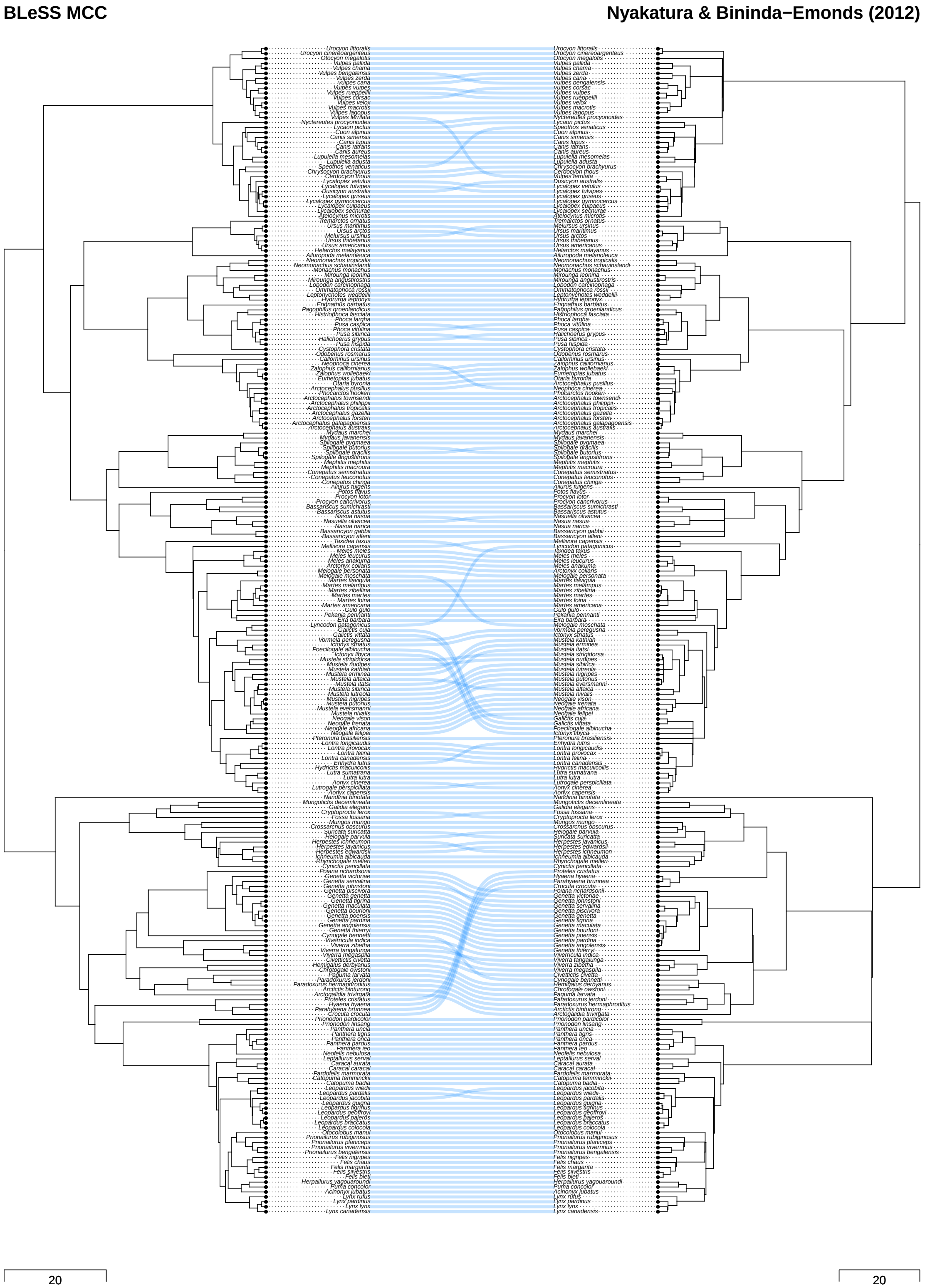

**Figure S28:** Rotation-optimized tanglegram illustrating the differences in topology and divergence times between the best BLeSS estimate of carnivoran phylogeny (see Table S14) and the time-calibrated supertree of Nyakatura and Bininda-Emonds (2012). The inter-tree edges connect corresponding tips, and their crossings indicate topological conflict. The scale bars denote branch durations in units of millions of years.

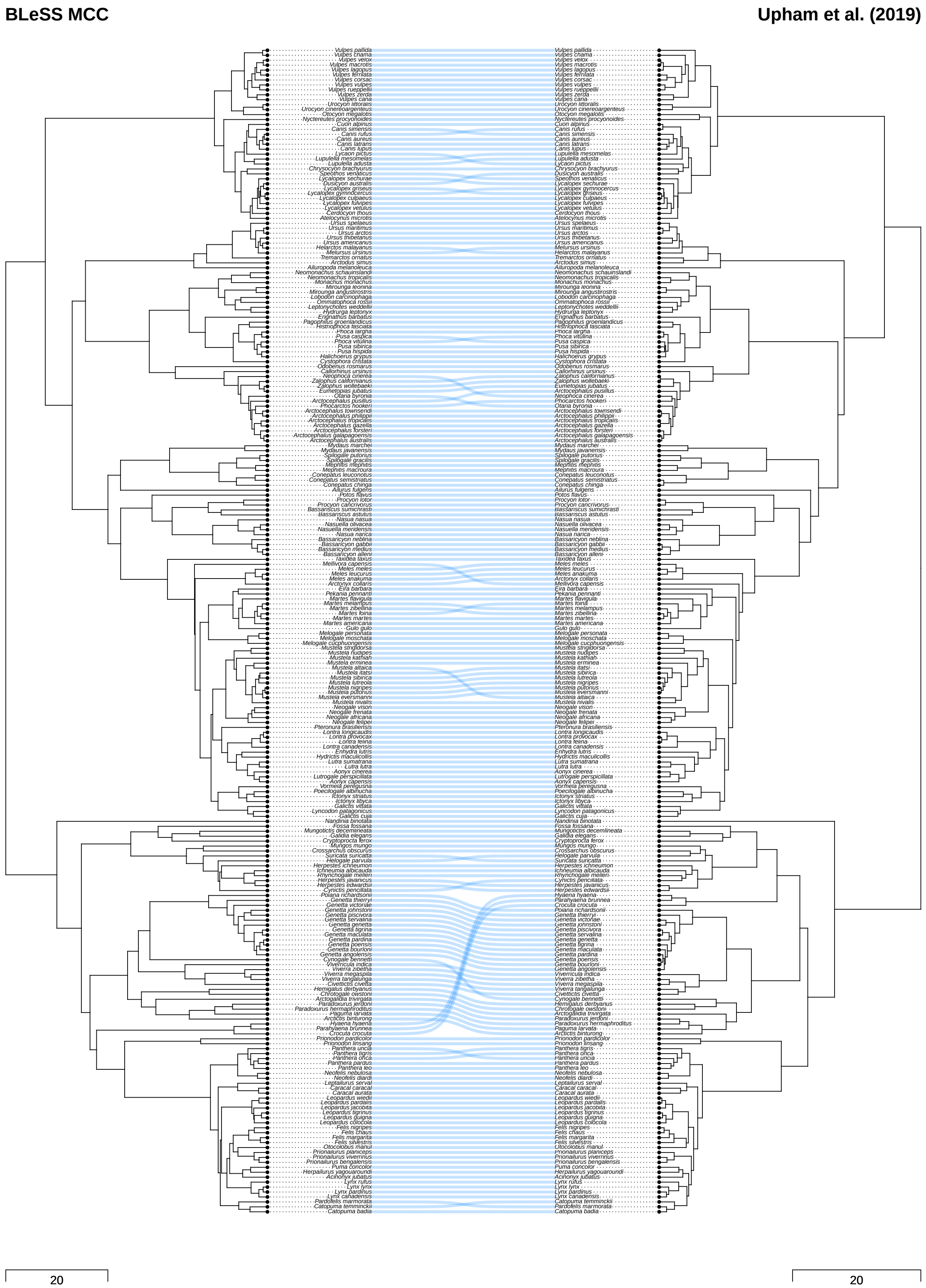

**Figure S29:** Rotation-optimized tanglegram illustrating the differences in topology and divergence times between the best BLeSS estimate of carnivoran phylogeny (see Table S14) and the backbone-and-patch time tree of Upham et al. (2019). The inter-tree edges connect corresponding tips, and their crossings indicate topological conflict. The scale bars denote branch durations in units of millions of years.

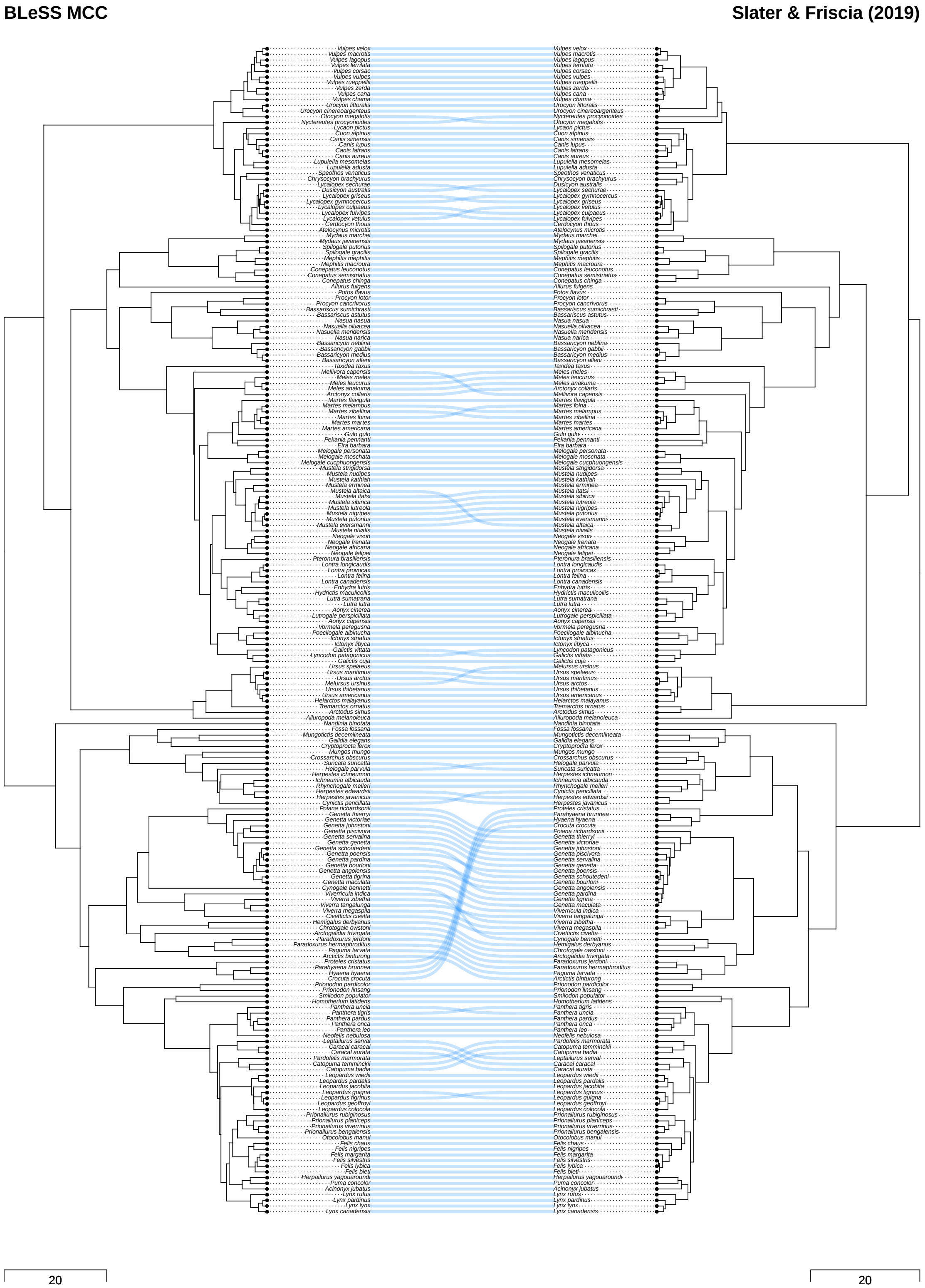

**Figure S30:** Rotation-optimized tanglegram illustrating the differences in topology and divergence times between the best BLeSS estimate of carnivoran phylogeny (see Table S14) and the supermatrix-based time tree of Slater and Friscia (2019). The inter-tree edges connect corresponding tips, and their crossings indicate topological conflict. The scale bars denote branch durations in units of millions of years.

### 3 Additional Figures

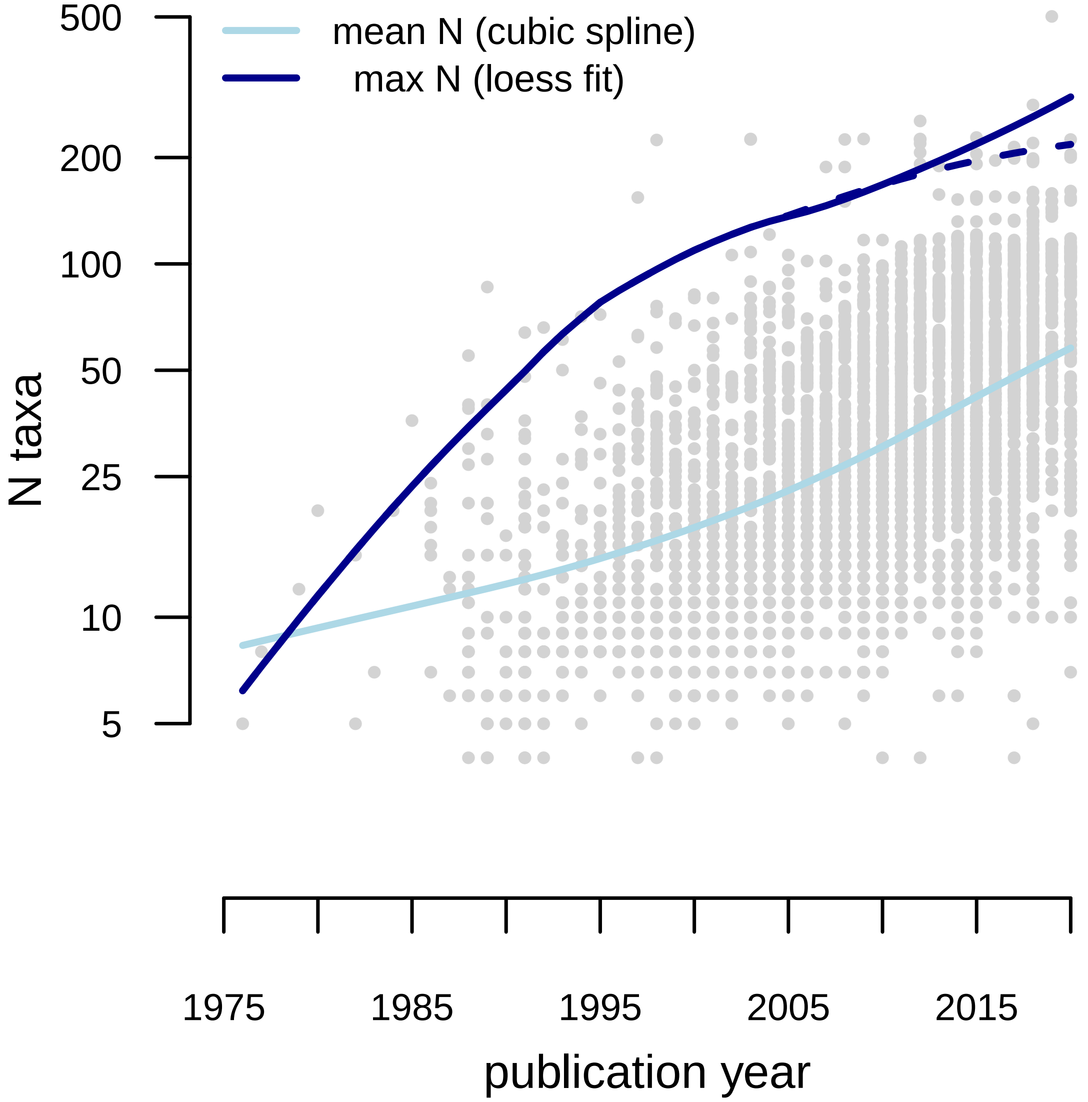

**Figure S31:** Number of taxa ($N$) included in morphological phylogenetic datasets over time, based on 3671 studies published between 1975 and 2020 (inclusive) and sourced from `https://graemetlloyd.com/matr.html`. A cubic spline (light blue line) fitted to $\log N$ shows that the average dataset only increased from 8.3 to 58 taxa during this period (note the log scale on the $y$–axis). A loess fit (dark blue solid line) shows that the maximum number of taxa increased as well, but a marked slow-down in the rate of increase becomes apparent after the removal of a single outlier (Hartman et al., 2019; dark blue dashed line), which contains twice as many taxa as any other study published during the analyzed period. Figure reprinted with permission from Lloyd and Slater (2020, Fig. 1), the preprint version of a study later published as Lloyd and Slater (2021).

Lindblad-Toh, K., Wade, C. M., Mikkelsen, T. S., Karlsson, E. K., Jaffe, D. B., Kamal, M., Clamp, M., Chang, J. L., Kulbokas, E. J., Zody, M. C., Mauceli, E., Xie, X., Breen, M., Wayne, R. K., Ostrander, E. A., Ponting, C. P., Galibert, F., Smith, D. R., deJong, P. J., Kirkness, E., Alvarez, P., Biagi, T., Brockman, W., Butler, J., Chin, C.-W., Cook, A., Cuff, J., Daly, M. J., DeCaprio, D., Gnerre, S., Grabherr, M., Kellis, M., Kleber, M., Bardeleben, C., Goodstadt, L., Heger, A., Hitte, C., Kim, L., Koepfli, K.-P., Parker, H. G., Pollinger, J. P., Searle, S. M. J., Sutter, N. B., Thomas, R., Webber, C., Baldwin, J., Abebe, A., Abouelleil, A., Aftuck, L., Ait-zahra, M., Aldredge, T., Allen, N., An, P., Anderson, S., Antoine, C., Arachchi, H., Aslam, A., Ayotte, L., Bachantsang, P., Barry, A., Bayul, T., Benamara, M., Berlin, A., Bessette, D., Blitshteyn, B., Bloom, T., Blye, J., Boguslavskiy, L., Bonnet, C., Boukhgalter, B., Brown, A., Cahill, P., Calixte, N., Camarata, J., Cheshatsang, Y., Chu, J., Citroen, M., Collymore, A., Cooke, P., Dawoe, T., Daza, R., Decktor, K., DeGray, S., Dhargay, N., Dooley, K., Dooley, K., Dorje, P., Dorjee, K., Dorris, L., Duffey, N., Dupes, A., Egbiremolen, O., Elong, R., Falk, J., Farina, A., Faro, S., Ferguson, D., Ferreira, P., Fisher, S., FitzGerald, M., Foley, K., Foley, C., Franke, A., Friedrich, D., Gage, D., Garber, M., Gearin, G., Giannoukos, G., Goode, T., Goyette, A., Graham, J., Grandbois, E., Gyaltsen, K., Hafez, N., Hagopian, D., Hagos, B., Hall, J., Healy, C., Hegarty, R., Honan, T., Horn, A., Houde, N., Hughes, L., Hunnicutt, L., Husby, M., Jester, B., Jones, C., Kamat, A., Kanga, B., Kells, C., Khazanovich, D., Kieu, A. C., Kisner, P., Kumar, M., Lance, K., Landers, T., Lara, M., Lee, W., Leger, J.-P., Lennon, N., Leuper, L., LeVine, S., Liu, J., Liu, X., Lokyitsang, Y., Lokyitsang, T., Lui, A., Macdonald, J., Major, J., Marabella, R., Maru, K., Matthews, C., McDonough, S., Mehta, T., Meldrim, J., Melnikov, A., Meneus, L., Mihalev, A., Mihova, T., Miller, K., Mittelman, R., Mlenga, V., Mulrain, L., Munson, G., Navidi, A., Naylor, J., Nguyen, T., Nguyen, N., Nguyen, C., Nguyen, T., Nicol, R., Norbu, N., Norbu, C., Novod, N., Nyima, T., Olandt, P., O'Neill, B., O'Neill, K., Osman, S., Oyono, L., Patti, C., Perrin, D., Phunkhang, P., Pierre, F., Priest, M., Rachupka, A., Raghuraman, S., Rameau, R., Ray, V., Raymond, C., Rege, F., Rise, C., Rogers, J., Rogov, P., Sahalie, J., Settipalli, S., Sharpe, T., Shea, T., Sheehan, M., Sherpa, N., Shi, J., Shih, D., Sloan, J., Smith, C., Sparrow, T., Stalker, J., Stange-Thomann, N., Stavropoulos, S., Stone, C., Stone, S., Sykes, S., Tchuinga, P., Tenzing, P., Tesfaye, S., Thoulutsang, D., Thoulutsang, Y., Topham, K., Topping, I., Tsamla, T., Vassiliev, H., Venkataraman, V., Vo, A., Wangchuk, T., Wangdi, T., Weiand, M., Wilkinson, J., Wilson, A., Yadav, S., Yang, S., Yang, X., Young, G., Yu, Q., Zainoun, J., Zembek, L., Zimmer, A., Lander, E. S., and members, B. S. P. (2005). Genome sequence, comparative analysis and haplotype structure of the domestic dog. *Nature*, 438(7069):803–819.

Lloyd, G. T. and Slater, G. J. (2020). A total-group phylogenetic metatree for Cetacea and the importance of fossil data in diversification analyses. *bioRχiv*. doi:10.1101/2020.06.24.169078.

Lloyd, G. T. and Slater, G. J. (2021). A total-group phylogenetic metatree for Cetacea and the importance of fossil data in diversification analyses. *Systematic Biology*, 70(5):922–939.

Lopes, F., Oliveira, L. R., Kessler, A., Beux, Y., Crespo, E., Cárdenas-Alayza, S., Majluf, P., Sepúlveda, M., Brownell Jr, R. L., Franco-Trecu, V., Páez-Rosas, D., Chaves, J., Loch, C., Robertson, B. C., Acevedo-Whitehouse, K., Elorriaga-Verplancken, F. R., Kirkman, S. P.,

Peart, C. R., Wolf, J. B. W., and Bonatto, S. L. (2021). Phylogenomic discordance in the eared seals is best explained by incomplete lineage sorting following explosive radiation in the Southern Hemisphere. *Systematic Biology*, 70(4):786–802.

Lynch, L. M. (2020). Fossil calibration of mitochondrial phylogenetic relationships of North American pine martens, *Martes*, suggests an older divergence of *M. americana* and *M. caurina* than previously hypothesized. *Journal of Mammalian Evolution*, 27(3):535–548.

McDonough, M. M., Ferguson, A. W., Dowler, R. C., Gompper, M. E., and Maldonado, J. E. (2022). Phylogenomic systematics of the spotted skunks (Carnivora, Mephitidae, *Spilogale*): Additional species diversity and Pleistocene climate change as a major driver of diversification. *Molecular Phylogenetics and Evolution*, 167:107266.

Meredith, R. W., Janečka, J. E., Gatesy, J., Ryder, O. A., Fisher, C. A., Teeling, E. C., Goodbla, A., Eizirik, E., Simão, T. L. L., Stadler, T., Rabosky, D. L., Honeycutt, R. L., Flynn, J. J., Ingram, C. M., Steiner, C., Williams, T. L., Robinson, T. J., Burk-Herrick, A., Westerman, M., Ayoub, N. A., Springer, M. S., and Murphy, W. J. (2011). Impacts of the Cretaceous Terrestrial Revolution and KPg extinction on mammal diversification. *Science*, 334(6055):521–524.

Mitra, S., Kunteepuram, V., Koepfli, K.-P., Mehra, N., Tabasum, W., Sreenivas, A., and Gaur, A. (2019). Characteristics of the complete mitochondrial genome of the monotypic genus *Arctictis* (Family: Viverridae) and its phylogenetic implications. *PeerJ*, 7:e8033.

Nascimento, F. O. D., Cheng, J., and Feijó, A. (2021). Taxonomic revision of the pampas cat *Leopardus colocola* complex (Carnivora: Felidae): an integrative approach. *Zoological Journal of the Linnean Society*, 191(2):575–611.

Nigenda-Morales, S. F., Gompper, M. E., Valenzuela-Galván, D., Lay, A. R., Kapheim, K. M., Hass, C., Booth-Binczik, S. D., Binczik, G. A., Hirsch, B. T., McColgin, M., Koprowski, J. L., McFadden, K., Wayne, R. K., and Koepfli, K.-P. (2019). Phylogeographic and diversification patterns of the white-nosed coati (*Nasua narica*): Evidence for south-to-north colonization of North America. *Molecular Phylogenetics and Evolution*, 131:149–163.

Nyakatura, K. and Bininda-Emonds, O. R. P. (2012). Updating the evolutionary history of Carnivora (Mammalia): a new species-level supertree complete with divergence time estimates. *BMC Biology*, 10:12.

Ogilvie, H. A., Mendes, F. K., Vaughan, T. G., Matzke, N. J., Stadler, T., Welch, D., and Drummond, A. J. (2022). Novel integrative modeling of molecules and morphology across evolutionary timescales. *Systematic Biology*, 71(1):208–220.

Paijmans, J. L. A., Barnett, R., Gilbert, M. T. P., Zepeda-Mendoza, M. L., Reumer, J. W. F., de Vos, J., Zazula, G., Nagel, D., Baryshnikov, G. F., Leonard, J. A., Rohland, N., Westbury, M. V., Barlow, A., and Hofreiter, M. (2017). Evolutionary history of saber-toothed cats based on ancient mitogenomics. *Current Biology*, 27(21):3330–3336.

Patou, M.-L., Debruyne, R., Jennings, A. P., Zubaid, A., Rovie-Ryan, J. J., and Veron, G. (2008). Phylogenetic relationships of the Asian palm civets (Hemigalinae & Paradoxurinae, Viverridae, Carnivora). *Molecular Phylogenetics and Evolution*, 47(3):883–892.

Patou, M.-L., Mclenachan, P. A., Morley, C. G., Couloux, A., Jennings, A. P., and Veron, G. (2009). Molecular phylogeny of the Herpestidae (Mammalia, Carnivora) with a special emphasis on the Asian *Herpestes*. *Molecular Phylogenetics and Evolution*, 53:69–80.

Patou, M.-L., Wilting, A., Gaubert, P., Esselstyn, J. A., Cruaud, C., Jennings, A. P., Fickel, J., and Veron, G. (2010). Evolutionary history of the *Paradoxurus* palm civets – a new model for Asian biogeography. *Journal of Biogeography*, 37(11):2077–2097.

Perri, A. R., Mitchell, K. J., Mouton, A., Álvarez-Carretero, S., Hulme-Beaman, A., Haile, J., Jamieson, A., Meachen, J., Lin, A. T., Schubert, B. W., Ameen, C., Antipina, E. E., Bover, P., Brace, S., Carmagnini, A., Carøe, C., Samaniego Castruita, J. A., Chatters, J. C., Dobney, K., dos Reis, M., Evin, A., Gaubert, P., Gopalakrishnan, S., Gower, G., Heiniger, H., Helgen, K. M., Kapp, J., Kosintsev, P. A., Linderholm, A., Ozga, A. T., Presslee, S., Salis, A. T., Saremi, N. F., Shew, C., Skerry, K., Taranenko, D. E., Thompson, M., Sablin, M. V., Kuzmin, Y. V., Collins, M. J., Sinding, M.-H. S., Gilbert, M. T. P., Stone, A. C., Shapiro, B., Van Valkenburgh, B., Wayne, R. K., Larson, G., Cooper, A., and Frantz, L. A. F. (2021). Dire wolves were the last of an ancient New World canid lineage. *Nature*, 591(7848):87–91.

Püschel, H. P., O'Reilly, J. E., Pisani, D., and Donoghue, P. C. J. (2020). The impact of fossil stratigraphic ranges on tip-calibration, and the accuracy and precision of divergence time estimates. *Palaeontology*, 63(1):67–83.

Rao, H., Yang, Y., Liu, J., Westbury, M. V., Zhang, C., and Shao, Q. (2020). Palaeoproteomic analysis of Pleistocene cave hyenas from east Asia. *Scientific Reports*, 10(1):16674.

Scheel, D.-M., Slater, G. J., Kolokotronis, S.-O., Potter, C. W., Rotstein, D. S., Tsangaras, K., Greenwood, A. D., and Helgen, K. M. (2014). Biogeography and taxonomy of extinct and endangered monk seals illuminated by ancient DNA and skull morphology. *ZooKeys*, 409:1–33.

Slater, G. J. and Friscia, A. R. (2019). Hierarchy in adaptive radiation: a case study using the Carnivora (Mammalia). *Evolution*, 73(3):524–539.

Trindade, F. J., Rodrigues, M. R., Figueiró, H. V., Li, G., Murphy, W. J., and Eizirik, E. (2021). Genome-wide SNPs clarify a complex radiation and support recognition of an additional cat species. *Molecular Biology and Evolution*, 38(11):4987–4991.

Tseng, Z. J., Wang, X., Slater, G. J., Takeuchi, G. T., Li, Q., Liu, J., and Xie, G. (2014). Himalayan fossils of the oldest known pantherine establish ancient origin of big cats. *Proceedings of the Royal Society B: Biological Sciences*, 281(1774):20132686.

Upham, N. S., Esselstyn, J. A., and Jetz, W. (2019). Inferring the mammal tree: Species-level sets of phylogenies for questions in ecology, evolution, and conservation. *PLoS Biology*, 17(12):e3000494.

Westbury, M. V., Le Duc, D., Duchêne, D. A., Krishnan, A., Prost, S., Rutschmann, S., Grau, J. H., Dalén, L., Weyrich, A., Norén, K., Werdelin, L., Dalerum, F., Schöneberg, T., and Hofreiter, M. (2021). Ecological specialization and evolutionary reticulation in extant Hyaenidae. *Molecular Biology and Evolution*, 38(9):3884–3897.

Wolsan, M. and Sato, J. J. (2020). Parallel loss of sweet and umami taste receptor function from phocids and otarioids suggests multiple colonizations of the marine realm by pinnipeds. *Journal of Biogeography*, 47(1):235–249.
